# Iterative community-driven development of a SARS-CoV-2 tissue simulator

**DOI:** 10.1101/2020.04.02.019075

**Authors:** Michael Getz, Yafei Wang, Gary An, Maansi Asthana, Andrew Becker, Chase Cockrell, Nicholson Collier, Morgan Craig, Courtney L. Davis, James R. Faeder, Ashlee N. Ford Versypt, Tarunendu Mapder, Juliano F. Gianlupi, James A. Glazier, Sara Hamis, Randy Heiland, Thomas Hillen, Dennis Hou, Mohammad Aminul Islam, Adrianne L. Jenner, Furkan Kurtoglu, Caroline I. Larkin, Bing Liu, Fiona Macfarlane, Pablo Maygrundter, Penelope A Morel, Aarthi Narayanan, Jonathan Ozik, Elsje Pienaar, Padmini Rangamani, Ali Sinan Saglam, Jason Edward Shoemaker, Amber M. Smith, Jordan J.A. Weaver, Paul Macklin

## Abstract

The 2019 novel coronavirus, SARS-CoV-2, is a pathogen of critical significance to international public health. Knowledge of the interplay between molecular-scale virus-receptor interactions, single-cell viral replication, intracellular-scale viral transport, and emergent tissue-scale viral propagation is limited. Moreover, little is known about immune system-virus-tissue interactions and how these can result in low-level (asymptomatic) infections in some cases and acute respiratory distress syndrome (ARDS) in others, particularly with respect to presentation in different age groups or pre-existing inflammatory risk factors. Given the nonlinear interactions within and among each of these processes, multiscale simulation models can shed light on the emergent dynamics that lead to divergent outcomes, identify actionable “choke points” for pharmacologic interventions, screen potential therapies, and identify potential biomarkers that differentiate patient outcomes. Given the complexity of the problem and the acute need for an actionable model to guide therapy discovery and optimization, we introduce and iteratively refine a prototype of a multiscale model of SARS-CoV-2 dynamics in lung tissue. The first prototype model was built and shared internationally as open source code and an online interactive model in under 12 hours, and community domain expertise is driving regular refinements. In a sustained community effort, this consortium is integrating data and expertise across virology, immunology, mathematical biology, quantitative systems physiology, cloud and high performance computing, and other domains to accelerate our response to this critical threat to international health. More broadly, this effort is creating a reusable, modular framework for studying viral replication and immune response in tissues, which can also potentially be adapted to related problems in immunology and immunotherapy.

## Introduction

The ongoing pandemic caused by the novel severe acute respiratory syndrome coronavirus 2 (SARS-CoV-2) has illuminated the global public health threat posed by highly pathogenic coronaviruses that emerge from zoonotic sources. With the majority of the world’s population immunologically naïve and no available antivirals or vaccines, over 233,000,000 infections and 4,500,000 deaths amassed worldwide by the end of September 2021^1^. Coronavirus disease 2019 (COVID-19)—caused by SARS-CoV-2 infection—is characterized by a range of respiratory symptoms, including fever and cough^2,3^, that can progress to acute respiratory distress syndrome (ARDS) in some patients^4,5^. Age and comorbidities appear to be the main risk factors for development of severe disease^6–8^. However, the dynamics of virus replication, interaction with host immune responses, and spread within the respiratory tract are still being established. Despite the fact that several vaccines are now available, these are not yet widely distributed and there remains a critical need to further understand the infection in order to quickly identify pharmacologic interventions and optimal therapeutic designs that work to lessen virus replication and disease severity. In addition, the emergence of break-through infections among vaccinated individuals highlights the importance of understanding the longevity and effectiveness of immunity, induced by vaccination or prior infection. However, this requires an international community effort that integrates expertise across a variety of domains and a platform that can be iteratively updated as new information and data arises.

To aid this effort, we have assembled an international, multi-disciplinary coalition to rapidly develop an open source, multi-scale tissue simulator that can be used to investigate mechanisms of intracellular viral replication, infection of epithelial cells, host immune response, and tissue damage. The aim of this project is to concentrate community modeling efforts to create a comprehensive multiscale simulation framework that can subsequently be calibrated, validated, and used to rapidly explore and optimize therapeutic interventions for COVID-19. Once the prototype has been completed (after several design iterations), this coalition will transition to maintain and support the simulation framework and aggregate calibrated/validated parameter values.

To address the acute need for rapid access to an actionable model, we are using a community-driven coalition and best open science practices to build and iteratively refine the model:

1. **Modular and reusable:** The model is built from mechanistic “first principles” based on domain expertise in immunology, virology, and tissue biology. Separate generalized modules for key processes (virus-receptor binding and endocytosis, replication, interferon responses, cell death, and local and systemic responses) ensure that the model can be adapted and calibrated to new viruses and new treatments without the need for relearning model rules from fresh data.
2. **Open source and GitHub:** All simulation source code is shared as open source on GitHub, with well-defined, versioned, and documented releases, and Zenodo-generated archives and DOIs.
3. **Interactive cloud-hosted models:** Every prototype version is rapidly transformed into a cloud-hosted, interactive model to permit faster scientific communication across communities, particularly with virologists, immunologists, pharmacologists, and others who have essential insights but ordinarily would not directly run the simulation models.
4. **Social media and virtual feedback:** We enlist community participation (feedback, modeling contributions, software contributions, and data contributions) through social media, virtual seminars, web forms, a dedicated Slack workspace, and weekly team meetings. We particularly encouraging feedback and data contributions by domain experts in virology, epidemiology, immunology, and mathematical biology.
5. **Frequent preprint updates:** Each model iteration is accompanied by a cloud-hosted, interactive app (see #3) and an updated preprint on *bioRxiv*.
6. **Integration of feedback:** All community feedback is evaluated to plan the next set of model refinements and recorded in an updated *bioRxiv* preprint.

Our first test of this workflow saw a first proof-of-concept software release (Steps 2-3) in 12 hours, and the first integration of community feedback and preprint dissemination was complete within a week. We have begun integrating community feedback, and it is our intention to continue iterate refinement.

### Goals and guiding principles

This project is community-driven, including the following contributions:

1. **Community priorities:** The community helps define the driving research questions, set the project scope, and select the critical biological components to be modeled.
2. **Consensus hypotheses:** The community drives a shared, clearly-written consensus specification of the underlying biological hypotheses.
3. **Mathematical modeling:** The community helps develop, review, and refine the mathematical interpretation of the biological hypotheses.
4. **Computational implementation:** The computational implementation is shared as open source with community definition of specifications, unit tests, coding, and code review.
5. **Community feedback:** Community feedback on the model realism, hypotheses, mathematics, computational implementation, and techniques is encouraged throughout the development process.
6. **Community parameters and data:** Community contributions of parameter estimates and data contributions are aggregated to assist in model development and constraint.

#### Project scope

While by definition the project scope can be refined by the community, the initial project scope is to:

1. Develop the general computational framework sufficiently to address many of the community-driven research questions.
2. Deliver a working simulation framework for use by others to perform calibration and validation. That is, the prototyping aims of this project are complete once the model is capable of demonstrating essential biological behaviors qualitatively.
3. To provide a software framework whose underlying hypotheses, mathematics, and computational implementation have been rigorously assessed by appropriate domain experts.

In particular, while this project will work to constrain, estimate, and calibrate parameters to the greatest extent possible, it is *not* within scope to delay software release until full calibration and validation. Those tasks are within scope of fully funded teams with dedicated experiments.

This project aims to deliver software that one can reasonably expect to calibrate and validate, thus freeing funded investigations from expensive early software development while providing a broad community consensus on key biological hypotheses. By rapidly prototyping this software, we aim to accelerate many funded research efforts.

#### Essential model components

As part of defining the project scope, we have identified the following critical model components:

1. Virus dissemination in epithelial tissue
2. Virus binding, endocytosis, replication, and exocytosis
3. Infected cell responses, including changes to metabolism, secreted signals, and death
4. Inflammatory response
5. Ramp up of the immune response (particularly in lymph nodes)
6. Immune cell infiltration
7. Immune cell predation of infected and other cells
8. Tissue damage by death of cells due to infection or host response

#### Guiding principles

The coalition aims to model not merely the disease endpoints, but the disease *dynamics.* This will allow scientists to investigate mechanistic “what if” questions on potential interventions: *What if* we could inhibit endocytosis? *What if* we could introduce a cytokine early or late in the disease course? *What if* the infected cell apoptosis could be accelerated or delayed?

To accomplish this, we use a modular design: an overall tissue-scale model integrates an array of targeted *submodels* that simulate critical processes (e.g., receptor binding and trafficking and virus replication). Each submodel is clearly specified to enable interoperability and to make it feasible for subteams to simultaneously develop and test the model components in parallel. Throughout development, we use open source methodologies that enhance communication, transparency, and reproducibility. See Box 1.

##### Box 1: Guiding principles for the rapid prototyping a modular, multiscale model.

###### Guiding principles: motivation

- The model should investigate the dynamics of infection and treatment, not merely endpoints.
- The model should help the community ask “what if” questions to guide experiments and interventions^9,10^.

###### Guiding principles: approach

- Community consensus will be gathered and efforts pooled into a “standardized” model that captures key SARS-CoV-2 dynamics. The model will be supplied to the community for parallel studies by multiple labs.
- The model framework will be built with relatively sparse data, relying upon domain expertise and observations to choose its general form and assess its qualitative behavior.
- The model will be modular. Each submodel will have well-defined inputs and outputs, allowing parallel development and replacement of submodels with improved versions without change to the overall integrated model or other submodels.
- As part of the model formulation, documentation, and dissemination, we will craft clearly delineated “conceptual model and hypotheses” to encourage development of independent models with distinct methodologies and software frameworks.
- The submodels will be independently executable and verifiable, allowing parallel development.
- The overall model framework will be released periodically released numbered versions (distributions) that bundle the best working version of each submodel as it exists at the time of release, allowing end-users (the community) to use well-defined, well-tested snapshots of the project.
- The model (and known parameter values) will be made publicly available as open source for maximum public benefit.
- The model will be made publicly available as an interactive web app to encourage community participation, to accelerate scientific dissemination, and to increase public educational benefit.
- Rapid prototyping will be used to encourage an iterative develop-test-refine cycle, build expertise, and gain community feedback.
- Data and parameter sharing is encouraged throughout this effort for use following the model’s “completion.”
- The model will be developed to a point that it has correct qualitative behavior so that calibration is likely to succeed. This is the “product” for use in subsequent investigations by multiple teams. See the ***scoping*** statements above.
- After the model prototyping is complete (the goal of this paper), a maintenance and support phase will be entered to fix bugs, support scientist users, and add features identified by the user community.

#### Critical questions for the model framework

The community identified a series of driving biological questions concerning COVID-19 to guide the development of the model framework (see **Box 2**). It is expected that the model will not initially be able to address all the questions listed; rather, the development plan envisions that with each iteration of the model framework it will expand in its representational capacity as directed by the topics listed in Box 2. Furthermore, as with all modeling projects, we anticipate that as the framework develops it will generate new questions and/or be responsive to the rapidly evolving knowledge-base concerning COVID-19.

##### Box 2: Community-selected scientific questions driving the model’s development.

1. What are the critical “choke points” in viral infection, replication, and propagation?
2. Which interventions could most effectively leverage identified vulnerabilities in viral replication?
3. What unanticipated dynamics can emerge from a single molecular-scale inhibition?
4. Does the initial level of exposure to the virus affect the severity of the progression of the disease and how could this be ameliorated?
5. What are the key points of virus-immune interactions that drive mild versus severe (e.g., ARDS) responses?
6. What are key differences at the target cell level during innate versus adaptive immune responses?
7. Are there threshold levels of infection at the cellular or tissue level that indicate a switch from asymptomatic to symptomatic or from mild to severe disease in a patient?
8. Through what mechanisms do certain patient characteristics, pre-existing conditions, or background medications increase the likelihood of adverse outcomes?
9. What interventions could accelerate immunity?
10. What interventions can reduce or reverse adverse immune reactions?
11. At what stage is an intervention most beneficial?
12. How does viral mutagenicity affect the robustness of a therapy or a therapeutic protocol?
13. How does cellular heterogeneity affect infection dynamics?
14. How does the nearby tissue environment, such as the mucus layer, affect infection dynamics?
15. How does the infection spread from its initial locus of infection to other tissues (in particular, from upper respiratory tract to the bronchi, bronchioles, and alveoli within lungs)? How does stochasticity impact these dynamics?
16. How do tissues recover after clearance of local infection? Can scarring be minimized to reduce long-term adverse effects in recovered patients?
17. How do adverse effects in SARS-CoV-2 infected epithelia differ (mechanistically) from other infections and other modes of epithelial dysfunction?

### Key biology for the simulation model

This rapid prototyping effort brings together specialists from a broad variety of domains: virology and infectious diseases, mathematical biology, computer science, high performance computing, data science, and other disciplines. Therefore, it is critical that all members of the project have access to a clear description of underlying biology that drive the model’s assumptions. In this section we outline key aspects of viral replication and host response in functional terms needed for development of agent-based, multi-scale and multi-physics models.

#### Cell infection and viral replication

The key cell-level process is viral infection of a single cell, followed by replication to create new virions:

1. SARS-CoV-2 is a single-stranded enveloped RNA virus^11^. A virion (single virus particle) has a lipid coating (envelope) that protects the virus when outside a cell (or host). Each virus has dozens of spike glycoproteins that bind to ACE2 (receptors) on select cell membranes^3,11^.
2. Virions travel in the tissue microenvironment to reach a cell membrane. The spike binds to an available ACE2 receptor on the cell membrane. Prior to binding to the ACE2 receptor, the spike is cleaved by the protease, TMPRSS2, which is required for efficient cell entry^12^. Multiple modes of transport (e.g., passive diffusion in fluids and porous tissues, mucociliary clearance, chemotaxis, ultrafiltration driven by hydrostatic and oncotic pressure through permeable cell junctions, and cellular active transport) may play a role at slow and fast time scales. There may also be surface contact transmission between neighboring cells.
3. The cell internalizes the adhered virus via endocytosis into a vesicle.
4. The endocytosed virion—now residing in a vesicle with lowered pH—is uncoated to release its mRNA contents into the cell cytoplasm.
5. Copying viral RNA creates a (-) RNA template, which is used for (+) RNA production.
6. RNA is used to synthesize viral RNA and proteins.
7. Viral proteins are transported to the interior surface of the cell membrane.
8. Viral proteins at the cell membrane are assembled into virions.
9. Assembled virions are exported from the cell by exocytosis.
10. When a cell dies and lyses, some or all partly and fully assembled virions can be released into the tissue microenvironment.

Once infected, an individual cell cannot “recover” (e.g., by actively degrading viral RNA and stopping endocytosis) to return to normal function. Rather, the cell is irreversibly committed to eventual death. For further detail, see review articles on RNA virus replication dynamics^13,14^.

#### Infected cell responses

Although infected cells (e.g., type 1 or type 2 alveolar cells in the lung) cannot recover, their response can slow viral replication and reduce infection of nearby cells. Infected cells do this by secreting type I interferons (IFN-α,β), which diffuse and bind to receptors on nearby cells to reduce viral replication following infection, activate an inflammatory response, and induce gene transcription^15^ to minimize cycling and/or induce apoptosis in these cells^16^. Secreted IFN-α,β are important activators and regulators of the innate and adaptive immune responses^16^. Many respiratory viruses, including influenza and SARS-CoVs^17^, have evolved to inhibit IFN activation^18^, and evidence is emerging that certain non-structural proteins produced by SARS-CoV-2 infected cells interfere with IFN-*α*, *β* and chemokines by inhibiting production and suppressing signaling^17,18^.

Eventually, infected cells die (by apoptosis, necroptosis, or pyroptosis^19^), lyse, and release unassembled viral components^19^. While the mechanism of cell death in SARS-CoV-2 is currently unknown, in other RNA virus infections, cells can undergo apoptotic, necrotic, or pyroptotic death over the course of viral infection^20^. Disruption of cell metabolism and competition for critical substrates may also contribute to cell death^21,22^.

#### Inflammatory and immune responses

Lethal SARS and MERS in humans has been correlated with elevated IFN-*α*,*β* ^23^, myeloid activity, and impaired T and B cells^24,25^, with the timing of IFN-*α*,*β* critical^26,27^. IFN-*α*,*β*s secreted by infected cells or by immune cells diffuse to surrounding cells and recruit innate immune cells, such as macrophages and neutrophils, to the area. Recent studies comparing SARS-CoV-2 with SARS-CoV have revealed that SARS-CoV-2 replicates more efficiently in pneumocytes and alveolar macrophages, but IFN-*α*,*β* secretion is blunted in SARS-CoV-2 infected cells^28^. In COVID-19 patients, decreased numbers of T cells, natural killer (NK) cells, and, to a lesser extent, B cells occur, and the extent of depletion of T cells has been correlated with disease severity^2,3,29^. Rapid inhibition of viral replication requires early and high levels of IFN-*α*,*β* secretion and activation^30^. The production of excess inflammatory cytokines, such as IL-1, IL-6 and TNF-*α* and other chemokines by infected cells results in increased macrophage and neutrophil presence, which correlates with lung dysfunction^31,32^. Delayed IFN-α,β production also promotes inflammatory macrophage recruitment that contributes to vascular leakage and impaired T cell function^26,27^. Activated macrophages also produce other proinflammatory cytokines like IL-1, IL-6, and TNF-α, among others, that enhance infiltration of immune cells and interact with endothelial cells to cause vasodilation^33^. The excess production of IL-1 and IL-6 may be related to several viral proteins shown to directly activate the inflammasome pathway, the innate immune response responsible for IL-1β secretion^34–36^. Moreover, epithelial tissue death can reduce tissue integrity, contributing to further immune infiltration, fluid leakage and edema, and acute respiratory distress^37–39^.

In severe cases, a “cytokine storm” of pro-inflammatory cytokines (e.g., IL-2, IL-7, IL-10, G-CSF, IP-10, MCP-1, MIP-1A, and TNF-α) induces extensive tissue damage^31^. During influenza virus infection, there is some evidence that ARDS is correlated with the extent of infection in the lower respiratory tract and increased cytokine activity resulting from exposure of the endothelium^40^. Increases in neutrophil counts and the neutrophil-to-lymphocyte ratio (NLR) have been observed in patients with severe COVID-19^41^. The NLR has also been shown to be an important clinical predictor of disease severity^42^, as it reflects the innate to adaptive immune response. Neutrophils generally produce reactive oxygen species ROS, which can induce the death of infected and healthy cells in the local environment, further contributing to tissue damage^37^.

Coronaviruses have been shown to evade and modulate various host immune responses^43–45^. In addition to those discussed above, some evidence suggests that an antibody to spike protein enhances disease during SARS-CoV infection by inducing macrophage switching from a wound healing phenotype to an inflammatory phenotype^46^. Furthermore, influenza viruses and other SARS-CoVs are known to infect macrophages and T cells^3,47^, raising the possibility for SARS-CoV-2 to similarly infect these cell types. However, it has not yet been shown that SARS-CoV-2 infection of these cells is productive or simply alters their cytokine expression^31^. However, the ACE2 receptor has been linked to acute lung injury for both influenza and SARS-CoV viruses^48,49^.

#### Links between inflammation and poor clinical outcomes

Death following severe COVID-19 is often correlated with pre-existing medical conditions such as diabetes, hypertension, cardiac disease and obesity^6,50,51^. While the primary target of SARS-CoV-2 infection is clearly the respiratory system, several secondary factors play a role in the clinical outcome for a given patient. The complex interactions of viral infection, cytokine production, immune response, and pre-existing diseases is an active field of current research. Although the underlying risk factors for an individual to develop ARDS in response to SARS-CoV-2 infection have not yet been elucidated, it appears that a dysregulated immune response is central to this aspect of the disease^2,3,29,52^. In particular, chemokines are released following viral infection, which leads to the invasion of neutrophils and macrophages and release of ROS. IL-6 levels have been associated with more severe disease as patients who required ventilation exhibit increased systemic IL-6 levels, as reported by studies from Germany and China^53–55^. In addition, replication in the lower airways and exposure of endothelial cells may further amplify the inflammatory response^40^. Collectively, this leads to extensive tissue damage and depletion of epithelial cells, which may be connected to lethality^56^. Within the alveolar tissue and systemically, the feedback between viral load, adaptive and innate immune responses, and tissue damage is clearly a complex system. By utilizing a multi-scale framework to implement these interactions, we aim to connect circulating biomarkers, putative treatments, and clinically observed disease progression to pathophysiological changes at the cell and tissue level.

### Anticipated data to drive development and validation

It is important that model development considers the types of measurements and biological observations that will be available for later model constraint, calibration, and validation. As participation by the virology and pharmacology communities broadens, we anticipate that this list will grow. While we will endeavor to constrain and validate submodels of the model framework independently, we anticipate human clinical data to not fully determine parameters of the model. To address this concern, we will apply a “virtual population” approach and sensitivity analysis to explore model variability within clinically relevant bounds^57,58^. We anticipate the following data:

#### Organoid data for viral replication and targeted inhibition

Aarthi Narayanan’s virology lab is optimizing SARS-CoV-2 cultures in organoid model systems. These 3D model systems infect epithelial cells co-cultured with fibroblasts and endothelial cells and track the viral replication kinetics under control conditions and after treatment by inhibitors. These experiments measure (at various time points) infectious viral titers and genomic copy numbers in supernatants (outside the cells), viral genomic copy numbers in the infected cellss host cell death and inflammatory responses, and ATP and mitochondrial disruptions. See Appendix 2 for further detail.

#### Inflammation, ACE2 binding, and host response data

The international focus on SARS-CoV-2 is generating an unprecedented amount of mechanistic clinical and preclinical data. Randomized controlled interventional trials in general or specific populations will be of particular value to test and refine the model. As of September 30 2021, there were 3,326 trials registered at clinicaltrials.gov under the search terms “COVID-19+Drug”. Within these 3,326 trials, there are multiple interventions at different points of the pathophysiology, including, but not limited to: broad acting antivirals (e.g., remdesivir), hyperimmune plasma, IL-6 antibody (e.g., tocilizumab), protease inhibitors (e.g., lopinavir/ritonavir), chloroquine/hydroxychloroquine, and Janus kinase inhibitors (e.g., baricitinib). As this platform develops, we anticipate predicting the responses to such therapies and refining the model with emerging data such that the range of clinical responses are captured with adequate fidelity. In addition, data collected from patients or animals during infection, including the presence of various immune cell subsets in lung tissue and systemic markers of inflammation, will serve to differentiate responses to SARS-CoV-2. These data will be similarly integrated to calibrate and validate the model to ensure accurate predictions of therapeutic outcomes based on clinical characteristics. The development of effective vaccines also leads to further questions concerning efficacy and the longevity of the induced immunity. As of September 20, 2021 there were 976 trials registered at clinicaltrials.gov under the search terms “COVID-19+vaccine”. These are testing the effectiveness of new vaccines, booster immunizations and whether existing vaccines can be mixed and matched. These new studies can be integrated to answer questions concerning second infections or break-through infections in vaccinated individuals.

### Relevant prior modeling

Spurred initially by the emergence of HIV and relevant to the present SARS-CoV-2 pandemic, the field of viral dynamics modeling has been instrumental for understanding the evolution of host-virus interactions^59–67^, predicting treatment responses^68–72^, and designing novel and more effective therapeutic approaches^73–75^. The classic within-host mathematical model of viral infection uses a system of ordinary differential equations (ODEs) to describe the dynamics between uninfected epithelial (“target”) cells, infected cells in the eclipse phase, infected cells producing virus, and infectious virus^76^. This basic model has been shown to capture dynamics of both acute and chronic infection^77^, and has been extended to also include multiple viral (potentially resistant) strains^73^ and various aspects of host immune responses^78,79^. While such cell population-level ODE models generally do not account for single-cell effects, they are effective for detailing viral load, host immune response, and pathology dynamics^80–85^. Moreover, they can often be used to constrain and estimate parameters for more detailed models, develop novel hypotheses, and design confirmatory experiments^86,87^.

Some have modeled intracellular virus replication, including detailed models used for understanding replication and intervention points^58,88^, typically using systems of ODEs^89,90^. These models often include virus-receptor binding, receptor trafficking, endocytosis, viral uncoating, RNA transcription, protein synthesis, viral assembly, and viral exocytosis. However, to date no such model has been integrated with detailed spatiotemporal models of viral propagation in 3D tissues with dynamical models of immune interactions.

Agent-based models have been used to simulate viral propagation in 2D tissues with simplified models of viral replication in individual cells, particularly in the field of influenza virus infection^91–95^, spatial patterns from singlecell infections for a variety of other viral infections^96–98^ such as SARS^99^, and oncolytic viral therapies^100–103^. These models have generally not included detailed intracellular models of viral replication and individual cell responses to infection. However, they demonstrate the potential for including detailed intracellular models of viral replication in 2D and 3D tissues with the milieu of immune and epithelial cell types expected in actual patients, while also offering the opportunity to test hypotheses on the impact of viral mutagenicity and host cell heterogeneity on disease progression.

Agent-based models have also been used extensively to characterize the inflammatory dysfunction that produces the most severe manifestations of COVID19. The dynamics of forward feedback inflammation-induced tissue damage was examined with an early agent-based model of systemic inflammation^104^; this model was further developed into the Innate Immune Response ABM (IIRABM), which was used to perform in silico trials of anti-mediator/cytokine interventions (not too different from the types being tried for COVID19)^105^. More recently, the IIRABM has been used as a test platform for the application of genetic algorithms^106^ and model-based deep reinforcement learning^107^ to discover multi-modal and potentially adaptive mediator-directed therapies for acute systemic inflammation; this work is particularly relevant given the attempts to use anti-cytokine biologics during the current COVID19 pandemic. Finally, the IIRABM, as an endothelial-based model, was integrated with models of epithelial dysfunction to simulate individual and multiple organ dysfunction and failure seen with systemic hyper-inflammation^108^. As noted above, there are significant differences between the pathophysiology of bacterial sepsis and that of severe viral infections, but it appears that at some level of tissue damage the dynamics of the innate system come to fore in terms of the clinical significance.

The rapid prototyping approach of this coalition will use a performance-driven agent-based modeling platform^109^ to combine detailed intracellular models of viral endocytosis, replication, and exocytosis, disruption of cell processes (e.g., metabolism and compromised membranes) that culminate in cell death, inflammation signaling and immune responses, tissue damage, and other key effects outlined above in a comprehensive, open source simulation platform. We will deploy and refine interactive, web-hosted versions of the model to critical contributions by virologists, infectious disease modelers, and other domain experts. We will frequently update preprints to foster the fastest possible scientific dialog to iteratively refine this community resource.

#### Related modeling efforts and other future data sources

We are coordinating with related modeling efforts by a number of groups, such as early pilot work by David Odde and colleagues at the University of Minnesota, and early simulation work in Chaste^110,111^ (James Osborne and colleagues), Morpheus^112^ (Andreas Deutsch and colleagues), CompuCell3D^113^, and Biocellion^114^ (Ilya Shmulevich and co-workers). Thomas Hillen has hosted a COVID-19 Physiology Reading Group^115^ to exchange information and progress. Andrew Becker, Gary An, and Chase Cockrell recently adapter their prior multiscale agentbased modeling framework (CIABM) to simulate immune responses to viral respiratory infections, with a focus on SARS-CoV-2^116^. We are in regular contact with these communities to share data and biological hypotheses and to seek feedback, parameter insights, and data and code contributions.

The COVID-19 Cell Atlas^117^ organizes a variety of cell-scale datasets relevant to COVID-19; these may be of particular importance to intracellular modeling components of the project. The COVID-19 Disease Map^118^ also has a wealth of host-pathogen interaction data. The Human Biomolecular Atlas Program (HuBMAP)^119^ is creating detailed maps of the human respiratory system at cell- and molecular-scale resolution; this will be an excellent data source for tissue geometry in later versions of the model.

## Methods

### PhysiCell: agent-based cell modeling with extracellular coupling

PhysiCell is an open source simulation agent-based modeling framework for multicellular systems in 2D and 3D dynamical tissue environments^109^. (See Metzcar et al. (2019) for a general overview of agent-based modeling techniques in tissue-scale biology^120^.) In this framework, each cell (of any type) is an off-lattice agent with independent cell cycle progression, death processes, volume changes, and mechanics-driven movement. Each cell agent can have independent data and models attached to it, allowing substantial flexibility in adapting the framework to problems in cancer biology, microbiology, tissue engineering, and other fields. PhysiCell is coupled to BioFVM (an open source biological diffusion solver)^121^ to simulate the chemical microenvironment. As part of this coupling, each individual agent can secrete or uptake diffusing substrates and track the total amount of material entering and leaving the cell.

Relevant applications of PhysiCell-powered models have included modeling cancer nanotherapy^122^, oncolytic virus therapies^123^, tissue biomechanical feedbacks during tumor metastatic seeding^124^, and cancer immunology^109,125,126^. The platform was built with a focus on computational efficiency and cross-platform compatibility: the same source code can be compiled and run without modification on Linux, MacOS, and Windows, and simulations of up to 10 diffusing substrates on 10 mm^3^ of tissue with 10^4^ to 10^6^ cells are routinely performed on desktop workstations. The platform has been combined with high-throughput computing^125^ and active learning techniques^126^ to power large-scale model exploration on high performance computing resources.

### Integration of intracellular models in PhysiCell agents

Custom functions can be attached to individual cell agents to model molecular-scale, intracellular processes and to couple these with cell phenotypic parameters. These internal models are often implemented as systems of ODEs. For example, cell uptake of diffusing substrates can be coupled with a metabolism model that is defined by a system of ODEs, and the resulting energy output can be used to set the cycle progression and necrotic death probability of a cell^127^. For small systems of ODEs, these models are currently coded “by hand” with standard finite difference techniques. More complex models are written in systems biology markup language (SBML)^128^ for reliable scientific communication. Development versions of PhysiCell can read and integrate an individual SBML-encoded model in each cell agent using *libRoadrunner—a* highly efficient SBML integrator^129^. Similar approaches have been used to integrate Boolean signaling networks^130^ in PhysiCell in the PhysiBoSS extension^131^.

These approaches will initially be used to assess (1) viral replication dynamics in each cell agent, (2) cell death responses to viral load, (3) cell responses to interferons, and (4) changes in the virion endocytosis rate based on the availability of ACE2 and its receptor trafficking dynamics.

### Systems-scale lymphatic systems modeling

The adaptive immune response to viral infection at any tissue is triggered by the underlying lymph node (LN)-based systemic immune system when the infected/activated antigen presenting cells such as the dendritic cells start to migrate from the tissue to the lymphatic circulation. The time course of arrival of the dendritic cells in the lymph node is set by the departure of activated dendritic cells from the spatial model. In the LN, the proliferation, activation and clearance of the two types of helper T cells (*T_H1_* and *T_H2_*) and cytotoxic T cells (*T_C_*) are simulated by a set of ODEs (Eq. 55–58) which then go to inform the spatial model of arrival rates of both helper T cells and cytotoxic T cells. The LN interaction network among the DC, Th1, Th2 are adopted assuming the entire secretory cascades of cytokines and interleukins are implicit to their concentrations. We assume that the DCs start secreting the primary pro-inflammatory cytokines while the Th1 and Th2 initiate the secondary pro- and anti-inflammatory secretions. On the consequence, all these secretions regulate the activities of the helper and cytotoxic T cells^132^. Any transport between the LN and PhysiCell model is done only in integers and these events are performed before any diffusion or continuum process to attempt to reduce the error in decoupled solvers^133^.

### HPC-driven model exploration and parameterization

The concurrent growth and advancements in the three areas of 1) mechanistic simulation modeling, 2) advanced, AI-driven model exploration algorithms, and 3) high-performance computing (HPC) provides the opportunity for large-scale exploration of the complex design spaces in detailed dynamical simulation models. However, if we do not take deliberate efforts to formally facilitate this intersection across our research communities, we risk producing a series of disparate individual efforts, limited in interoperability, transparency, reproducibility and scalability. The EMEWS (extreme model exploration with Swift) framework^134^ was developed to directly address this issue and to provide a broadly applicable cyberinfrastructure to lower the barriers for utilization of advanced, large-scale model exploration on HPC resources. The EMEWS paradigm allows for the direct exploitation of cutting edge statistical and machine learning algorithms that make up the vibrant ecosystem of free and open source libraries that are continually added to and updated as research frontiers are expanded, all while controlling simulation workflows that can be run anywhere from desktops to campus clusters and to the largest HPC resources.

We have utilized EMEWS for learning-accelerated exploration of the parameter spaces of agent-based models of immunosurveillance against heterogeneous tumors^125,126^. The approach allowed for iterative and efficient discovery of optimal control and regression regions within biological and clinical constraints of the multi-scale biological systems. We have applied EMEWS across multiple science domains^135–138^ and developed large-scale algorithms to improve parameter estimation through approximate Bayesian computation (ABC) approaches^139^. These approaches, applied to the multi-scale modeling of SARS-CoV-2 dynamics, will provide the ability to robustly characterize model behaviors and produce improved capabilities for their interpretation.

### nanoHUB platform

The nanoHUB platform (nanohub.org)^140^ is a free, cloud-based service offering lectures, tutorials, and, of particular interest to us, interactive Web-based simulation tools. As its name implies, it is primarily focused on nanoscale science education and research. To make their simulation tools easier to use, nanoHUB provides a custom toolkit for developing graphical user interfaces (GUIs). However, since 2017, they have adopted and promoted the use of Jupyter notebooks^141^, with accompanying Python modules to provide GUI widgets and visualization. Cloud-based computing and data analysis platforms are well established now, in both academic and commercial settings. Those platforms, such as nanoHUB, that provide easy-to-use web-based GUIs and APIs and offer affordable pricing will likely have their rate of adoption continue to increase, especially among researchers who may lack the expertise or resources to install complex pieces of software.

### xml2jupyter and cloud deployment of PhysiCell models

Compiled PhysiCell models generate executable software that runs at the command line. Model parameters are set by editing XML (extensible markup language) configuration files, and the models save data as a combination of vector graphics outputs (scalable vector graphics: SVG) and XML and MATLAB data (.mat) files based on the draft MultiCellDS data standard^142^.

To facilitate rapid cloud-hosted dissemination of PhysiCell-powered models on the nanoHUB platform, we developed *xml2jupyter* to automatically generate a Jupyter-based GUI for any PhysiCell model^143^. The Jupyter notebook includes widgets to set parameters, initiate a simulation run, and visualize diffusing substrates and cell agents. In turn, we also developed a protocol to deploy the PhysiCell model and the Jupyter notebook interface on nanoHUB as a cloud-hosted, interactive model. This allows developers to rapidly convert a locally executable command-line model to a cloud-hosted shared model with graphical interface in a matter of minutes to hours (depending upon testing speed on nanoHUB).

In our rapid prototyping, we use rapidly-generated nanoHUB apps for scientific communication across disciplines; virologists, pharmacologists, and other domain experts can explore and visualize the model prototypes without need to download, compile, or understand the code. This facilitates faster multidisciplinary dialog and helps to draw in broader community feedback and contributions.

### Modular design

The model is being evolved with a modular architecture. The overall model and each individual model component (submodel) have a separate GitHub software repository in the pc4COVID-19 GitHub organization, available at https://github.com/pc4covid19.

Each submodel repository consists of a *master* branch that matches the latest numbered release and a *development* branch. Contributors will fork the development branch, complete their milestones, and submit a pull request to incorporate their progress in the development branch. Whenever a submodel team is ready to make a numbered release, they will use a pull request from the development branch to the master branch and create a numbered release.

The overall model framework and each submodel will keep a versioned design document to include:

- A unique name for the model component
- A clear version number and last update timestamp
- A list of contributors, including 1-2 chief scientists who serve as primary points of contact
- A “plain English” description of the primary purpose of the component
- A statement of model inputs with units of measure
- A clear statement of the biological hypotheses and assumptions of the component
- A record of the current mathematical form of the model (generally maintained in a separate Overleaf LaTeX document), with a snapshot of the equations in the main design document
- Any computational implementation details needed to understand the code
- A link to a GitHub repository
- A list of model parameters, units, biophysical meaning, best estimate, and data source(s) for the parameter estimate (see the discussion in MultiCellDS^142^)
- A clear list of model outputs with units
- A set of qualitative and/or quantitative **unit** tests to ensure proper functionality of the module.

A snapshot of this design document will be included in each release of the (sub)model.

The overall model releases will include a clear list of the version of each submodel included in its release.

### Coalition structure

After group discussion and prioritization, coalition members self-assigned themselves to one or more subteams responsible for developing the submodels. Each *subteam* has 1-2 chief scientists in charge of managing development, while a technical contact approves pull requests from the subteam’s contributors and coordinates with the integration team (see below). The submodel chief scientist(s) meet regularly with their team to assign tasks, set milestones, and assess when to make a release. The submodel chief scientist(s) also coordinate their progress with the other submodel teams.

The *integration team*—consisting of the overall leads (as of October 2021: Paul Macklin, Randy Heiland, Michael Getz, and Yafei Wang) and other contributors—is responsible for developing and maintaining the overall integrated model, integrating and testing updated submodels, and providing technical support to the subteams.

The *core team* consists of the overall leads and the chief scientists. They meet to coordinate progress of the submodels, refine project scope, exchange ideas on model hypotheses, evaluate community feedback, and plan overall strategy. They cooperate with the overall leads to create model releases (which will always bundle the most stable version of each submodel), update the nanoHUB models, and update the *bioRxiv* preprint.

The current list of subteams can be found in **Box 3**.

#### Box 3: Current subteams.

##### Integration

Coordinates overall project and leads multiscale model integration and dissemination.

*Chief scientist*(*s*): Paul Macklin, Michael Getz

*Technical contact*(*s*): Michael Getz, Randy Heiland, Yafei Wang

*Members*: Paul Macklin, Randy Heiland, Michael Getz, Yafei Wang

##### Viral Replication

Builds the submodel of viral replication (and release) within individual cells.

*Chief scientist*(*s*): James Faeder

*Technical contact*(*s*): Yafei Wang, Ali Sinan Saglam

*Members:* Jim Faeder, Yafei Wang, Paul Macklin, Ali Sinan Saglam, Caroline Larkin

##### Infected cell response

Builds the submodel of individual cell responses to infection, such as secretion of chemokines and apoptosis.

*Chief scientist*(*s*): Jason Shoemaker, James Glazier, Sara Hamis, Fiona Macfarlane

*Technical contact*(*s*): Jordan Weaver, Josua Aponte, Sara Hamis, Fiona Macfarlane

*Members:* Jason Shoemaker, Jim Faeder, Penny Morel, James Glazier, Ashok Prasad, Elsje Pienaar, Jordan Weaver, T.J. Sego, Josua Aponte, Yafei Wang, Sara Hamis, Fiona Macfarlan

##### Pharmacodynamics

Modifies the submodels to simulate pharmacologic interventions.

*Chief scientist*(*s*): Robert Stratford, Morgan Craig

*Technical contact*(*s*): Tarunendu Mapder, Yafei Wang, Sara Hamis, Fiona Macfarlane

*Members:* Robert Stratford, Morgan Craig, Sara Quinney, Mark AJ Chaplain, Tarunendu Mapder, Yafei Wang, Sara Hamis, Fiona Macfarlane, Richard F. Bergstrom

##### Receptor trafficking

Builds the submodel of ACE2 receptor trafficking, including virus binding and endocytosis.

*Chief scientist*(*s*): Padmini Rangamani

*Technical contact*(*s*): Andy Somogyi

*Members:* Padmini Rangamani, Andy Somogyi

##### Tissue immune response

Builds the submodels of individual immune cells and their interactions within an infected tissue.

*Chief scientist*(*s*): Morgan Craig, Courtney Davis, Amber Smith, Adrianne Jenner, Penny Morel

*Technical contact*(*s*): Adrianne Jenner

*Members:* Adrianne Jenner, Courtney Davis, Morgan Craig, Amber Smith, Penny Morel, Sofia Alfonso, Rosemary Aogo, Elsje Pienaar, Dennis Hou

##### Lymph node

Builds the submodel of immune cell expansion at nearby lymph nodes to drive immune cell recruitment.

*Chief scientist*(*s*): Gary An, Tarunendu Mapder

*Technical contact*(*s*): Tarunendu Mapder

*Members:* Gary An, Chase Cockrell, Marc-Andre Rousseau, James Glazier, T.J. Sego, Tarunendu Mapder, Juliano Ferrari Gianlupi

##### Tissue damage

Builds models of tissue damage (and potentially recovery).

*Chief scientist*(*s*): Ashlee Ford Versypt, Amber Smith

*Technical contact*(*s*): Mohammad Aminul Islam

*Members:* Amber Smith, Ashlee Ford Versypt, Thomas Hillen, Mohammad Aminul Islam

##### Drug testing/experiment

Explores drug inhibiting virus endocytosis and replication in cell culture

*Chief scientist*(*s*): Aarthi Narayanan

*Technical contact*(*s*): Kenneth Risner

*Members:* Aarthi Narayanan, Kenneth Risner

##### SBML integration

Refines PhysiCell integration with libRoadrunner to support direct execution of SBML models

*Chief scientist*(*s*): Randy Heiland

*Technical contact*(*s*): Randy Heiland, Andy Somogyi

*Members:* Andy Somogyi, Randy Heiland, Furkan Kurtoglu, Pablo Maygrundter, Jim Faeder

##### Visualization and analytics

Refines standalone and integrated visualization and analytics for nanoHUB apps.

*Chief scientist*(*s*): Randy Heiland, Amber Smith, Courtney Davis

*Technical contact*(*s*): Randy Heiland, Dennis Hou

*Members:* Randy Heiland, Amber Smith, Courtney Davis, Hadi Taghvafard, Andy Somogyi, Furkan Kurtoglu, Pablo Mayrgundter, Dennis Hou

### Three main phases of community-driven development

#### Phase 1: Building the coalition and model infrastructure

In the first phase, the overall and integration leads build the overall tissue model structure (a model that integrates several independent *submodels*) and create “placeholder” models that serve as working proof-of-concept starting points for further expansion. This phase also builds and organizes the subteams responsible for the submodels and provides them with training and documentation on the model and submodel architecture.

We anticipate that Phase 1 will require six-to-eight weeks, although Phases 1 and 2 may overlap as individual subteams assume full leadership of their submodel code repositories.

#### Phase 2: Community-driven development

In this phase, the integration team transitions the primary development of each of the submodels to appropriate domain experts in the subteams, to ensure that each submodel reflects the best available science. During this phase, the integration team supports each subteam in mathematical model development, PhysiCell implementation, and nanoHUB deployment for rapid subteam testing, dissemination, and community feedback on the submodels.

The integration team continues to lead overall model integration, testing, and deployment as a cloud-hosted model, and development of further infrastructure (e.g., HPC investigations) and PhysiCell and xml2jupyter refinements needed by the subteams (e.g., full support for SBML for molecular-scale model integration).

Once the integrated model can qualitatively produce expected viral and immune behaviors (as determined by the core group) and receives no major domain expert or community critiques, the major goal of the coalition (and this paper) will be met: to create a SARS-CoV-2 modeling framework suitable for subsequent calibration, validation, and exploration by the community. It will be submitted to scientific peer review, disseminated to the community, and maintained. This will mark the conclusion of Phase 2.

We anticipate that Phase 2 will require six to twelve months.

#### Phase 3: widespread scientific use and model maintenance

Once the overall model and submodels are largely complete, the model will be a mature, open source community resource available for use in scientific investigations. Moreover, due to our iterative approach, we envision that teams will have begun using earlier versions of the model for investigations by this point. The integration team will transition to supporting parallel investigations by independent groups using the models, and aggregating and sharing best data, parameter estimation, and results. The integration team and subteams will coordinate to encourage full scientific publication of the overall model, submodels, and resulting scientific investigations.

This phase will also focus on code hardening, documentation, and development of training and outreach materials. This phase is critical for *knowledge capture*, so that the model remains usable beyond the involvement of the original teams and can be rapidly adapted to emerging health challenges in the future. We also envision continued refinement of the model to reflect improved biological knowledge.

### Iterative development

We use ***iterative prototyping*** using lessons learned from each step to drive iteration towards improving the model. Each submodel undergoes its own development sprints, contained within a broader development cycle for the overall integrated model (See **Fig. 0.1**.).

**Fig 0.1:**
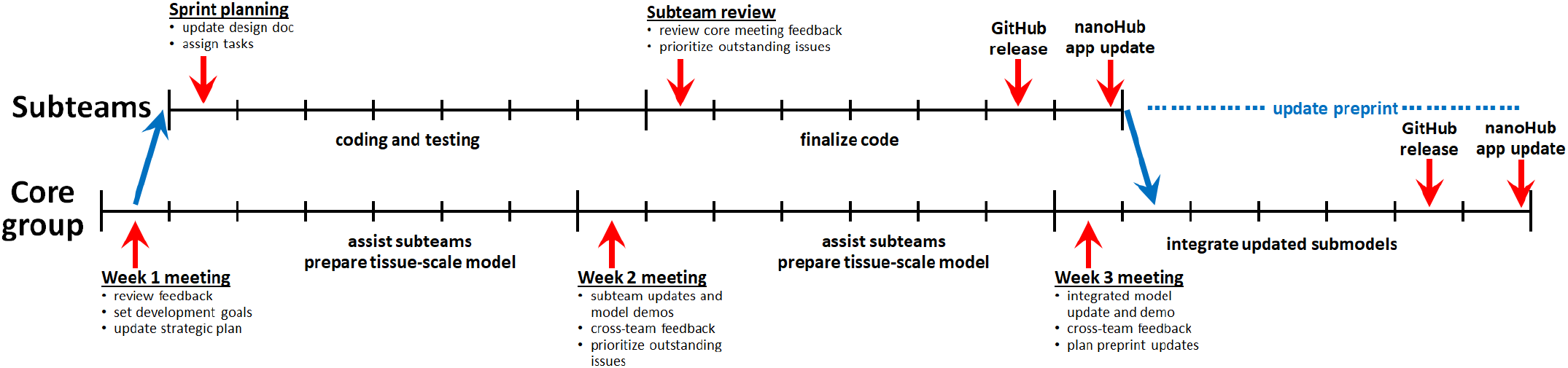
Overall development cycle (3-week example): Throughout the overall development cycle, the core team (integration leads + subteam leads) set priorities and coordinate work of the subteams. Each subteam performs a short sprint to update its submodel. At the end of the development cycle, the integration team bundles the most up-to-date submodels for the next overall model release, while the subteams update the preprint and refine domain knowledge and mathematics for the next sprint.

#### Overall integrated model development cycle

We aim for a short development cycle for the overall integrated model, although development cycles may last longer to accommodate building computational infrastructure, development of new core features to support model development, and training.

##### Start of cycle

The design cycle starts with an initial core team meeting where we discuss feedback from prior design cycles and set priorities for the current design cycle. In particular, we discuss:

- What changes are needed to the submodels? Prioritize changes that can be made within a 7-10 day sprint.
- What changes are needed in the overall integrative framework to facilitate the improved submodels? Set framework goals for early and mid-cycle development.
- Are any funding, personnel, scope, or other changes needed?

Within the working week, the subteams meet to further set and accomplish their sprint goals. (See **Submodel** design cycle). The integration team (1) works on refinements to the PhysiCell and nanoHUB frameworks to facilitate subteam work, (2) provides technical consulting to the subteams to implement their model refinements, and (3) makes any final edits needed to the preprint from the last design cycle.

##### Mid-cycle advances

The design cycle continues with a core team meeting to discuss the current subteam model sprints:

- Each team gives a brief report on their model advances and a live demo of either the standalone C++ code or a nanoHUB submodel app.
- The teams “cross-pollinate” to exchange ideas and give feedback on each of the submodels.
- The core team decides on any additional changes needed to continue the design cycle.
- The integration team and subteam chief scientists set final deadlines to end the sprints and release the updated submodels.

Within the working week, the subteams continue and complete their developing and testing for their respective sprints, create new submodel releases on GitHub, and update their submodel nanoHUB apps. The integration team continues support for the subteam work and completes any changes to the overall integrative model needed for the upcoming integration.

As the subteams advance towards their releases, the integration team updates and tests the respective submodels in the overall framework and updates the overall nanoHUB app.

##### End of cycle

The design cycle nears completion with a core team meeting to discuss integration and preprinting:

- The integration team gives an update on current integration testing.
- The team coordinates any remaining submodel releases.
- The team sets plans for updating the preprint.

Within the working week, the **subteams** complete their releases (if not already complete in week 2) and begin revising the preprint. They also begin testing the integrated model as it comes online to integrate new simulation results and insights into the preprint.

The **integration team** updates the submodels, performs final testing, and creates both GitHub and nanoHUB releases. Once complete, the integration team joins the subteams on preprint updates. We note that in non-summer months, the coalition was not able to maintain a three-week development pace.

#### Submodel design cycle

Each submodel is developed in parallel with a unified design cycle (a 7-to-14-day software *sprint)* in coordination with the other subteams during the weekly core team meetings and via a dedicated Slack workspace.

##### Start of sprint

The sprint cycle starts with an initial subteam meeting to communicate the results and priorities of the core team meeting to the subteam. The team edits the submodel design document, discusses any necessary changes to the mathematics and parameter values, and assigns implementation tasks. The team coordinates with the integration team via the Slack workspace for any needed assistance on model implementation.

##### End of sprint

The design cycle continues with a core team meeting to discuss the current subteam model sprints:

- Each team gives a brief report on their model advances and a live demo of either the standalone C++ code or a nanoHUB submodel app.
- The teams “cross-pollinate” to exchange ideas and give feedback on each of the submodels.
- The core team decides on any additional changes needed to continue the design cycle.
- The integration team and subteam chief scientists set final deadlines to end the sprints and release the updated submodels.

Within the working week, the **subteams** continue and complete their developing and testing for their respective sprints, create new submodel releases on GitHub, and update their submodel nanoHUB aps. The **integration team** continues support for the subteam work and completes any changes to the overall integrative model needed for the upcoming integration.

As the subteams advance towards their releases, the **integration team** updates and tests the respective sub-models in the overall framework and updates the overall nanoHUB app.

See **Appendix 4: Submodel development details** for more implementation details.

## Results

### Summary of Versions 1-4

The **Version 1** model prototyped the initial modular framework: a layer of susceptible, uninfected cells was modeled as cell agents. Each cell agent had an independent viral replication model (a system of five ordinary differential equations), and a (death) response model to viral load. As virus was assembled in each cell, it was released into the extracellular environment. Extracellular viral dissemination was modeled as a reaction-diffusion equation, and virus binding to (susceptible) cells was modeled as an uptake term that transferred extracellular diffusing virus to the intracellular ODE model. The model was initialized by placing a single virion in the center of the (extracellular) simulation domain. The model was able to capture a single expanding viral plaque, with greatest (single-cell) viral load in the center and expanding outwards, with greatest death and tissue disruption in the center of the plaque.

The **Version 2** model further modularized the framework into more distinct software modules of the submodels, while also introducing an ACE2 receptor trafficking model (four ODEs for internal and external bound and unbound receptors) to each cell agent. The model also introduced a more realistic starting condition of multiple virions entering the extracellular at random locations according to a user-defined multiplicity of infection (MOI). This model was able to further explore the impact of receptor trafficking limits on the rate of viral spread. Versions 1-2 were performed during Phase 1 (building the coalition and model infrastructure).

The **Version 3** model was the first released developed in Phase 2 (community-driven development). Its major change was to introduce an agent-based immune model of macrophages, neutrophils, and effector T cells. Resident macrophages, upon phagocytosing dead infected cells, secreted pro-inflammatory factors to stochastically recruit neutrophils and effector T cells that could hunt and kill infected cells. These model additions allowed us to explore the impact of increasing or decreasing resident macrophage and T cell populations.

The **Version 4** model integrated a lymphatic system model: a systems of ordinary differential equations now represent arriving dendritic cells that drive expansion of T cell populations. Immune cells traffic between the spatially-resolved tissue model (Versions 1-3) and this new lymphatic compartment. Version 4 also refined and expanded the version 3 minimal model of the tissue-level immune response to SARS-CoV-2. Dendritic cells and CD4^+^ T cells were added to capture antigen presentation dynamics and the interplay between macrophages and T cell signaling. Macrophage activation and phagocytosis mechanisms have also been refined, and we have introduced a model of cell death via pyroptosis. Fibroblast-mediated collagen deposition has been included to account for the fibrosis at the damaged site in response to immune response-induced tissue injury. Epithelial cells’ production of Type-I Interferons and Interferon Stimulated Genes’ effects on viral replication have been included.

Full details on these prior model versions can be found in the supplementary materials.

### Version 5 (developed November 20, 2020-July 19, 2021)

#### Biological hypotheses

The version 5 model added numerous changes from previous versions. Namely additions for later time scale immune players, plasma cells, B cells, and Antibody [Immunoglobulin (Ig)] were added allowing further system testing. Below, assumptions are indicated by X.C.Y, where X denotes prototype, C denoted modeling component, and Y denotes a biological hypothesis, for easy reference. Assumptions in **bold** have been added or modified from the previous model version.

5.T.1 Virus diffuses in the microenvironment with low diffusion coefficient
5.T.2 Virus adhesion to a cell stops its diffusion (acts as an uptake term)
5.T.3 Pro-inflammatory cytokine diffuses in the microenvironment
5.T.4 Pro-inflammatory cytokine is taken up by recruited immune cells
5.T.5 Pro-inflammatory cytokine is eliminated or cleared
5.T.6 Chemokine diffuses in the microenvironment
5.T.7 Chemokine is taken up by immune cells during chemotaxis
5.T.8 Chemokine is eliminated or cleared
5.T.9 Debris diffuses in the microenvironment
5.T.10 Debris is taken up by macrophages and neutrophils during chemotaxis
5.T. 11 Debris is eliminated or cleared
**5.T.12** Anti-inflammatory cytokine secretion is triggered at the site that a CD8^+^ T cell kills an epithelial cell for a set amount of time
5.T.13 Anti-inflammatory cytokine diffuses in the microenvironment
5.T.14 Anti-inflammatory cytokine is eliminated or cleared
5.T.15 Collagen is non-diffusive
**5.T.16** Immunoglobulin (Ig) diffuses in the microenvironment
**5.T.17** Virions and Ig react in tissue removing both components
**5.T.18** Reactive oxidative species (ROS) diffuses in the microenvironment
5.RT.1 Virus adheres to unbound external ACE2 receptor to become external (virus)-bound ACE2 receptor
5.RT.2 Bound external ACE2 receptor is internalized (endocytosed) to become internal bound ACE2 receptor
5.RT.3 Internalized bound ACE2 receptor releases its virion and becomes unbound internalized receptor. The released virus is available for use by the viral lifecycle model **V**
5.RT.4 Internalized unbound ACE2 receptor is returned to the cell surface to become external unbound receptor
5.RT.5 Each receptor can bind to at most one virus particle.
5.V.1 Internalized virus (previously released in 4.RT.3) is uncoated
5.V.2 Uncoated virus (viral contents) lead to release of functioning RNA
5.V.3 RNA creates viral protein at a constant rate unless it degrades
5.V.4 Viral RNA is replicated at a rate that saturates with the amount of viral RNA
5.V.5 Viral RNA undergoes constitutive (first order) degradation
5.V.6 Viral protein is transformed to an assembled virus state
5.V.7 Assembled virus is released by the cell (exocytosis)
5.VR.1 After infection, cells secrete chemokine
5.VR.2 As a proxy for viral disruption of the cell, the probability of cell death increases with the total number of assembled virions
5.VR.3 Apoptosed cells lyse and release some or all their contents
5.VR.4 Once viral RNA exceeds a particular threshold *(max_apoptosis_half_max),* the cell enters the pyroptosis cascade
5.VR.5 Once pyropotosis begins, the intracellular cascade is modelled by a system of ODEs monitoring cytokine production and cell volume swelling
5.VR.6 Cell secretion rate for pro-inflammatory increases to include secretion rate of IL-18
5.VR.7 Cell secretes IL-1β which causes a bystander effect initiating pyroptosis in neighboring cells
5.VR.8 Cell lyses (dying and releasing its contents) once its volume has exceeded 1.5× the homeo static volume
5.E.1 Live epithelial cells undergo apoptosis after sufficient cumulative contact time with adhered CD8^+^ T cells.
5.E.2 Live epithelial cells follow all rules of RT
5.E.3 Live epithelial cells follow all rules of V
5.E.4 Live epithelial cells follow all rules of VR
5.E.5 Dead epithelial cells follow all rules of D.
5.E.6 Infected epithelial cells secrete pro-inflammatory cytokine
5.E.7 Antigen presentation in infected cells is a function of intracellular viral protein
5.D.1 Dead cells produce debris
5.MPhi.1 Resident (unactivated) and newly recruited macrophages move along debris gradients.
5.MPhi.2 Macrophages phagocytose dead cells. Time taken for material phagocytosis is proportional to the size of the debris
5.MPhi.3 Macrophages break down phagocytosed materials
5.MPhi.4 After phagocytosing dead cells, macrophages activate and secrete pro-inflammatory cytokines
5.MPhi.5 Activated macrophages can decrease migration speed
5.MPhi.6 Activated macrophages have a higher apoptosis rate
5.MPhi.7 Activated macrophages migrate along chemokine and debris gradients
5.MPhi.8 Macrophages are recruited into tissue by pro-inflammatory cytokines
5.MPhi.9 Macrophages can die and become dead cells only if they are in an exhausted state
5.MPhi.10 Macrophages become exhausted (stop phagocytosing) if internalised debris is above a threshold
**5.MPhi.11** CD8^+^ T cell contact stops activated macrophage secretion of pro-inflammatory cytokine and switches to M2 phase, secreting anti-inflammatory cytokine.
5.MPhi.12 CD4^+^ T cell contact induces activated macrophage phagocytosis of live infected cells
5.N.1 Neutrophils are recruited into the tissue by pro-inflammatory cytokines
5.N.2 Neutrophils die naturally and become dead cells
5.N.3 Neutrophils migrate locally in the tissue along chemokine and debris gradients
5.N.4 Neutrophils phagocytose dead cells and activate
5.N.5 Neutrophils break down phagocytosed materials
5.N.6 Activated neutrophils reduce migration speed
5.N.7 Neutrophils uptake virus
**5.N.8** Neutrophils secrete ROS upon phagocytosis.
5.DC.1 Resident DCs exist in the tissue
5.DC.2 DCs are activated by infected cells and/or virus
5.DC.3 Portion of activated DCs leave the tissue to travel to the lymph node
5.DC.4 DCs chemotaxis up chemokine gradient
5.DC.5 Activated DCs present antigen to CD8^+^ T cells increasing their proliferation rate and killing efficacy (doubled proliferation rate and attachment rate)
5.DC.6 Activated DCs also regulate the CD8^+^ T cell levels in within a threshold by enhancing CD8^+^ T cell clearance.
5.CD8.1 CD8^+^ T cells are recruited into the tissue by pro-inflammatory cytokines
5.CD8.2 CD8^+^ T cells apoptose naturally and become dead cells
5.CD8.3 CD8^+^ T cells move locally in the tissue along chemokine gradients
5.CD8.4 CD8^+^ T cells adhere to infected cells. Cumulated contact time with adhered CD8^+^ T cells can induce apoptosis (See 4.E.1)
5.CD8.5 Activated DCs present antigen to CD8^+^ T cells, which increases the CD8^+^ T cell proliferation rate
5.CD8.6 Activated DCs also regulate the CD8^+^ T cell levels in within a threshold by enhancing CD8^+^ T cell clearance.
**5.CD8.7** CD8^+^ T cells have a max generation counter and will not proliferate after the set generation.
5.CD4.1 CD4^+^ T cells are recruited into the tissue by the lymph node
5.CD4.2 CD4^+^ T cells apoptose naturally and become dead cells
5.CD4.3 CD4^+^ T cells move locally in the tissue along chemokine gradients
5.CD4.4 CD4^+^ T cells are activated in the lymph node by three signals: (1) antigenic presentation by the DCs, (2) direct activation by cytokines secreted by DCs, (3) direct activation by cytokines secreted by CD4^+^ T cells.
5.CD4.5 CD4^+^ T cells are suppressed directly by cytokines secreted by CD4^+^ T cells.
**5.CD4.6** CD4^+^ T cells have a max generation counter and will not proliferate after the set generation.
5.F.1 Fibroblast cells are recruited into the tissue by anti-inflammatory cytokine
5.F.2 Fibroblast cells apoptose naturally and become dead cells
5.F.3 Fibroblast cells move locally in the tissue along up gradients of anti-inflammatory cytokine
5.F.4 Fibroblast cells deposit collagen continuously
5.LN.1 Lymph node is activated through DC presentation of antigen in the lymph node leading to recruitment of CD8^+^ T cells, CD4^+^ T cells and Ig to the local tissue.
**5.LN.2** Lymph node transport time can be simulated by DDEs in the lymph node.
**5.LN.3** In the lymph-node DCs directly participate in B cell activation and CD8^+^ and CD4^+^ T cell re cruitment.
**5.LN.4** In the lymph-node DCs regulate both the activation and deactivation of CD8^+^ T cells.
**5.LN.5** B cell activation requires two simultaneous signals: from DC and from CD4^+^ T cells.
**5.LN.6** As soon as activated, B cells start to differentiate into plasma cells.
**5.LN.7** Plasma cells are the major contributor of the antibody (IgM/IgG).
**5.LN.8** Antibodies are passively diffusible through the tissue barrier and able to bind and neutralize free virions.

### Key mathematical details retained from prior model versions

#### Extracellular virion transport (Tissue submodel ***T***)

To rapidly implement extracellular viral transport using existing model capabilities, we approximated the process as diffusion with a small diffusion coefficient as in prior nanoparticle models. Using the standard BioFVM formulation^121^, if *ρ* is the concentration or population density of virions (virions / μm^3^), then the population balance is modeled as a partial differential equation (PDE) for diffusion, decay, and sources and sinks from interactions with cells:

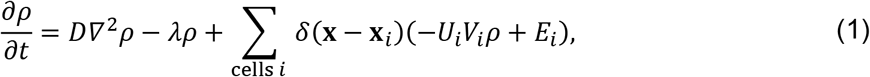

where *D* is the diffusion coefficient, *λ* is the net decay rate (which can include other removal processes), *δ* is the Dirac delta function, *x_i_* is the position of the center of cell *i*, *U* is the uptake rate (by adhering to ACE2 and initiating endocytosis), *V* is the volume of cell *i*, and *E* is the virion export rate from the cell. Note that in the default BioFVM implementation, uptake processes are spread across the volume of a cell.

Note that virus propagation may require more explicit modeling of cell-cell surface contact in later versions, and cilia-driven advective transport and virion deposition (e.g., through airway transport).

#### Intracellular viral replication dynamics (Virus intracellular model ***V***)

Within each cell, we track *V* (adhered virions in the process of endocytosis), *U* (uncoated viral RNA and proteins), *R* (viral RNA ready for protein synthesis; *R* = 1 denotes the total mRNA of one virion), P (synthesized viral proteins; P = 1 denotes sufficient viral protein to assemble a complete virion), and A (total assembled virions ready for exocytosis). Virion import (a source term for V) is handled automatically by the mass conservation terms for PhysiCell in the PDE solutions.

We model these dynamics of internalized virions through a simplified system of ODEs:

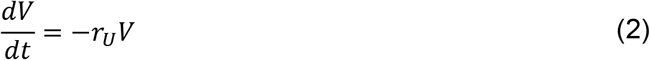

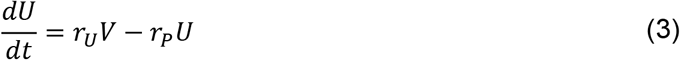

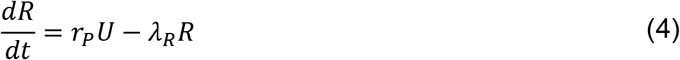

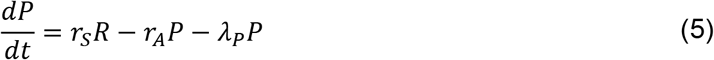

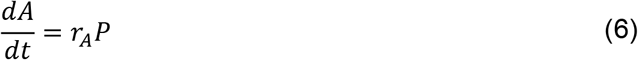

Here, *r_U_* is the viral uncoating rate, *r_P_* is the rate of preparing uncoated viral RNA for protein synthesis, *r_s_* is the rate of protein synthesis, *r_A_* is the rate of virion assembly, *λ_R_* is the degradation rate of RNA, and *λ_P_* is the degradation rate of viral protein. We model exocytosis by setting the net export *E* of the assembled virions, in units of virions per time:

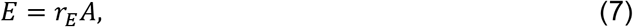

where *r_E_* is the assembled virus export rate.

#### Cell response (Viral response submodel ***VR***)

In this proof of concept prototype, we modeled the cell death response to cell disruption but did not model interferon processes. As a simplification, we modeled cell disruption as correlated with assembled virions *A*, and we used a Hill function to relate the apoptosis rate of a cell to *A*:

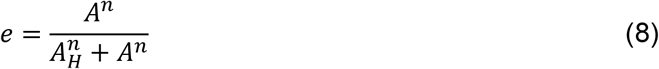

where *e* is the effect, *n* is the Hill coefficient, and *A_H_* is the number of virions at which half of the maximum effect is achieved. After calculating this effect *e*, we set the (non-necrotic) death rate as

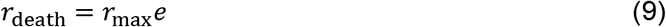

where *r*_max_ is the maximum death rate (at full effect, *e* = 1). As analyzed for agent-based models with stochastic death rates^109,144^, in any time interval [*t*, *t*+Δ*t*], the cell has probability *r*_death_Δ*t* of starting a death process, and the mean cell survival time (for fixed *e* and thus fixed *r*_death_) is 1/*r*_death_.

In PhysiCell, we can set the lysing cells to release any fraction (0 ≤ *f*_release_ ≤ 1) of *V*, *A*, *U*, *R*, and *P* into the extracellular environment as diffusing substrates.

#### ACE2 receptor trafficking (submodel ***RT***)

For each cell, we track *R*_EU_ (external unbound ACE2 receptors), *R*_EB_ (external virus-bound receptors), *R*_IB_ (internalized virus-bound receptor), and *R*_IU_ (internalized unbound receptor). We model hypotheses 2.RT.1-2.RT.5 as a system of ordinary differential equations. To attempt to deal with a discrete to continuum transition the rule for low virion uptake by cells was changed to a probability draw when flux values are under one (i.e less than one virion will be recruited in a time-step)

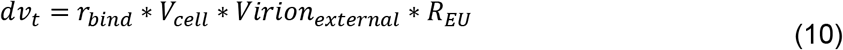

If 0 < *dv_t_* < 1

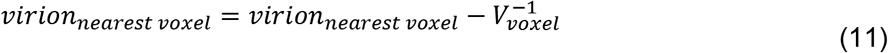

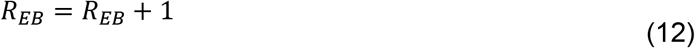

If *dv_t_* > 1

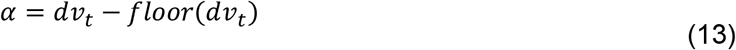

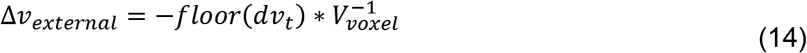

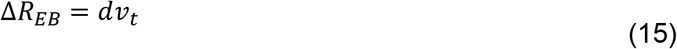

if *U*(0,1) < *α*,

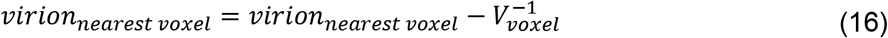

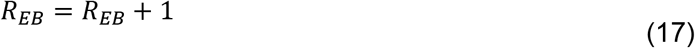

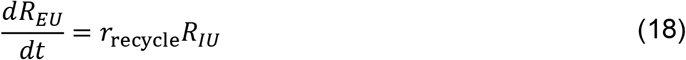

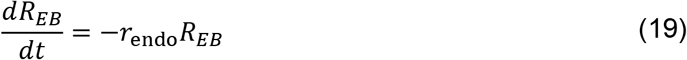

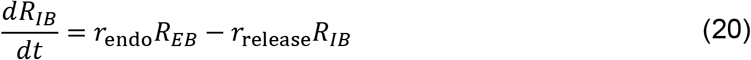

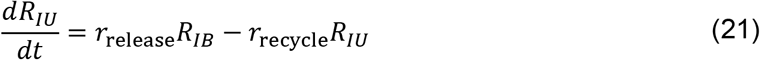

Thus, the virus endocytosis rate varies with the availability of unbound externalized ACE2 receptor, as expected. To link with the viral replication submodel, the unbinding of virus from internalized receptor must act as a source term for the internalized virus:

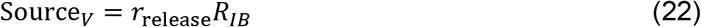

#### Intracellular viral replication dynamics (Virus lifecycle model ***V***)

We make a small modification to the internalized virus model to account for the coupling with the receptor traf-ficking model:

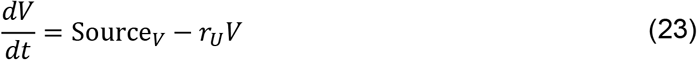

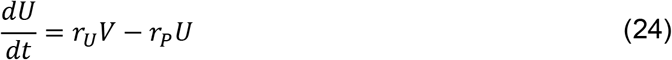

The model now incorporates viral RNA replication within the host cell. Only the viral RNA ordinary differential equation is directly modified as follows:

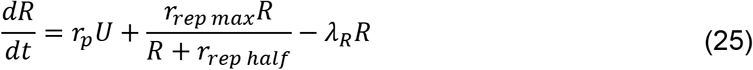

Here, r_rep max_ is the maximum replication rate of viral RNA and r_rep half_ represents the viral RNA concentration where the viral replication rate is half of r_rep max_.

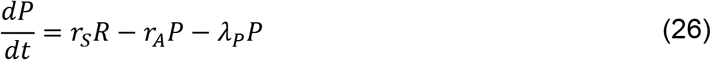

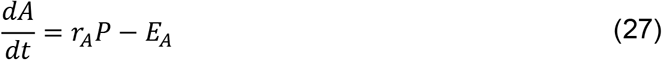

We model exocytosis by setting the export rate *E_A_* of the assembled virions, in units of virions per time where only whole virions are secreted:

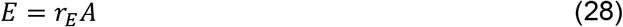

#### Extracellular transport (Tissue submodel ***T***)

Extracellular densities of pro-inflammatory cytokine and chemokine were modelled using the standard BioFVM formulation^121^, similar to that for extracellular virus (introduced above), i.e.:

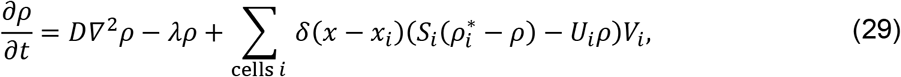

where *D* is the diffusion coefficient of each substrate, *λ* is the net decay rate, *δ* is the discrete Dirac delta function, *x_i_* is the position of the centre of cell *i*, *S_i_* is the secretion rate of cell *i*, 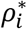 is the saturation density at which cell *i* stops secreting, *U_i_* is the uptake rate of the substrate by cell *i*, and *V_i_* is the volume of cell *i*. The concentration *ρ*, represents the density of pro-inflammatory cytokine *ρ*_cytokine_, chemokine *ρ*_chemokine_ or dead cell debris *ρ*_debris_. Similarly, diffusion, decay, secretion, and uptake parameters are all substrate specific rates, i.e. the diffusion coefficients are *D*_cytokine_, *D*_chemokine_ and *D*_debris_; the decay rates are *λ*_cytokine_, *λ*_chemokine_ and *λ*_debris_; the secretion rates are *S*_cytokine_, *S*_chemokine_ and *S*_debris_; the uptake rates are *U*_cytokine_, *U*_chemokine_ and *U*_debris_; and the saturation densities are 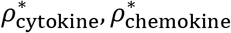 and 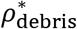.

#### Cell response (Viral response submodel ***VR***)

We made a small addition to the cell response model. After infection, cells start secreting chemokine at a rate

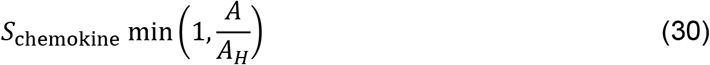

where *A* is the intracellular assembled virion count and *A_H_* is the number of assembled virions at which half of the maximum effect of virus-induced cell apoptosis is achieved. Secretion continues until the cell dies either through lysis or CD8^+^ T cell induced apoptosis.

#### Signaling, degradation, and phagocytosis of apoptotic cells (Dead cell dynamics ***D***)

Cells that die release debris that attracts phagocytes and signals that that they can be cleared from the microenvironment. They secret these signals at a rate *S*_debris_.

#### Chemotaxis (Chemotaxis model ***MPhi*, *N***, and ***CD8***)

Macrophages and neutrophils undergo chemotaxis up the chemokine gradient and dead-cell debris gradients released by infected cells and dead cells respectively. The velocity of cell chemotaxis is

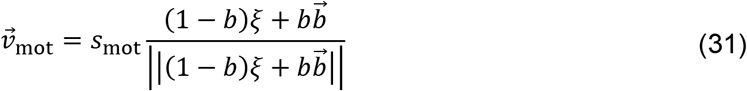

where *s_mot_* is the speed of chemotaxis (cell-type-specific), 0 ≤ *b* ≤ 1 is the migration bias (also cell-type-specific), *ξ* is a random unit vector direction in 3D (or 2D) and 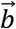 is the migration bias direction defined by

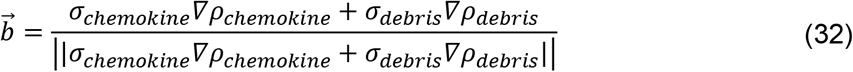

where *σ_chemokine_* and *σ_debris_* are the sensitivity of chemotaxis along either the chemokine or dead-cell debris gradient. CD8^+^ T cells also undergo chemotaxis, but along the chemokine gradient, i.e. *σ_debris_* = 0 and *σ_chemokine_* = 1. Chemotaxing cells take up chemokine at a rate *U_chemokine_*.

#### Phagocytosis dynamics (Phagocytosis of apoptotic cells ***MPhi*** and ***N***)

Once a macrophage or neutrophil has found a cell to phagocytose, it reduces its speed from *s_mot,a_* (active chemotaxis speed) to *s_mot,p_* (phagocytosis/attached speed) and starts searching locally for material to phagocytose.

If there is a dead cell in contact with a macrophage or neutrophil (i.e., if there is a dead cell in the cell’s ~30 *μm* mechanical interaction voxel as in PhysiCell^109^), the immune cell will phagocytose the dead cell with rate *r_phag_*, which is cell-type specific and reflects the efficacy with which each immune cell subtype clears debris. If the immune cell is in contact with a dead cell over a period of [*t*, *t* + Δ*t*], then the probability of phagocytosis is *r_phag_*Δ*t*. When an immune cell phagocytoses a dead cell, the immune cell absorbs the volume of that cell and subsequently increases its volume, i.e., the phagocytosing cell gains:

a. all of the dead cell’s fluid volume.
b. all of the dead cell’s nuclear solid and cytoplasmic solid volume (which are added to the nuclear cytoplasmic solid volume)

This implies that after phagocytosis within time Δ*t*, the volume of a macrophage or neutrophil *i* that phagocytoses a dead cell *j* will be given by

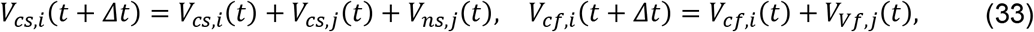

where *V_cs,k_* is the volume of the cytoplasmic solid volume in cell *k*, *V_ns,k_* is the volume of nuclear solid volume in cell *k*, *V_cf,k_* is the cytoplasmic fluid volume in cell *k*, and *V_Vf,k_* is the total fluid volume of cell *k*. Because this will typically increase the cell’s volume above its “target” equilibrium volume, the standard PhysiCell volume model^109^ will begin to shrink the cell’s volume back towards its resting volume, allowing us to model degradation of phagocytosed materials. After phagocytosing dead material, macrophages start secreting pro-inflammatory cytokines at a rate *S_cytokine_*.

#### Neutrophil viral clearance (***N***)

Neutrophils take up extracellular virus at a rate *U*. We assume this uptake rate is equivalent to the ACE2 receptor binding rate *r_bind_*.

#### Immune cell recruitment (***Mphi*, *N***, and ***CD8***)

Macrophages and neutrophils are recruited to the tissue by pro-inflammatory cytokines through capillaries/vasculature in the lung. The density of vasculature accounts for approximately 8.8% of the tissue^145^. Accordingly, at the start of each simulation we randomly assign 8.8% of the tissue voxels as vasculature points through which immune cells arrive randomly throughout the course of the simulation. (Note that the v1-v5 models simulate a single layer of epithelium where immune cells are allowed to move freely through or just above the tissue; this 2-D formulation is implemented as a single layer of 3D voxels^109^.)

At regular time intervals Δ*t*_immune_, we integrate the recruitment signal to determine the number of immune cells recruited to the tissue. The number of cells recruited into the tissue between *t* and *t* + Δ*t*_immune_ varies with the pro-inflammatory cytokine recruitment signal across the tissue:

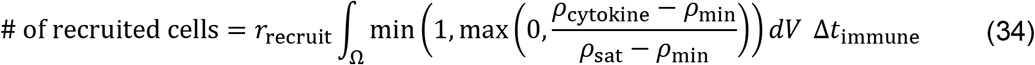

where Ω is the computational domain, *r*_recruit_ is the recruitment rate (per volume), *ρ*_min_ is the minimum recruitment signal, *ρ*_sat_ is the maximum (or saturating), and *ρ*_cytokine_ is the pro-inflammatory cytokine concentration. The value of *ρ*_min_, *ρ*_sat_, and *r*_recruit_ are cell-type specific, i.e. macrophages and neutrophils have different minimum and saturating recruitment signal concentrations which results in heterogenous arrival times into the tissue.

Recruited cells are randomly seeded at vessel locations. In the v5 model, we set Δ*t*_immune_ = 10 min. Notice that the number of recruited cells scales with duration of the time interval and the size of the tissue. Recruitment of immune cells always round down then do a probability draw to see if an extra cell is recruited. This leads to a smooth transition in earlier states of recruitment.

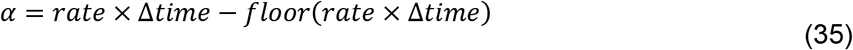

if *U*(0,1) < *α*,

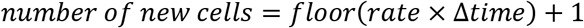

#### CD8^+^ T cell induction of infected cell apoptosis (***CD8*** dynamic model)

When a CD8^+^ T cell is in contact with a cell (based on PhysiCell’s mechanical interaction testing; see the note in phagocytosis above) with intracellular assembled virion is greater than 1, i.e. *A* > 1, the T cell attempts to attach to the infected cell with an attachment rate *r*_attach_. Following prior immune modeling work^125,126^, if the cell is in contact for a duration Δ*t*, then the probability of forming an attachment in that time period is *r*_attach_Δ*t*. While the cells are attached, the immune cell’s cumulative CD8^+^ T cell contact time is increased by Δ*t*. The T cell has a mean attachment lifetime *T*_attach_. Between t and *t* + Δ*t*, the probability of detaching is given by Δ*t*/*T*_attach_.

We assume that an infected cell will undergo apoptosis after its cumulative attachment time exceeds a threshold *T*_CD8_contact_death_. This can be either from a single or multiple T cell attachments. All attached T cells detach when a cell undergoes apoptosis. When CD8^+^ T cells adhere to another cell, their motility is turned off, i.e. *s_mot,p_* = 0, and when they detach from a cell, their speed returns to their active chemotaxis speed *s_mot,a_*.

#### IFN response

An early version of the IFN model was added in v4. In this model IFN interferes with protein synthesis reducing the rate of viral protein synthesis.

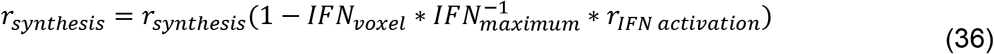

Where 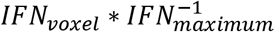 is bounded to 1.

IFN secretion is controlled through RNA detection and paracrine signals.

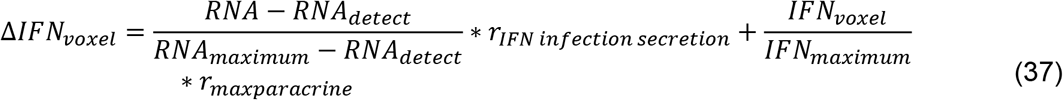

Where the fractions are bounded to 1.

#### Pyroptosis

Once the viral RNA levels within a cell exceed the threshold (R≥ 200), or IL-1β levels in the microenvironment reach the threshold (Cytokine≥ 100), the cell can undergo pyroptosis, a form of inflammatory cell death^146^. The pyroptosis cascade within each cell is modelled via a system of ODEs capturing the key components of the pathway. Many aspects of the pathway are dependent on whether the inflammasome base is still forming (*F_ib_* = 1) or whether it has formed (*F_ib_* = 0).

This then initiates the translocation of NF-κB into the nucleus at the rate *k_nfkb_ctn__*. The NF-κB can translocate back to the cytoplasm at the rate *k_nfkb_ntc__*. Therefore, we describe the evolution of nuclear NF-κB, *NFkB_n_*, through the equation:

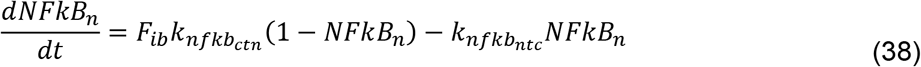

Nuclear NF-κB then regulates the transcription of inactive NLRP3 protein, we assume this transcription follows a standard hill function form with a transcription coefficient of *a_nlrp3_*. This inactive NLRP3 can then become activated at the rate *k_nlrp3_ita__*. We additionally assume some natural decay of the NLRP3 at the rate *d_nlrp3_*. We therefore describe the evolution of inactive NLRP3, *NLRP*3_*i*_, through the equation:

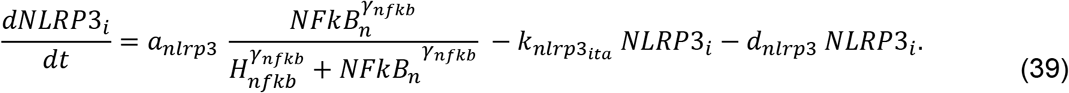

Once activated NLRP3 can oligomerize to form the inflammasome base, at the rate *k*_*nlrp*3_*atb*__. This process continues until enough NLRP3 has oligomerised/bound together to form the inflammasome base when *F_ib_* switches to zero. We describe the evolution of active NLRP3, *NLRP3_a_*, through:

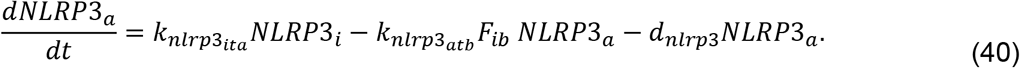

The evolution of bound NLRP3, *NLRP*3_*b*_, is described through:

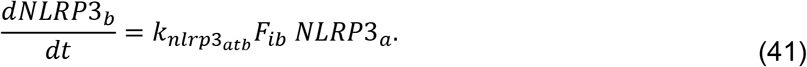

Once *NLRP*3_*b*_ ≥ 1, then the inflammasome base has formed and *F_ib_* switches to zero.

Once the inflammasome base has formed, ASC protein is recruited and binds to the inflammasome at the rate *k_asc_ftb__*. The change of bound ASC, *ASC_b_*, is described through:

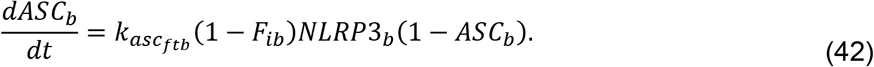

Pro-caspase 1 is then recruited to the inflammasome site, and is cleaved by bound ASC to become caspase 1, at the rate *k*_*c*1_*ftp*__. Therefore, caspase 1, *C*_1_, evolves through:

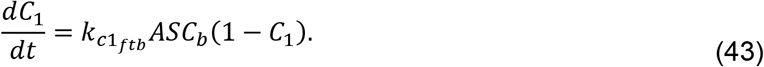

Caspase 1 has the capacity to cleave gasdermin and pro-interleukins within the cell. We assume that caspase cleavage of these molecules follows a hill function with coefficients *a*…. Therefore, the cleaved N terminal of gasdermin, *GSDMD_n_*, evolves through:

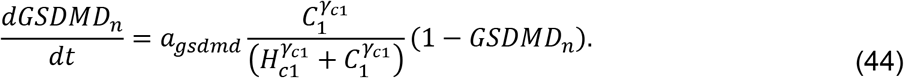

Similarly to NLRP3, we assume that pro-interleukin 1β, *IL*1*b_p_*, is transcribed by NF-κB and decays at the rate *d_il_*. The pro-form is then cleaved by caspase 1 to form the cytoplasmic form, that is:

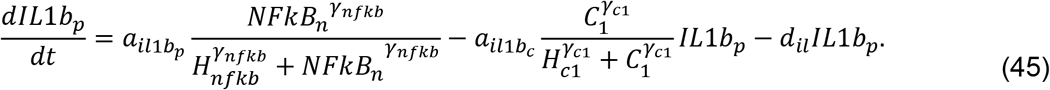

Cytoplasmic interleukin 1β, *IL*1*b_c_*, can also decay in the same manner as the pro-form. Additionally, *IL*1*b_c_*, can transport out of the cell via pores formed by gasdermin on the cell surface, at the rate *k*_*il*1*b_cte_*_. That is, *IL*1_*b*_*c*__, evolves via:

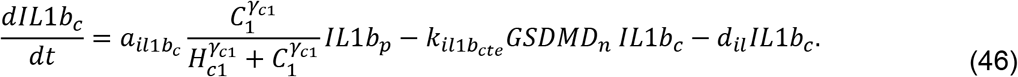

External interleukin 1β levels only depend on transport out of the cell in the ODE model,

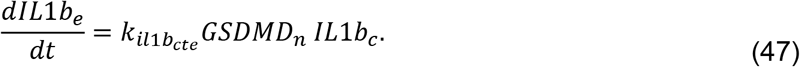

We only consider the level of cytoplasmic interleukin 18, *IL*18_*c*_, which is cleaved from its pro-form by caspase 1 and transports out of the cell via gasdermin pores, in the same way as interleukin 1β. That is, the evolution can be described by:

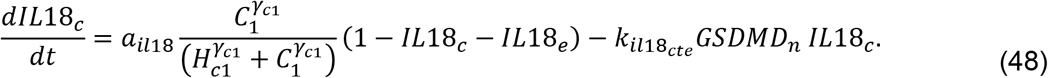

External interleukin 18 levels only depend upon transport out of the cell in the ODE model,

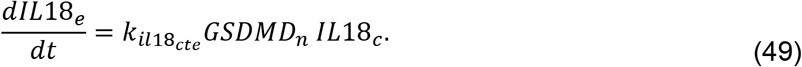

Finally, once gasdermin pores form on the cell surface external material can enter the cell causing the cell to swell. Therefore, we allow the cell volume, *V*, to increase through the equation:

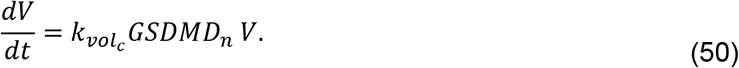

Once the cell volume reaches 1.5 times the original value, then the cell bursts and all cellular processes cease.

The cytokines released by the cell, IL-1β and IL-18 are then modelled in the epithelial environment. We allow IL-1β to potentially initiate pyroptosis in nearby epithelial cells, through a bystander effect^146–148^. Whereas IL-18 acts as a chemoattractant for immune cells, directing them to the local environment^146,147,149–151^. Both cytokines are modelled as a diffusible field.

### Lymph node model

The time course of arrival of the dendritic cells to the LN can be presented as

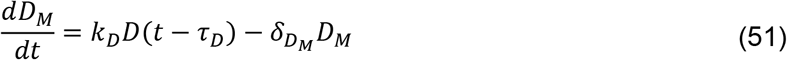

where, *D* presents the number of dendritic cells in the tissue and *D_M_* is that migrated into the LN. *k_D_* and *τ_D_* are the antigen presentation rate by dendritic cells and time taken by the DCs to migrate into the LN. The natural death rate of the *D_M_* is denoted as *δ_D_M__*.

In the LN, the proliferation, activation, and clearance of the two types of helper T cells (*T_H1_* and *T_H2_*) and cytotoxic T cells (*T_C_*) are modelled according to the following ODEs

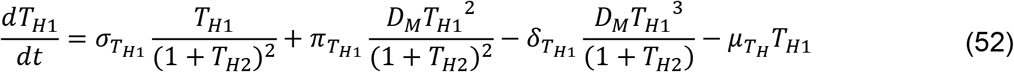

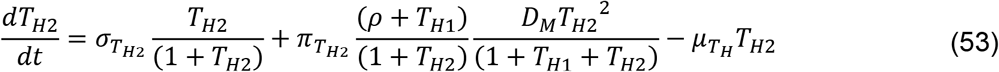

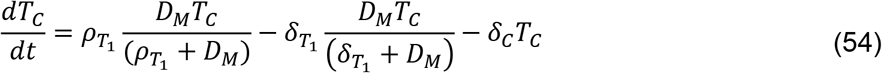

where, *σ_T_H__* and *π_T_H__* denote the proliferation and activation rate constants for type 1 and type 2 helper T cells, respectively, *δ*_*T*_*H*1__ represents the DC-mediated deactivation of type 1 helper T cells, and *μ_T_H__* is the same natural death rate of both cell types. The TCs/CD8^+^ T cells are simultaneously activated and cleared by DCs by two TC population-dependent rates at rates *ρ*_*T*_1__, *ρ*_*T*_2__ and *δ*_*T*_1__, *δ*_*T*_2__.

#### Lymph node additions to Version 5

Version 5 added the plasma cell (P) and B cell (B) species to the lymph node to allow production of antibody, or immunoglobulin (Ig). The B cell activation requires two exclusive signals: antigen presentation by D_M_ and accessory signal from T_H2_. The active B cells differentiate and mature to terminal plasma cells which are the major contributor in antibody production. There are two different hypotheses on the B cell activation and antibody production: (1) activated B-cells start differentiating into plasma cells at a very early stage and produce antibody leading towards early seroconversion; (2) activated B cells keep proliferating until a threshold and then start the differentiation process and push an enormous amount of antibody. This is the case of delayed but sharp sero-conversion. It also introduced a capacity of available Tc cells within the system. We now define the rate of Tc (CD8^+^ cells) in the lymph node as,

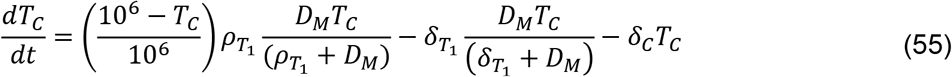

Where we assumed that 10^6^ CD8^+^ T cells are available in the lymph node for activation. We then define B Cell and plasma cell activation as

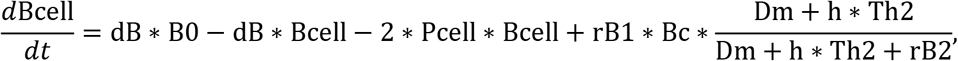

where dB is the basal rate of decay and production such that B0 (considered to be the number of inactive B cells) is the fixed point in the absence of stimulus, rB1 is the rate of DC assisted B Cell proliferation, h the Th2 weight parameter, rB2 denotes the damping parameter, and pSc is the B cell to plasma cell reaction rate, given by

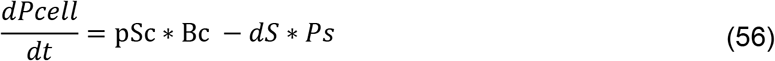

where pSc is still the B cell to plasma cell reaction rate and dS is the plasma cell clearance rate. Antibody dynamics are modelled by

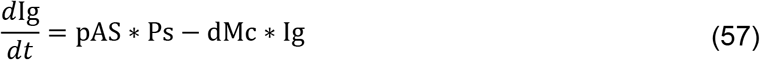

where pAS is the rate of Ig production by plasma cells and dMc is the rate of Ig degradation, for transport of Ig into the tissue see section Immunoglobulin (Ig) recruitment to the tissue.

#### Addition of time delays to the system

Version 5 added transmigration delay for both DCs and CD8^+^/CD4^+^ cells set at 0.5 days and 0.25 days respectively^152,153^. The transmigration of resident DCs from the peripheral tissues to the lymph-nodes occurs via lymphatics. The DCs enter the terminal lymphatics by squeezing through the discrete cell junctions of the lymphatic endothelium. The transit from the terminal lymphatics to the nearest lymph-node of DCs occurs by guided signals and chemical gradient. The T cells crawl on the surface of the endothelium and follow cytokine gradient, which regulate the T cell motility. Time delay was added as a proxy for the transportation time of cells to and from the lymph node.

#### Parameter changes to the Lymph node

Various rates were modified in v5 to match earlier time scale data for CD8 and Antibody rates in healthy mice^154^. This leads to a generally faster arrival rate of cells from the lymphatic compartment.

### Tissue damage additions to Version 5

In version 4, the source of anti-inflammatory cytokine was activation of latent TGF-β. We assumed the latent TGF-β is activated and secreted continuously at the location of dead epithelial cells killed by CD8+ T cells. In version 5, we consider secretion of TGF-β from damaged sites which only occurs for a certain time instead of continuous secretion^155^. We also consider secretion of TGF-β from the macrophage^155–157^. We also assume macrophages start secreting TGF-β when CD8^+^ T cells inhibit the production of pro-inflammatory cytokine from the macrophage. The resident fibroblasts then become activated and recruited by TGF-β. Fibroblasts chemotaxis towards the gradient of anti-inflammatory cytokines and deposit collagen (Fig. 5.1).

**Fig. 5.1:**
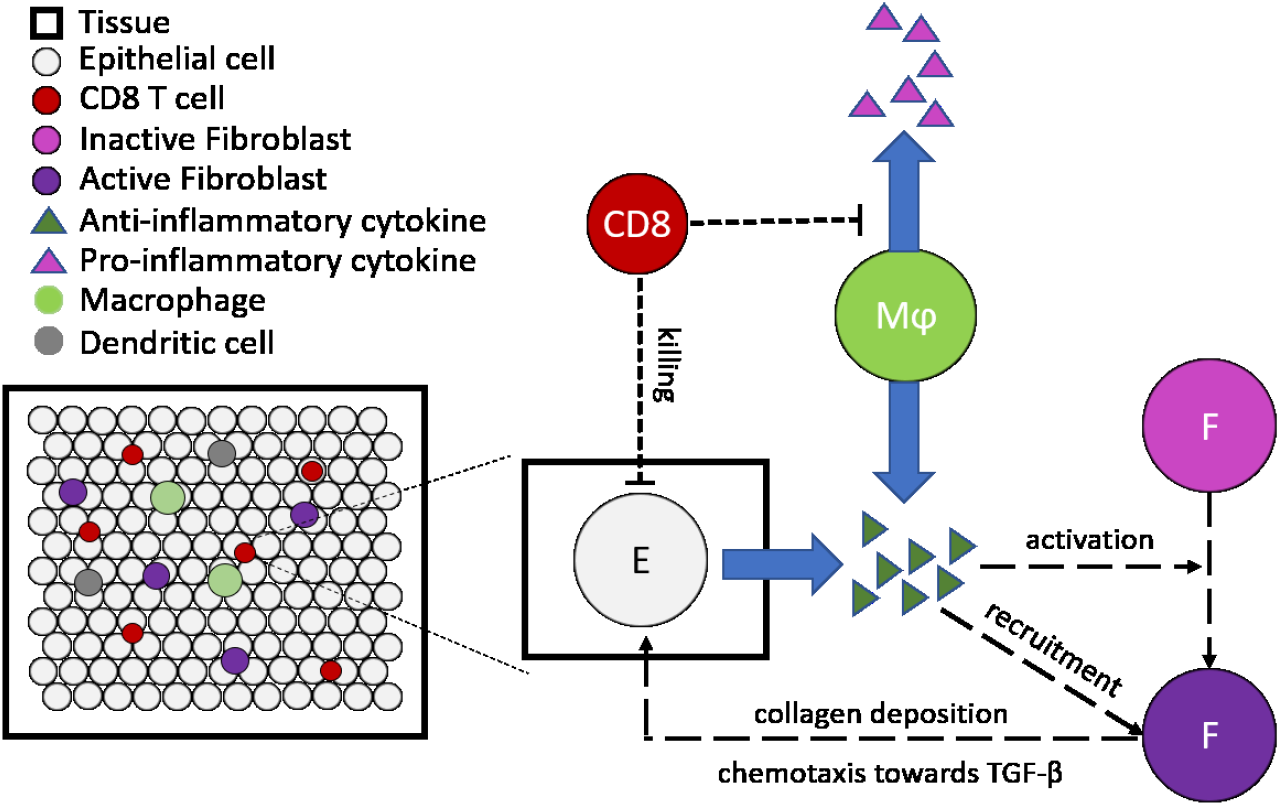
Fibrosis submodel schematic. Latent TGF-β becomes activated at the site and time that a CD8^+^ T cell kills an epithelial cell. Macrophages start secreting TGF-β when CD8^+^ T cells inhibit the production of pro-inflammatory cytokines. Fibroblasts become activated and recruited by TGF-β and deposit collagen.

#### Immune additions to Version 5

##### Immunoglobulin (Ig) dynamics in the tissue

Antibodies are protective proteins produced by the immune system in response to viral infections that assist with the removal of free virions. These antibodies generally have a high degree of specificity for the infecting virus and play an important role in the prevention of future infection with the same virus. In patients with severe COVID-19, antibody production appears to be delayed, but of higher magnitude than in patients with milder disease^158^. In our model, Ig arrives at the infection site from the lymph node. Once in the tissue, Ig diffuses and neutralizes virus through opsonization. We model this by assuming the virus and Ig bind together and are instantly removed from the system^159^. Infected cells can bind Ig, which we assume allows macrophages to identify infected cells and phagocytose them. We assume this occurs irrespective of whether a macrophage is hyperactivated or not.

##### Immunoglobulin (Ig) recruitment to the tissue

Immunoglobulin is recruited by the lymph node system built in the v4 COVID model, leveraging the addition of B cells and plasma cells to the lymph node, see section **Lymph node additions to Version 5**. Once a full Ig has been formed in the lymph node it is then removed and added to the tissue. The position of this added Ig is decided by picking a random voxel within the bounds of the system, as we do not currently consider entry through discrete vasculature points. We then add a single Ig to the local concentration of the continuous field (adding 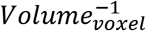). If multiple Ig are recruited on a single time step, then multiple random voxel assignments are performed for every Ig. Ig recruitment events occur on the diffusion time scale.

##### Neutrophil Reactive Oxygen Species (ROS) release and tissue damage

Neutrophils phagocytose induce necrosis in epithelial cells through the release of NETs and other antimicrobial signals such as reactive oxygen species (ROS)^160,161^. Once activated, neutrophils secrete ROS which diffuse through the tissue. ROS causes epithelial (uninfected or infected) cells to undergo apoptosis with probability

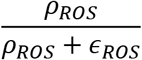

where *ρ_ROS_* is the density of ROS, and *ε_ROS_* is the half-effect of ROS on epithelial cell death.

##### T cell proliferation and death in the tissue

Both CD4+ and CD8+ T cells will proliferate on arrival in the lung^162,163^. We assume that T cells proliferate for a limited number of generations (*T_generation_*) within the lung tissue^162^ and that proliferation is halted, increasing the rate of apoptosis, once a cell has proliferated *T_generation_* times.

#### Other implementation notes

This simplified immune model does not yet include many key immune agents, including natural killer (NK) cells, the complement system, and most cytokines. Anti-inflammatory cytokines are modeled, but this model cannot return to a sufficient homeostasis (a post-infection immune state that is similar to its pre-infection state) following potential infection clearance. Currently, the system will eventually deplete neutrophil, macrophages, and dendritic cells in the absence of pro-inflammatory cytokine recruitment signals. Dynamics of cytokine binding and unbinding to receptors are also omitted. The model does not yet incorporate known SARS-CoV-2 immune evasion techniques, such as a delayed IFN-I response and lymphopenia (decreased CD8^+^ T cells) from early in infection. In addition, the antigen presentation and subsequent T cell activation in the lymph node is not explicitly modeled and instead exists as a simplified ODE model. Many of these important mechanisms are planned for inclusion in future versions. See further discussion in the modeling results below.

#### Software release

The core model associated with the v5 prototype is Version 0.5.0. The nanoHUB app associated with the v5 prototype is Version 5.0. GitHub releases and Zenodo snapshots are given in the Appendix.

The cloud-hosted interactive model can be run at https://nanohub.org/tools/pc4covid19.

#### Model behavior: what does the current version teach us?

Except as noted below, all simulation results use the v5 model default parameters, which are supplied in the XML configuration parameter file of the version 0.5.0 core model repository found at https://github.com/pc4covid19/COVID19/blob/master/PhysiCell/config/PhysiCell_settings.xml.

##### Overview of v5 model results with base parameters

Initial simulations with all the added model features used the v5 base parameter set (i.e., the default configuration file included with the version 0.5.z releases on GitHub), and the results are shown in Figure 5.2. Following the introduction of virus, there is a peak of viral replication that occurs around day 4, followed by rapid clearance of the virus by day 6 (Fig 5.2B, blue line). Interferon (IFN) production (Fig 5.2B, orange line) occurs very rapidly and is maintained for the whole simulation, until cell death causes a decrease. Fig 5.2C shows dynamics of the immune cell populations in the tissue, which illustrates a rapid and transient influx of CD4^+^ T cells followed by a more sustained recruitment of CD8 T^+^ cells that remain in the tissue for the full 15 days.

**Fig. 5.2:**
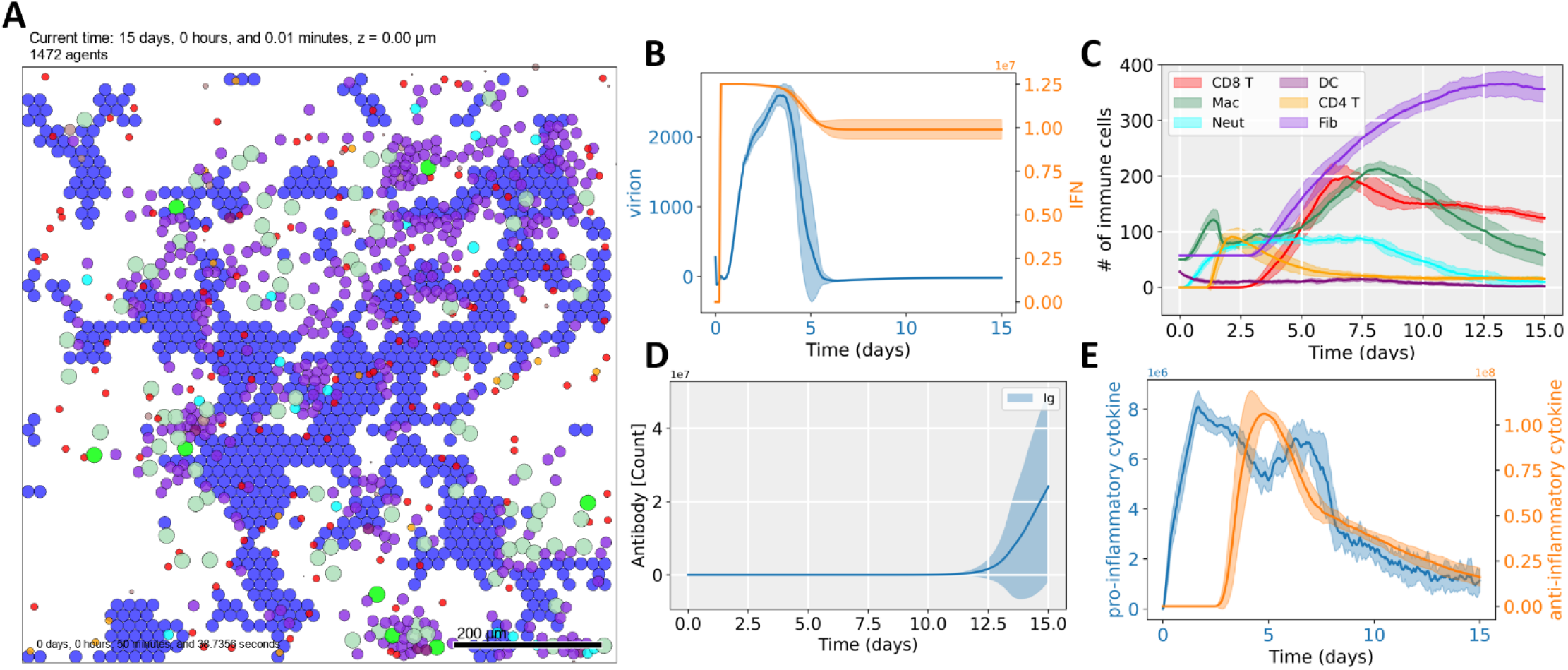
15-day dynamics of the version 5. Results for 12 replicates show removal of virion from the system in ~6 days and subsequent system recovery to the infection. The system sees acceptable tissue loss when compared to previous versions (A). Viral presence is removed within 6 days of infection (B, blue line) coupled by a rapid and sustained IFN presence (B, orange line). Immune cells within the tissue (C). Total Antibody contained in the tissue (D). Pro-inflammatory (E, blue line) and anti-inflammatory (E, orange line) plots across the time course. See supplemental 1 for a localized figure and total collagen plot. For all plots, N=12 average and standard deviation of the related species were plotted.

Antibody recruitment does not provide any major protection to the system for an initial infection as Ig production does not ramp up until post 12 days (Fig 5.2D) and most virion has already cleared. We observe a large standard deviation at late time for Ig, likely due to the rapid and sudden increase in production at later times. Interestingly, due to CD8^+^ T cell contact switching macrophages to M2 phase secreting anti-inflammatory cytokine, we now see the system turn off the pro-inflammatory cytokine signal (Fig 5.2E, blue line) once CD8^+^ T cells flood the system (Fig 5.2C, red line). Further, under the current rates and rule sets we observe two times which we observe decreases in pro-inflammatory secretion. First is the change to M2 phase, noted by the rapid increase in pro-inflammatory cytokine at ~4 days (Fig. 5.2E, orange line). Second is after the mass cell death by viral infection which occurs at ~6 days which is likely due to a combination of removal of cell dependent secretion, macrophage exhaustion, and removal of the activating stimulus (cell death and virion).

The kinetics of collagen deposition (Fig 5.2-supplement 1A) follows the time course of tissue damage and rises rapidly after ~4 days when the infection starts clearing. In Fig. 5.2-supplement 1B, we can see the dynamics of latent TGF-β activation sites as secreting agents rise after CD8^+^ T cell introduction to the system. Fig. 5.2-supplement 1C shows the distribution of collagen deposition at damaged sites, the amount and distribution of collagen deposition vary due to the dynamics of fibroblast cell population and sources of TGF-β. We observed activation of latent TGF-β causes localized collagen deposition whereas secretion of TGF-β from macrophage contributes to more disperse collagen deposition.

**Fig. 5.2-supplement 1:**
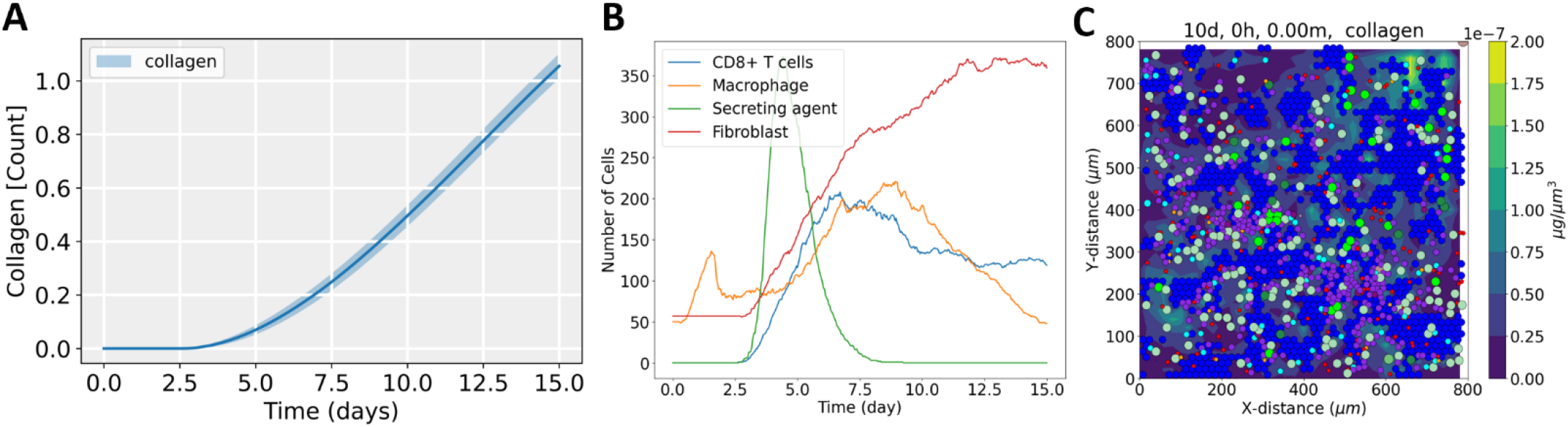
Dynamics of latent TGF-β activation sites (secreting agents), CD8+ T cells, macrophage, fibroblasts, and collagen deposition at TGF-β secretion rate 1 min^-1^ from damaged sites and 0.07 min^-1^ from macrophages. A. population dynamics of CD8+ T cells, macrophage, secreting agent, and fibroblasts, the shaded area represents 5^th^ and 95^th^ percentile of 15 iterations, B. amount and distribution of collagen in tissue at day 15.

##### Impact of a time delay

Since version 5 incorporated time delays for DC and T cells to simulate migration of these cells between the tissue and the LN, we asked what the effect of varying this time delay would have on the system. We therefore ran simulations with 0 delay on any migration, delay on the DC migration from the tissue to the LN (0.5 days), delays on both DC (to the LN) and T cell (from the LN to the tissue) migration (0.5 and 0.25 days respectively, base parameters), and a double delay case of DC (1 day) and T cells (0.5 day). The arrival of CD4 and CD8 T cells are delayed as expected (Fig 5.3A), but these changes do not have a dramatic impact on the dynamics of macrophages, neutrophils and fibroblast recruitment. However, we do see a difference in the cell survival of the system at 15 days (Fig 5.3B): delaying the arrival of T cells results in more destruction of the epithelial layer. This would match expected results, as it has been shown that patients with severe disease have delayed T cell recruitment to the lung. Thus, the dynamics of the T cell response has a major impact on the severity of the disease overall.

**Fig. 5.3:**
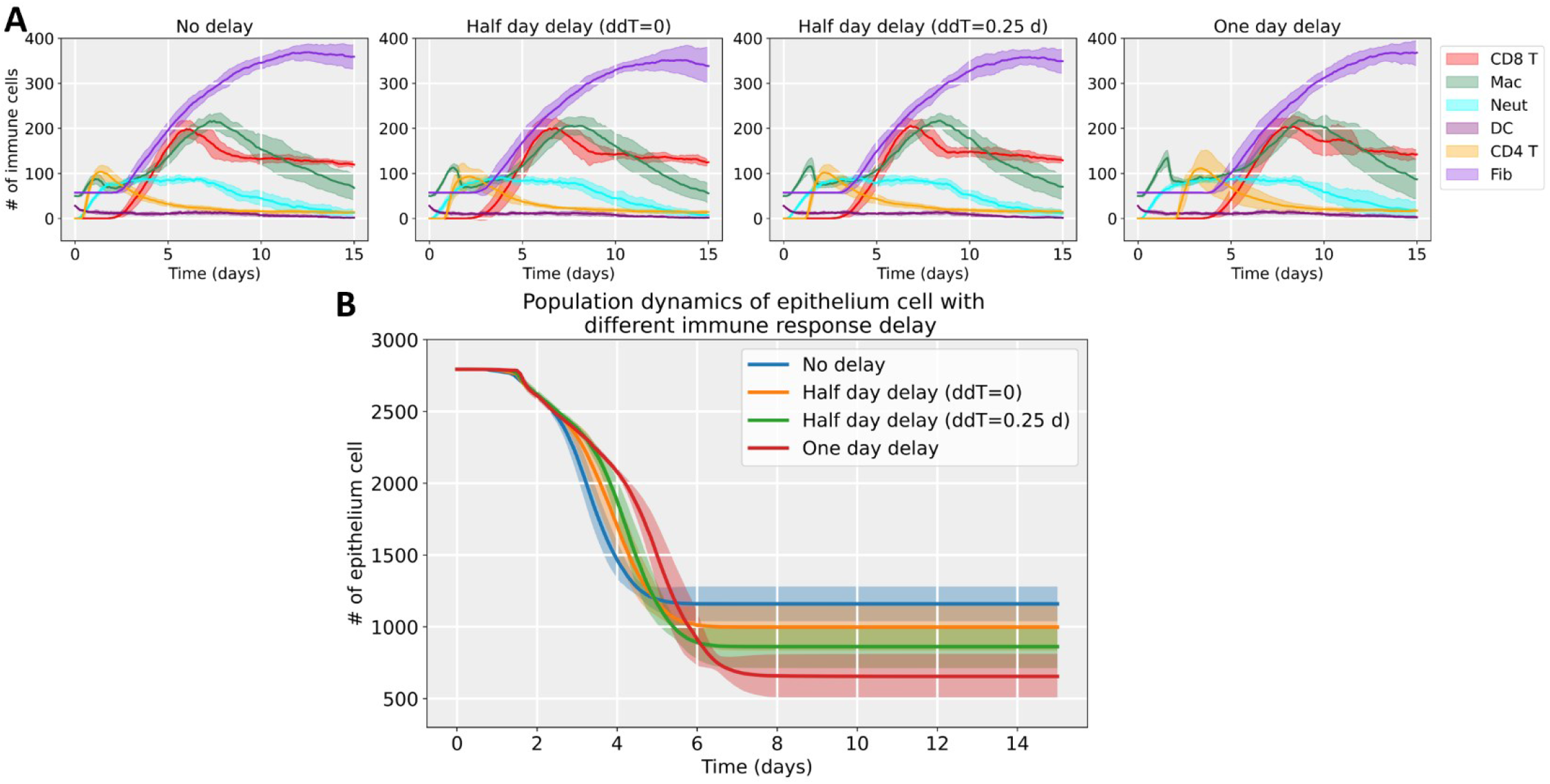
A. Simulations of the COVID-19 tissue simulator with multiple time delay values of none, DC only, both DCs and T cells, and doubled delays on both DC and T cells. B. Survival of the epithelial cells for each scenario is plotted indicated reduced survival as the delays are increased.

##### Impact of prior immune responses on subsequent viral exposure

An advantage of our model is that we can explore the impact of a prior successful response on subsequent viral exposure and infection, as well as the impact of waning immunity. To investigate these scenarios, we used the final simulation results (at *t* = 15 days) to help initialize the initial immune state for new simulations, with a varied level of initial immunity “educated” by the prior infection. Suppose *IC_e_* represents this “educated” immune state (lymph node model status, immune cell counts, and diffusible non-viral factor concentrations) and if *IC_n_* represents the “naïve” immune conditions prior to infection. Then to study the impact a fully or partially educated immune state, we can linearly interpolate these to get the initial immune conditions *IC*(*f*) = (1 - *f*)*IC_n_* + *f IC_e_*, where 0 ≥ *f* ≥ = 1 represents relative level of remaining adaptive immunity from the prior immune response and viral recovery. If *f* = 1, then the prior immunity remains at full strength, *f* = 0 corresponds to conditions where all prior immunity has waned and the system has returned to a naïve state, and intermediate values correspond to intermediate levels of immunity.

Due to large slowdowns when recruiting large numbers of Ig, the *f* = 1 case needs to be capped off at some value to allow rapid solving of the system (1000 per diffusive time step in this case). The system was then challenged with a new infusion of virus and the viral, cellular and tissue dynamics were followed over time (Fig 5.4). In the case where all immune parameters are maintained at the educated level (*f* = 1), there is very little tissue damage (Fig 5.4A). Reducing the value to *f* = 0.03125 (Fig 5.4B) has little impact on tissue damage whereas a reduction to *f* = 0.002 (Fig 5.4C) results in more extensive tissue damage. Examination of the cellular dynamics (Fig 5.4D-E) reveals that the CD8 response is quite different between these scenarios. In the fully immune case (*f* = 1) case CD8^+^ T cells are still present in the tissue at the time of the second viral inoculation (Fig 5.4E), whereas these cells are not present in the two other scenarios. In contrast, CD8^+^ cells are recruited to the tissue about 2 days earlier in the *f* = 0.03125 case (Fig 5.4E) compared to the *f* = 0.002 case (Fig 5.4F) demonstrating again the impact of a delay of the T cell immune response. Examination of the viral load in these scenarios (Fig 5.4G) reveals that viral is rapidly cleared in the two first scenarios, whereas these is a significant increase in the viral load in the *f* = 0.002 case, which approaches that seen in the naïve case. While antibodies were present at the start under all three conditions (Fig 5.4H) the overall level varied between the three cases by at least two orders of magnitude. These results may provide some insight into the levels of antibody that are required for sustained immunity. These results suggest that immunity following an initial infection may be quite robust and that severe disease would only occur once immunity has waned quite dramatically.

**Fig. 5.4:**
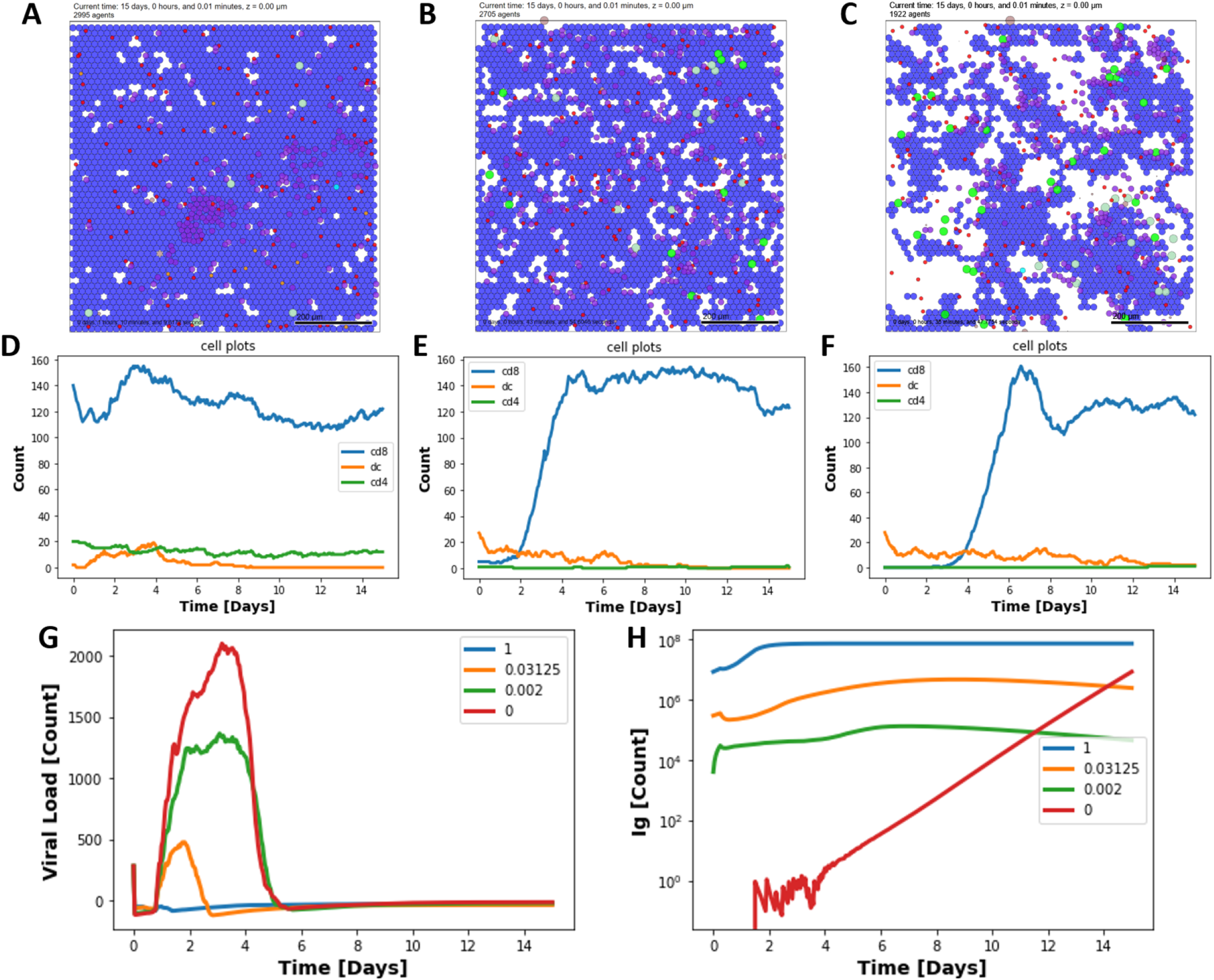
results of multiple immune cases ranging from a very recently infected system (A, *f*=1) to a system that carries substantial (B, *f*=0.03125) or traces of immune memory (C, *f*=0.002). D, E and F show the dynamics of the DC and T cells for the three different scenarios. The viral load (G) and immunoglobulin levels (H) are shown for the three scenarios and include the naïve (*f*=0) case for comparison.

##### Global sensitivity analysis

To better understand model robustness, we performed linear regression sensitivity analysis for the parameters maximum IFN secretion rate via paracrine, viral diffusion coefficient, ACE2 binding rate constant, and T cell recruitment rate constant. We compared these parameters against their impact on the final cell count values (Fig 5.5). Multiple simulations of the v5 model were performed for 15 days with different values for the 4 parameters. Parameter values are generated using Latin hypercube sampling, staying within an order of magnitude of the initial values of each parameter. The linear regression is done by using the log10 of the final cell count and log10 of each parameter and then performing a 4-dimensional linear regression. We then project the 4D vector of parameters onto the coefficient vector of the linear regression comparing against the log10 of the final cell count result from the simulation ran using the parameters to obtain a linear fit to the data. Only samples that ended up having over 0 final cell counts and under the maximum cell counts were used for the fit. The valid parameter space for those values is large and doesn’t help with the sensitivity analysis, these represent the cases where either nothing has died, or everything has died. The fit coefficients obtained show the ACE2 binding constant is the parameter that contributes the most to having higher live cell counts followed by the viral diffusion coefficient and IFN secretion rate. The T cell recruitment rate constant doesn’t seem to affect how many cells are alive by the end of 15 days as the earlier players are much more sensitive. We can then conclude that the viral spreading rate is one of the most important parameters within the system even within the presence of IFN action.

**Fig. 5.5:**
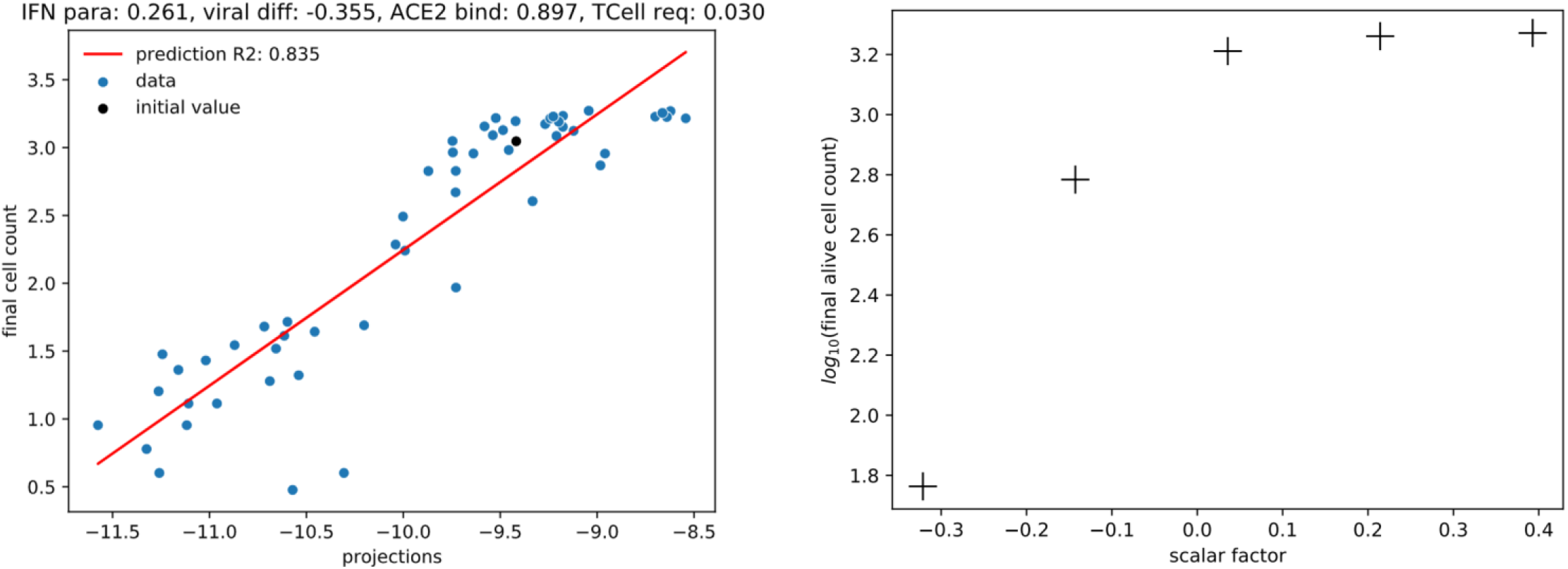
Linear regression sensitivity analysis results for parameters maximum IFN secretion rate via paracrine, viral diffusion coefficient, ACE2 binding rate constant and TCell recruitment rate constant and their impact on the final cell count values. Each point is a simulation of the V5 model for 15 days with different values for the 4 parameters. The x-axis is calculated by projecting the 4D vector of parameters onto the coefficient vector of the linear regression. The y-axis is calculated by taking the log10 of the final cell count result from the simulation ran using the parameters. The red line is the linear fit to the data, blue points are samples and black points are the initial parameter vector. Only samples that ended up having over 0 final cell counts and under the maximum cell counts are used for the fit. The title shows the normalized linear regression coefficient values for each parameter.

### Discussion of current model version

This version of the model has included many new components, including later (longer time scale) immune responses, the introduction of Ig and ROS. In addition, v5 introduced different macrophage activation states (M1/M2) and refinement of fibrosis in response to tissue damage. The addition of these components has introduced many parameters that creates challenges in calibrating the model.

The viral replication still creates an infection that spreads very rapidly. This can result in an inability to clear virus unless aggressive IFN action, high ACE binding rates, or reduced diffusion of virion is introduced. In this absence, even under even the most favorable later-stage immune response conditions, the system will suffer. Localizing the initial viral seeding (as might be expected if inhaled viral particles are not uniformly distributed throughout the respiratory system) may ameliorate these effects. At present virus is introduced at the start of the simulation randomly across the whole tissue at an MOI of 0.1, resulting in widespread virus seeding and rapid spreading before the immune response begins. A more realistic approach may be to seed the virus in a more discrete-localized area, which will be explored in future versions as a special case. In addition, due to require rapid binding we are observing failures of model assumptions that can temporarily create locally negative concentrations. There could be multiple avenues to fix this issue, either though reduction in diffusion rates to allow for reduction in binding rates or though changing the system to a purely discrete handing for the virion particles.

As in prior versions, several parameter estimations remain challenging for Version 5, resulting in particular in overactive IFN. In addition to halting the spread of virion, the relatively high binding rates of ACE2 to virion can temporarily result in negative virion concentration values in the tissue (although total intracellular plus extracellular viral mass is conserved). We postulate that in the absence of a sufficiently slow virion diffusion rate a tissue can more effectively form viral plaques by aggressively uptaking the virion, in essence creating a diffusive buffer in the system. This effect can benefit the immune response, as the immune system can more readily target one localized infection as opposed to multiple patches through the system (i.e a fast diffusive system). The ***global sensitivity analysis*** confirmed that the viral diffusion rate is one of the most important parameters in this model.

This version included time delays in the migration of DC and T cells between the LN and tissue. The impact of varying these time delays revealed a detrimental effect on epithelial cell survival, such that more tissue was damaged as the recruitment of T cells to the tissue was delayed. These delays had little impact on the dynamics of other cell types in the tissue so this result is most likely due to the direct impact of CD8^+^ T cells on viral clearance.

The addition of antibody did not impact the base scenario as antibody levels did not rise until late in the simulation and after the virus was cleared. However, antibody levels did appear to play a role in the response to a second infection. When antibody levels were allowed to wane significantly the response to a second infection was suboptimal resulting in greater viral loads and more tissue damage, The analysis of the the impact of a waning immune response also highlighted the importance of tissue resident CD8 T cells, as well as the rapid infection of a T cell response. One important result from this analysis was that the immune response that is generated following the first infection is quite robust and even if the resposne has waned by more than 30-fold the virus can be cleared with minimal tissue damage.

Data from influenza infections has been used to calibrate this model^154^ in a preliminary sense and will need to be properly expanded to properly inform rates and time scales. This will likely lead to changes to the LN model, if the currently used functional forms are incorrect. For instance, it is known that Th1 cells are necessary for optimal CD8^+^ T cell responses, and the present model does not include this requirement. The model could be refined by adding the requirement that Th1 cells are necessary for the proliferation and expansion of CD8^+^ T cells. In this scenario, activated DCs carrying viral antigen migrate to the LN where they activate CD8 T cells and induce the expansion of Th1 and Th2 cells. Th1 cells interact with CD8^+^ T cells to induce their proliferation and expansion. Th1 and Th2 cells are mutually inhibitory^164^, but in the present model the inhibition of Th2 cells by Th1 cells is not included, which results in the domination of the Th2 response. The Th2 response is necessary for optimal antibody production and thus it will be important to model this accurately. Recent data suggest that patients with severe COVID-19 disease had robust but delayed IgG anti-spike antibody responses ^165^. Thus, the balance and timing between the Th1 and Th2 responses may be critical in determining whether the disease is mild or severe.

### Priorities for Version 6

After extensive discussion of the Version 5 model results, the coalition set priorities for Version 6 development. As after Version 5, all major cell-cell interactions are included, and so Version 6 will focus on virtual experiments to probe the impact of initial immune state, prior immunity, and interactions with treatment. We will refine the source code, algorithms, and parameter estimates to help drive this model exploration. First, there are multiple unintended interactions that exist in the current model that will be addressed in the next version. For instance, the system can see macrophage activation in the absence of viral load through the uptake of debris from random cell death; this can readily be addressed by only stimulating macrophage activation when phagocytosing cells with viral proteins. In addition, it is recognized there is likely missing interactions in the LN ODE model that could improve LN to tissue coupling. This could allow for a more robust LN response and therefore a healthier system. We plan to include a more robust binding process of virion requiring multiple ACE2 receptors for virion uptake.

Finally, we see some non-physiologic behavior in substrates such as IFN (system sustains a response well after infection) and virion (locally negative virion concentrations for brief times). The coalition plans to thoroughly review the overall system to check all features added in versions 1-5 for logical self-consistency and for consistency with known immunology, to prepare for final model release and publication. Version 6 plans to include multiple initial immune conditions to simulate states the tissue may exist (i.e., disease states).

## Discussion

Within three weeks of the World Health Organization’s declaration of a global pandemic of COVID-19^166^, community-based prototyping built upon an existing PhysiCell 3D cell-modeling framework to rapidly develop Version 1 of an intracellular and tissue-level model of SARS-CoV-2^109^. A growing coalition of domain experts from across STEM fields are working together to ensure accuracy and utility of this agent-based model of intracellular, extra-cellular, and multicellular SARS-CoV-2 infection dynamics. Version 1 development underscored the necessity of clearly explaining model components, defining scope, and communicating progress as it occurs for invaluable real-time feedback from collaborators and the broader community. This rapid prototyping helped to grow the coalition and recruit complementary expertise; for instance, a team modeling lymph node dynamics and immune infiltration joined during the Version 1 cycle after seeing initial progress.

The version 1 prototype also showed the scientific benefit of rapid prototyping: even a basic coupling between extracellular virion transport, intracellular replication dynamics, and viral response (apoptosis) showed the direct relationship between the extracellular virion transport rate and the spread of infection in a tissue. More importantly, it showed that for viruses that rapidly create and exocytose new virions, release of additional assembled virions at the time of cell death does not significantly speed the spread of infection. Moreover, decreasing the cell tolerance to viral load does not drastically change the rate at which the infection spreads, but it does accelerate the rate of tissue damage and loss, which could potentially trigger edema and ARDS earlier. This suggests that working to slow apoptosis may help preserve tissue integrity and delay adverse severe respiratory responses. That such a simple model could already point to actionable hypotheses for experimental and clinical investigations points to the value of rapid model iteration and investigation, rather than waiting for a “perfect” model that incorporates all processes with mechanistic molecular-scale detail.

Version 2 showed promise of increasing mechanistic details to evaluate potential inhibitors. For example, it was found that that reducing the expression of ACE2 receptors could paradoxically lead to faster spread of the infection across the tissue, although the individual infected cells would replicate virus more slowly. On the other hand, taking advantage of high receptor expression but interfering with viral release from internalized receptors may help slow infectious dynamics. Generally, adding sufficient actionable cell mechanisms to the model framework allows us to ask pharmacologically driven questions on potential pharmacologic interventions, and how these findings are affected by heterogeneity, stochasticity, and the multiscale interactions in the simulated tissue.

Version 3 allowed our first investigations of immune system responses. We found that T cell behaviors are critical to controlling the spread of an infection through the tissue. In particular, rapid recruitment as well as the presence of “educated” CD8^+^ T cells prior to infection (e.g., after responding to infection in a nearby tissue) had a significant protective effect, even in the current model that does not explicitly model antibodies. This is consistent with emerging studies that link T cell responses to patients with the best recovery^167–169^.

Version 4 added multiscale interaction with the lymphatic system, particularly to allow dendritic cells to present antigens to T cells to drive expansion and immune response, trafficking of dendritic cells and T cells between the local, spatially resolved tissue and the lymphatic system. It also introduced interferon responses, which were found to have a profound impact on the spread of viral plaques. Refined models of infected cell death (pyroptosis) and tissue damage (fibrosis) were also introduced.

Version 5 introduced significant new features, with a focus on antibody responses, anti-inflammatory responses, bystander cell killing due to reactive oxygen species, and other refinements to the immune system model. We were able to use this model to ask questions about the timing of certain immune response components as well as the impact of a waning immune response on the ability to handle a second infection. This type of analysis can be expanded to include vaccination, as well as infection with a related but not identical virus. Notably, the Version 5 model was the first that could fully eliminate a viral infection without complete loss of host epithelial cells.

As work on future versions progresses, teams will work in parallel on submodels to add, parameterize, and test new model components. It will be important to balance the need for new functionality with the requirement for constrained scope, while also balancing the importance of model validation with timely dissemination of results. Thus, this preprint will continue to be updated with every development cycle to invite feedback and community contributions.

As of September 2021, we anticipate that Version 6 will fully transition us to Phase III (widespread community use), with a focus on code hardening, documentation, training materials, and improved parameter estimates based on community-wide data sharing.

### Getting involved

To get involved, we welcome biological expertise, especially related to model assumptions, hypotheses, infection dynamics, and interpretation of results. Mathematical contributions to the underlying model or model analysis and data contributions for crafting, parameterizing, and validating model predictions are particularly sought.

We encourage the community to test the web-hosted hosted model at https://nanohub.org/tools/pc4COVID-19. This model will be frequently updated to reflect progress, allowing the public to take advantage of this rapid prototyping effort.

We avidly encourage the community to test the model, offer feedback, and join our growing coalition via Google survey (https://forms.gle/12vmLR7aiMTHoD5YA), by direct messaging Paul Macklin on Twitter (@MathCancer), or by joining the pc4COVID-19 Slack workspace (invitation link). Updates will frequently be disseminated on social media by Paul Macklin (@MathCancer), the PhysiCell project (@PhysiCell), the Society for Mathematical Biology subgroup for Immunobiology and Infection Subgroup (@smb_imin), and others.

We also encourage developers to watch the pc4COVID-19 GitHub organization and to contribute bug reports and software patches to the corresponding (sub)model repositories. See https://github.com/pc4COVID-19

We are encouraged by the fast recognition of the computational and infectious disease communities that we can make rapid progress against COVID-19 if we pool our expertise and resources. Together, we can make a difference in understanding viral dynamics and suggesting treatment strategies to slow infection, improve immune response, and minimize or prevent adverse immune responses. We note that this work will not only help us address SARS-CoV-2 but will also provide a framework for readiness for future emerging pathogens.

## Acknowledgements

PM thanks the Jayne Koskinas Ted Giovanis Foundation for Health and Policy for generous support. PM, RH, and YW thank the National Institutes of Health (U01-CA232137-01) for support. PM, RH, JAG, YW, and JFG thank the National Science Foundation for funding and resources via the nanoBIO Node for nanoHUB (1720625). AS thanks the NIH for support from the NIAID (R01 AI139088). MC, ALJ, and Sofia Alfonso were supported by NSERC Discovery Grant RGPIN-2018-04546 and NSERC ALLRP 554923 – 20. ANFV acknowledges support from the NIH (R35-GM133763). We thank the NCN CP for fast-tracked deployment of models on nanoHUB.

We thank the scientific community for model feedback, including Simon Parkinson, Richard Allen (Pfizer Inc.), David Dai (Pfizer Inc.), Rohit Rao (Pfizer Inc.), and the co-authors of this manuscript. We thank Furkan Kurtoglu (Indiana University) for contributions to SBML integration efforts and other multiscale design aspects. We thank Mark Chaplain for his assistance in coordinating with the UK Royal Society’s RAMP Initiative.

All the authors dedicate this work in memory of Bing Liu, our co-author, colleague, and friend. His insights and community-minded contributions are sorely missed.

## Appendices

## Appendix 1: Code availability

All code is being made available as open source under the standard 3-Clause BSD license. Users should cite this preprint (or the final published paper, as the case may be).

### Core model releases

#### Version 1 model

Version 0.1.0 (released March 26, 2020)

**GitHub:** https://github.com/pc4covid19/COVID19/releases/tag/0.1.0

**Notes:** First release.

Version 0.1.1 (released March 26, 2020)

**GitHub:** https://github.com/pc4covid19/COVID19/releases/tag/0.1.1

**Notes:** Minor bugfixes and first inclusion of “math” directory.

Version 0.1.2 (released March 26, 2020)

**GitHub:** https://github.com/pc4covid19/COVID19/releases/tag/0.1.2

**Zenodo:** https://doi.org/10.5281/zenodo.3733336

**Notes:** First release with Zenodo integration. Last release in 0.1.x chain (v1 model chain).

Version 0.1.3 (released April 1, 2020)

**GitHub:** https://github.com/pc4covid19/COVID19/releases/tag/0.1.3

**Zenodo:** https://doi.org/10.5281/zenodo.3737166

**Notes:** First release after transferring the COVID-19 tissue-level model (overall model) from Paul Macklin’s personal GitHub account to the new pc4COVID-19 GitHub organization.

#### Version 2 model

Version 0.2.0 (released April 9, 2020)

**GitHub:** https://github.com/pc4covid19/COVID19/releases/tag/0.2.0

**Zenodo:** https://doi.org/10.5281/zenodo.3747011

**Notes:** First v2 prototype. Introduced modular design and ACE2 receptor trafficking

Version 0.2.1 (released April 10, 2020)

**GitHub:** https://github.com/pc4covid19/COVID19/releases/tag/0.2.1

**Zenodo:** https://doi.org/10.5281/zenodo.3747011

**Notes:** Minor bugfixes for cell visualization.

#### Version 3 model

Version 0.3.0 (released July 3, 2020)

**GitHub:** https://github.com/pc4covid19/COVID19/releases/tag/0.3.0

**Zenodo:** http://doi.org/10.5281/zenodo.3929320

**Notes:** First v3 prototype. First integration of new immune submodel. Upgrade to PhysiCell Version 1.7.1, allowing use of XML-based cell definitions to define the behavior of immune cell types. Upgrade to PhysiCell Version 1.7.2 beta to improve multithreaded performance, add new cell-cell interaction features, and fix concurrency issues on some platforms.

Version 0.3.1 (released July 3, 2020)

**GitHub:** https://github.com/pc4covid19/COVID19/releases/tag/0.3.1

**Zenodo:** http://doi.org/10.5281/zenodo.3929726

**Notes:** This release improves parameter estimates for digestion of phagocytosed material and has an immune model refinement to prevent runaway macrophage death.

Version 0.3.2 (released July 15, 2020)

**GitHub:** https://github.com/pc4covid19/COVID19/releases/tag/0.3.2

**Zenodo:** https://dx.doi.org/10.5281/zenodo.3946820

**Notes:** This release simplifies the macrophage rules.

#### Version 4 model

Version 0.4.0 (released Nov 20, 2020)

**GitHub:** https://github.com/pc4covid19/COVID19/releases/tag/0.4.0

**Zenodo:** https://dx.doi.org/10.5281/zenodo.4282875

**Notes:** v4 prototype

#### Version 5 model

Version 0.5.0 (released October 19, 2021)

**GitHub:** https://github.com/pc4covid19/COVID19/releases/tag/0.5.0

**Zenodo:** https://doi.org/10.5281/zenodo.5580731

**Notes:** v5 prototype

### nanoHUB cloud-hosted model releases

The latest version can always be accessed directly at https://nanohub.org/tools/pc4COVID-19

#### Version 1 model

Version 1.0 (released March 26, 2020):

**GitHub:** https://github.com/rheiland/pc4covid19/releases/tag/v1.0

**Zenodo:** http://doi.org/10.5281/zenodo.3733276

**nanoHUB DOI:** http://dx.doi.org/doi:10.21981/19BB-HM69

**Notes:** First published version.

#### Version 2 model

Version 2.1 (released April 14, 2020):

**GitHub:** https://github.com/pc4covid19/pc4covid19/releases/tag/v2.1

**Zenodo:** http://doi.org/10.5281/zenodo.3766879

**nanoHUB DOI:** http://dx.doi.org/doi:10.21981/2B1H-GX51

**Notes:** Second published version. Moved GitHub repository to the pc4COVID-19 GitHub organization. Added another tab in the GUI for generating animation of cells (from SVG output files).

#### Version 3 model

Version 3.0 (released July 3, 2020):

**GitHub:** https://github.com/pc4covid19/pc4covid19/releases/tag/v3.0

**Zenodo:** http://doi.org/10.5281/zenodo.3929539

**nanoHUB DOI:** http://dx.doi.org/doi:10.21981/V52J-0S03

**Notes:** The major change to the GUI in this release is the addition of a “Cell Types” tab. This allows editing parameters associated with <cell_definitions> in the configuration file. This version also includes a <style> block in the Jupyter notebook that fixed an unwanted scrollbar in the lengthy About tab.

Version 3.1 (released July 3, 2020):

**GitHub:** https://github.com/pc4covid19/pc4covid19/releases/tag/v3.1

**Zenodo:** http://doi.org/10.5281/zenodo.3929960

**nanoHUB DOI:** http://dx.doi.org/doi:10.21981/YNWZ-GE50

**Notes:** Minor updates to “About” text, e.g., explaining nature of stochastic results. Edits to immune_submod-els.cpp (see details in the core model repository).

Version 3.2 (released July 21, 2020):

**GitHub:** https://github.com/pc4covid19/pc4covid19/releases/tag/v3.2

**Zenodo:** http://doi.org/10.5281/zenodo.3954019

**nanoHUB DOI:** http://dx.doi.org/doi:10.21981/843E-JE78

**Notes:** Update to use core model 0.3.2

#### Version 4 model

Version 4.0 (released November 20, 2020):

**GitHub:** https://github.com/pc4covid19/pc4covid19/releases/tag/v4.0

**Zenodo:** https://doi.org/10.5281/zenodo.4283005

**nanoHUB DOI:** http://dx.doi.org/doi:10.21981/8T5J-9G97

**Notes:** Update to use core model 0.4.0

Version 4.1 (released November 25, 2020):

**GitHub:** https://github.com/pc4covid19/pc4covid19/releases/tag/v4.1

**Zenodo:** https://doi.org/10.5281/zenodo.4290732

**nanoHUB DOI:** http://dx.doi.org/doi:10.21981/M5GC-6E79

**Notes:** Bug fix: Cell Types custom_data were not setting/getting values to/from XML.

Version 4.2 (released January 20, 2021):

**GitHub:** https://github.com/pc4covid19/pc4covid19/releases/tag/v4.2

**Zenodo:** https://doi.org/10.5281/zenodo.4453795

**nanoHUB DOI:** http://dx.doi.org/doi:10.21981/2WY0-SX97

**Notes:** Bug fix: Selectable colormaps and fixed ranges.

#### Version 5 model

Version 5.0 (released November 3, 2021):

**GitHub:** https://github.com/pc4covid19/pc4covid19/releases/tag/v5.0

**Zenodo:** https://doi.org/10.5281/zenodo.5644308

**nanoHUB DOI:** http://dx.doi.org/doi:10.21981/J2AG-5F18

**Notes:** Update to use core model 0.5.0. Outputs at 180 mins. Analysis plots now uses “days” on x-axis.

## Appendix 2: Organoid platform details

Aarthi Narayanan’s virology lab is optimizing SARS-CoV-2 cultures in organoid model systems. The viral replication kinetics will be assessed by infection of different lung epithelial, fibroblast and endothelial cells, in addition to standard cell lines such as Vero cells, which are one of the cell lines in use for inhibitor assessment studies. Primary cells and/or cell lines will be infected with SARS-CoV-2 at increasing multiplicities of infection (MOI) and infectious viral titers in the supernatants assessed by plaque assays at multiple time points post infection (pi). This will stretch from ~3 hours post infection (phi) to 48 hpi depending on the cell type and the MOI.

In parallel, the viral genomic copy numbers will be assessed in the same supernatant samples by qRT-PCR with virus specific primers. This will provide information on how the production of infectious virions compares with the number of genomic copies available outside the cell. If the numbers are skewed in the direction of genomic copies (which may happen in the context of some kinds of inhibitors), it will shed light on the mechanisms of inhibition involving inhibition of infectivity of progeny virions.

The viral genomic copy numbers inside the cells will also be assessed by qRT-PCR and compared to the genomic copies outside the cell. This will provide direction on the efficacy of particle packaging and the extent of production of infectious versus noninfectious virus. While it will not provide directly pertinent information about the possibility of heterogeneity of released virus populations and quasispecies, it can provide initial clues in that direction, which can then trigger more specific questions and relevant approaches. These approaches will be pursued for cell lines, primary cells and, hopefully, subsequently transitioned to organoid platforms.

From a host response point of view, we will pursue two aspects: host cell death and inflammatory responses. For cell survival and death measurements, we will employ an assay that measures ATP activity in cells (hence a reflection of a live cell) in the context of infection and inhibitor treatments. For inflammatory responses, we will assess supernatants for inflammatory mediators by ELISA (multiplexed). The cells will be lysed to obtain RNA, which will be queried for transcription of several genes associated with inflammatory responses using gene expression arrays (multiplexed).

Additional host response events will include mitochondrial activity and ROS production assessments in the context of infection and inhibitor treatments. The impact of anti-inflammatory strategies on mitochondrial activity and cell survival will be assessed to determine correlations between viral replication dependent and independent events.

## Appendix 3: Overall design cycle development details

In each prototyping or design cycle:

1. The core team sets priorities for the design iteration:

a. Discuss feedback and identify highest priority model refinements.
b. Collaborate to update the submodel design documents to address feedback.
c. Update the overall model design document as needed.
d. Assess new data to refine parameter estimates.
e. Refine submodel input/output formats as necessary.
f. Assess next release dates for the submodels.
2. Submodel teams meet to refine their code and put out their next releases. The chief scientists communicate releases to the overall leads.
3. The integration team integrates the latest submodel releases into a new release candidate for the overall model.

a. Address any bug reports.
b. Test new or altered functions.
c. Satisfy all qualitative and/or quantitative unit tests.
d. Qualitatively test the model for new or improved behaviors over the last iteration.
4. The integration team prepares a software release:

a. Update documentation.
b. Create a new numbered release on GitHub.
c. Update list of available validation data and best parameter estimates.
d. Create a Zenodo snapshot.
e. Announce on Twitter (via @PhysiCell, @MathCancer, and @SMB_imin).
5. The integration team updates the cloud-hosted model for multidisciplinary testing:

a. Update the nanoHUB app repository with new code.
b. Run xml2jupyter to update the Jupyter interface.
c. Update project on nanoHUB, test/refine until successful release.
d. Update documentation, numbered GitHub release, Zenodo snapshot of deployed model.
e. Perform live demos with domain experts and community to gather feedback.
6. The whole team seeks additional community feedback via Twitter and the pc4COVID-19 Slack workspace. The team integrates comments received from scientific peer review as appropriate.
7. The core team evaluates progress:

a. Distill feedback to assess the need for new model hypotheses, behaviors, or components.
b. Assess which biological behaviors are currently exhibited by the model.
c. Refine the design protocol (e.g., with refined model specification methods) as necessary.
d. Assess the need for an additional design iteration.
8. Update preprint for scientific dissemination. Return to Step 1 if there is substantial feedback, or if the core team determines that further refinements are within project scope.

## Appendix 4: Submodel development details

In each software sprint, each submodel team will

1. Set priorities for the design iteration:

a. Discuss feedback and identify highest priority model refinements.
b. Refine model assumptions and hypotheses.
c. Assess new data to refine parameter estimates.
2. “Translate” biological hypotheses into agent model rules and other mathematical model components:

a. Run the new hypotheses and rules by domain experts as their time permits.
b. Define new qualitative and/or quantitative unit tests for new behaviors and functions.
c. Assign implementation tasks.
3. Perform computational implementation of refined mathematical model (and submodels):

a. Address any bug reports.
b. Add or modify functions based on new rules in steps 1-2.
c. Test new or altered functions.
d. Satisfy all qualitative and/or quantitative unit tests.
e. Qualitatively test the model for new or improved behaviors over the last iteration.
4. Create a software release:

a. Update documentation.
b. Create a new numbered release on GitHub.
c. Update list of available validation data and best parameter estimates.
d. Create a Zenodo snapshot.
e. Communicate with the core team on the software release.
5. Create a cloud-hosted submodel for multidisciplinary testing:

a. Update the nanoHUB app repository with new code.
b. Run xml2jupyter to update the Jupyter interface.
c. Update project on nanoHUB, test/refine until successful release.
d. Update documentation, numbered GitHub release, and Zenodo snapshot of deployed model.
e. Perform live demos with the core team as needed.

While waiting for the start of the next software sprint, each submodel team will

1. Perform model evaluation:

a. Distill feedback to assess the need for new model hypotheses, behaviors, or components.
b. Assess which biological behaviors are currently exhibited by the model.
c. Refine the design protocol (e.g., with refined model specification methods) as necessary.
d. Assess the need for an additional design iteration.
2. Help update the preprint for scientific dissemination.

In the next software sprint, return to Step 1 if there is substantial feedback, or if the core team determines that further refinements are within project scope.

