## Supplementary Materials for "Iterative community-driven development of a SARS-CoV-2 tissue simulator"

### Supplementary material for Iterative community-driven development of a SARS-CoV-2 tissue simulator

### Previous Version Results

#### Version 1 (March 25-March 31, 2020)

Version 1 was designed as proof of concept rapid prototype to capture essential (but highly simplified) elements of viral endocytosis, protein synthesis, viral assembly, release, and diffusion to infect other cells. The model was tailored to RNA viruses on a tissue monolayer (modeled as a layer of epithelium over a basement membrane). This version was kept deliberately simple to create an early starting framework to help coalesce community feedback and contributions. It was also designed to test the use of interactive cloud-hosted models to help accelerate feedback by virologists and other domain experts through live demos.

The proof of concept model was created by the overall leads (Macklin, Heiland, Wang) while assembling the modeling coalition as an initial starting point and feasibility test for rapid prototyping. Feedback on this version drove the formulation of the design protocols reported above.

##### Submodels

The Version 1 model includes the following submodel components:

- **T:** tissue (which contains epithelial and other cells)
- **V:** viral endocytosis, replication, and exocytosis responses
- **VR:** cell response to viral replication, including cell death and IFN synthesis
- **E:** epithelial cell (incorporates V and VR).

The overall model components are summarized in **Fig 1.1**.

##### Biological hypotheses

In this proof of concept prototype, we modeled a simplified set of biological hypotheses:

1.T.1 Virus diffuses in the microenvironment with low diffusion coefficient
1.T.2 Virus adhesion to a cell stops its diffusion (acts as an uptake term)

1.V.1 Adhered virus undergoes endocytosis and then becomes uncoated
1.V.2 Uncoated virus (viral contents) lead to release of functioning RNA
1.V.3 RNA creates protein at a constant rate, unless it is degraded
1.V.4 Protein is transformed to an assembled virus state
1.V.5 Assembled virus is released by the cell

1.VR.1 As a proxy for viral disruption of the cell, the probability of cell death increases with the total number of assembled virions
1.VR.2 Apoptosed cells lyse and release some or all of their contents

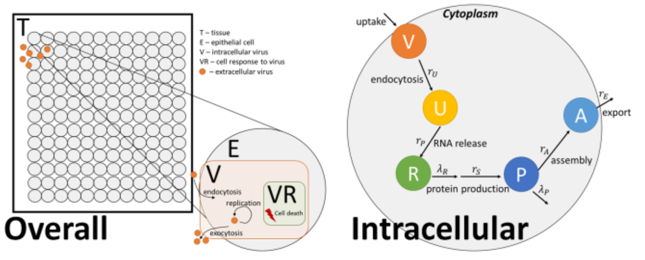


**Fig. 1.1: Version 1 model schematic:** ***Left:*** In the overall model structure, a ***tissue component*** **(T)** contains multiple ***epithelial cells*** **(E)**. Viral particles diffuse through this domain and can enter the cells (by adhering as an uptake term) or be exported by cells. Each cell includes an intracellular ***virus*** model **(V)** for viral replication kinetics and a ***viral response*** model **(VR)** which uses a pharmacodynamic model to control cell apoptotic response to viral load. ***Right:*** Each epithelial cell individually models its viral kinetics: adhered virions complete endocytosis, are uncoated, functionalize their RNA, synthesize viral components, and assemble them into virions that are subsequently exported to the external tissue compartment (T).

(In the above, X.C.Y denotes prototype X, model component C, biological hypothesis Y, allowing us to easily refer to any individual hypothesis or assumption in discussion and community feedback.) In the next version of this model, we will use the design document protocols for each of these components.

##### Unit tests

The first prototype should demonstrate the following behaviors for a single cell infected by a single virion:

- The virion progresses to the uncoated state.
- The uncoated virion progresses to the RNA state.
- With export and death off, RNA produces protein.
- With export and death turned off, protein produces and accumulates assembled virus (linearly).
- With export off and death on, cell undergoes apoptosis with increasing likelihood as assembled virus accumulates.
- With export on and death on, surrounding cells get infected and create virion.
- Cells nearest the initial cell are infected first.
- Apoptosis is most frequent nearest to the initial infected cell.

#### Translation to mathematics, rules, and model components

#### Other implementation notes

To differentiate between incoming imported and exported virions within the computational implementation, we modeled two diffusing fields (for extracellular concentrations of $V$ and $A$). However, the models only require extracellular $V$. At the end of each computational step (advancing by one diffusional time step), we iterate through each voxel and transfer all of the extracellular diffusing $A$ to $V$. We also created diffusing fields for uncoated virions, RNA, and viral proteins, although these were removed from later model versions.

#### Software release

The core model associated with the v1 prototype is Version 0.1.3. The nanoHUB app associated with the v1 prototype is Version 1.0. GitHub releases and Zenodo snapshots are given in the Appendix.

#### Cloud-hosted model

We rapidly created and deployed a cloud-hosted model with an interactive web-based GUI running on nanoHUB (nanohub.org) using xml2jupyter Version 1.1[1]. The web-hosted model can be run at:

https://nanohub.org/tools/pc4COVID-19.

This workflow uses a Python script that converts a PhysiCell configuration file (in XML) into a Jupyter notebook and adds additional Python modules for the GUI. The automated process of converting a standalone PhysiCell model into an interactive Jupyter notebook version (a GUI) takes just a few minutes. The resulting GitHub repository is shared with the nanoHUB system administrators who install it for testing as an online, executable model (an "app"). After we perform usability and other testing and finalize documentation, it is published and becomes available for public use. The whole process (including the initial development of the core PhysiCell model) took less than 12 hours for the Version 1 GUI on nanoHUB (**Fig. 1.2**).

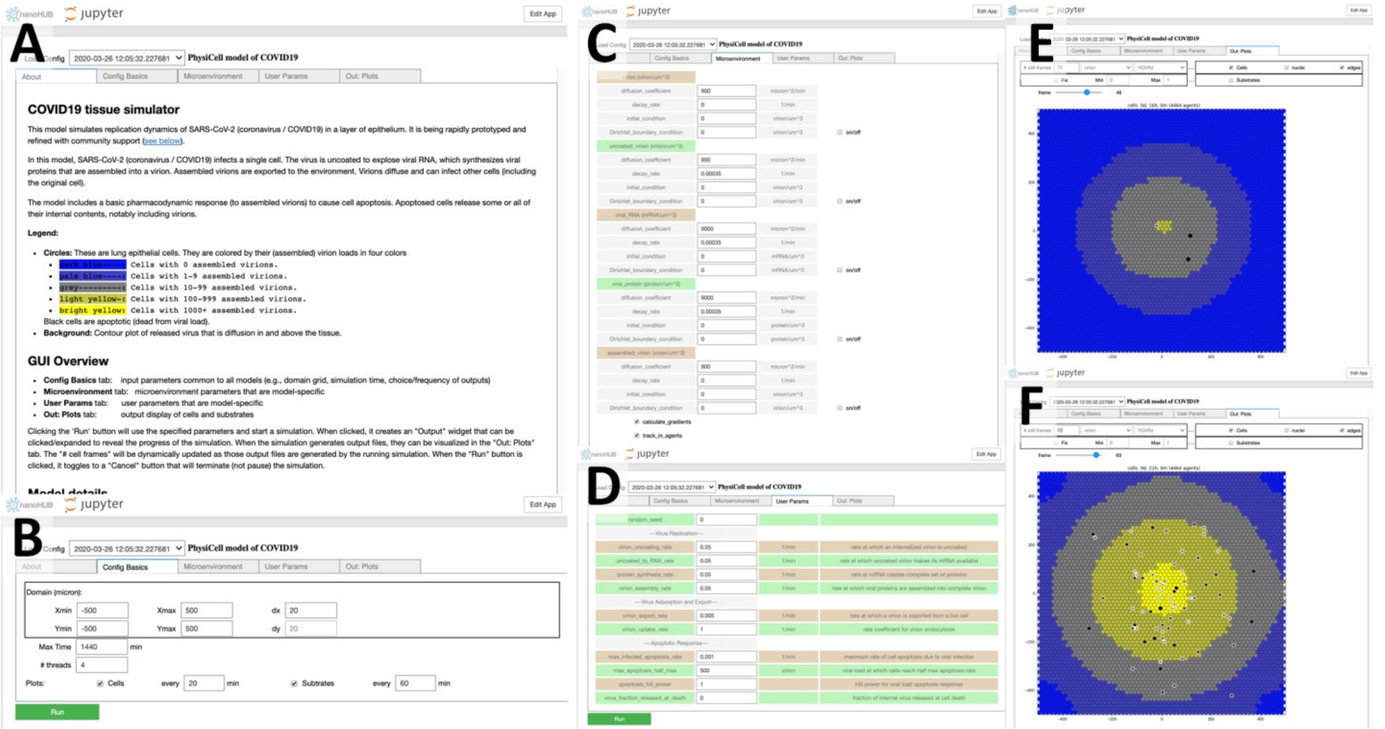


**Fig. 1.2: Version 1 on nanoHUB: A.** The *About* tab includes background information on the model, run directions, and a legend. **B.** The *Config basics* tab allows users to set the domain size. (These should be not be edited in this version.) **C.** The *Microenvironment* tab lets users set diffusion and decay coefficients, as well as initial and boundary conditions. The track_in_agents option must be enabled for the model to work correctly. **D.** The *User Params* tab sets key model parameters. **E-F.** Sample outputs in the *Out: Plots* tab. Users can scroll the "frame" tab to advance the simulation time and choose whether to plot the cells, a diffusing substrate, or both. These plots display a sheet of epithelium (circles are individual cells). Brighter yellow cells contain more assembled virion. Black cells are apoptotic, and clear spaces show tissue damage (where cells have apoptosed and have been removed.) This version of the nanoHUB model was released at DOI 10.21981/19BB-HM69.

#### Model behavior: what does the current version teach us?

Except as noted below, all simulation results use the v1 model default parameters, which are supplied in the XML configuration parameter file of the version 0.1.2 core model repository.

In all plots, dark blue cells have 0 assembled virus, pale blue cells have 1-9 assembled virions, grey cells have 10-99 assembled virions, light yellow cells have 100-999 assembled virions, and bright yellow cells contain 1000 or more assembled virions. Black cells are apoptotic, and white spaces show regions devoid of cells (extensive tissue damage). See the legend in **Fig. 1.2** (A) and the caption in **Fig. 1.3** (A).

##### Behavior with default parameters

Running the overall model (with virus release turned on and off as appropriate for the respective unit tests) shows that the v1 prototype satisfies all the qualitative unit tests. A single cell is infected with a virion in the center of the tissue. Over time, the virion is uncoated to create functionalized RNA, which is synthesized to viral proteins and assembled to functional virus. The graphical output shows this center cell turning to a bright yellow as assembled virions accumulate. By enabling the substrate plot, we can see the diffusive field of virions first has zero concentration (no virions have been released), but as the first cell's viral production increases, it releases virus particles that begin diffusing into the domain (**Fig. 1.3 A**).

Over time, neighboring cells also become infected and progress towards a higher viral load (increasingly bright shades of yellow). The infection propagates outward from the initially infected cell into the remaining tissue. As each cell's viral load (here measured as number of assembled virions) increases, the viral response model calculates the increasing effect *e*, and cells have greater probability of apoptosis. Cells nearest to the initial site of infection apoptose earliest. As these cells degrade, they are removed from the simulation, leading to the creation

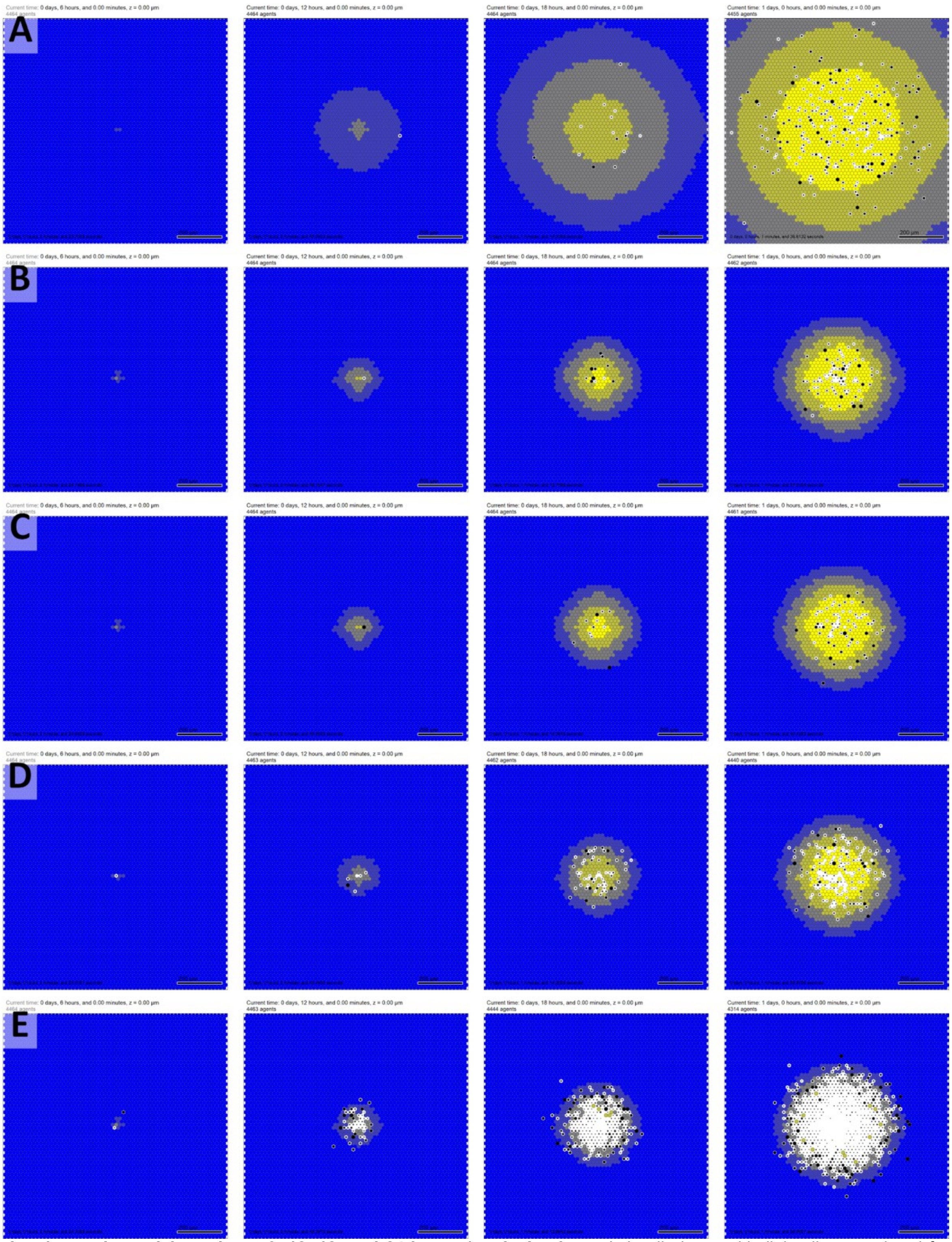


**Fig. 1.3: Version 1 sample model results at 6, 12, 18, and 24 hours (vertical columns).** In all plots, epithelial cells are colored from blue (no assembled virions) to bright yellow (1000 or more virions). Black cells are apoptotic, and white regions show damaged tissues where apoptotic cells have degraded to expose (unmodeled) basement membrane. Bar: 200 $\mu$m. **A.** Simulation time course for the default parameters. Note the spread of the infection from an initial infected cell at the center, with apoptotic death events focused near the center. **B.** Decreasing the diffusion coefficient of virions by a factor of 10 drastically reduces the rate of spread, although focusing exocytosed virions in a smaller diffusion distance increases the number of virions infecting nearby cells, leading to faster apoptosis. **C.** Allowing apoptosed cells to release their assembled virions at lysis had a negligible effect for these parameters, given the dominant effects of releasing virions throughout the cell survival times. **D.** Decreasing the tolerance (half max) of cells to assembled virions prior to apoptosis accelerates tissue damage but does not drastically accelerate the spread of the infection. **E.** Increasing the apoptosis rate (or decreasing the survival time) for infected cells drastically increases tissue degradation.

of a degraded, cell-free region near the center of the tissue. This degraded region spreads outwards from the initial site of infection over time.

See **Fig. 1.3 A** for a simulation with default parameters. The nanoHUB distribution of this model takes approximately 60-90 seconds to execute.

##### Impact of the virion diffusion coefficient

We tested the effect of the viral diffusion coefficient by reducing it from 900 $\mu m^2$/min to 90 $\mu m^2$/min. Because the viral particles spread more slowly after their release, the overall spread of the infection is slowed (**Fig. 1.3 B**). We left $D$ = 90 $\mu m^2$/min for all subsequent investigations of the v1 model.

##### Impact of the viral release at cell death

We tested the effect of releasing all assembled viral particles at the time of cell death by setting $f_{release}$ = 1. For this set of model parameters, the release of assembled virions had a negligent impact of the overall spread of infection: Compare the final frame of **Fig. 1.3 B** (no release: $f_{release}$ = 0) to **Fig. 1.3 C** (complete release: $f_{release}$ = 1). This is because cells release far more virions during their infected lifetimes, so the effect is dominant over the one-time release of virions at cell death. We expect this behavior would change if the cells exocytosed virions more slowly.

##### Impact of the cell tolerance to viral load

We decreased the cell tolerance to viral load by decreasing the $A_H$ of Equation **Error! Reference source not found.** from 500 virions to 10, while leaving $f_{release}$ = 1. As expected, cell death and tissue damage occurred much more quickly under these parameters (**Fig. 1.3 D**). Interestingly (and contrary to intuition), this did not significantly alter the rate at which the infection spread through the tissue. Compare the final frame of **Fig. 1.3 C** (higher tolerance to viral load) to **Fig. 1.3 D** (lower tolerance to viral load). This shows the importance of creating spatiotemporal models of viral replication in tissues, as the balance of competing processes can lead to unexpected dynamics at the tissue, organ, and organism levels.

##### Impact of the cell survival time under high viral loads

We decreased the cell tolerance to viral load further by decreasing the mean cell survival time under high viral loads, which is equivalent to increasing the maximum apoptosis rate $r_{max}$. Following prior analyses[2,3], $1/r_{max}$ is the mean expected survival time as $A \to \infty$. We increased $r_{max}$ from 0.001 $min^{-1}$ (1000 minute expected lifetime at high loads) to 0.01 $min^{-1}$ (100 minute expected lifetime at high viral loads). This drastically accelerated the rate of tissue damage, leaving much more basement membrane (the assumed surface under the epithelial monolayer) exposed (**Fig. 1.3 E**). In a later version of this model framework, we would expect this to lead to earlier onset of fluid leakage, edema, and ultimately adverse respiratory outcomes such as ARDS. Interestingly, this did not significantly increase the rate of spread of the infection. Compare the final frame of **Fig. 1.3 D** (higher tolerance to viral load) to **Fig. 1.3 E** (lower tolerance to viral load).

#### Selected feedback from domain experts and the community

We gathered feedback from the multidisciplinary community, several of whom joined the coalition for future work. We summarize the feedback below.

A virologist noted that more detail on endocytosis, viral uncoating, and synthesis would expose more actionable points in the replication cycle. Preliminary SARS-CoV-2 experiments in her laboratory suggest that the time course (and thus general order of magnitude of rate parameters) is very similar to Venezuelan equine encephalitis virus (VEEV) dynamics measured earlier[4,5]. The exponential progression matches observations: the first cell is infected with one virion and so at first produces virus slowly, but neighboring cells can be infected with multiple virions and thus create virus particles more quickly.

A community member identified typographical errors in the original documentation but verified that that mathematics in the C++ implementation were not affected. He emphasized the importance of implementing RNA decay (as a rate limiting step in virus replication) and the importance of integrating ACE2 receptor trafficking (as a rate limiting step in virus adhesion and endocytosis).

A mathematician noted the potential to simplify the model by removing the diffusing *U, R,* and *P* fields and reported bugs in the initialization (where no cells are initially infected for some domain sizes, due to hard-coding of the initial seeding). Other mathematicians emphasized the importance of varying virion "uptake" with ACE2 receptor availability and hence the need to integrate receptor trafficking.

A mathematical biologist noted prior work on other respiratory viruses will help estimate parameters and build initial immunologic regulation models. Lung pathology and disease severity are closely tied to the immunologic reaction, and prior data and images from influenza will help with calibrating spatial considerations. She expects animal and drug data available for SARS-CoV-2 in the coming months. She noted the importance of distinguishing between mild and severe infections and ARDS. Matching the output to data will be imperative, with one quick possibility to make this match data and distinguish between possibilities is to plot the resulting viral load. She suggested that it would be helpful to show multi focal points of initial infection seeding (possibly of different initial seeding size) that merge together over time, which would match observations of lung histology. Future work will have a better impact if the models uses a true lung tissue geometry with immune cells limiting the peripheral spread. The current model seems more relevant to in vitro growth of a single plaque, which may be scrutinized.

A quantitative systems pharmacologist pointed out the need for clearer scoping and diagrams to clearly lay out the design of each submodel component. We will need procedures to choose future incorporations and changes of scope. He also pointed out the need to understand what happens if you bind up a lot of ACE2 with receptor; there are early insights online[6].

A bioengineer with tissue damage and inflammation expertise noted that the diffusion coefficient of 900 $\mu m^2$/min = 15 $\mu m^2$/s = 1.5e-11 $m^2$/s is not particularly small; prior analyses[7] considered virion diffusion in an lung epithelial monolayer for influenza with D = 3.18e-15$m^2$/s estimating from experimental data. The virions for SARS-CoV-2 could be more mobile though; it is uncertain. There are data[8] about the diffusion coefficient for albumin in tissue being on the order of 10-50 $\mu m^2$/s. She stated that it makes sense for a virion (radius of 25-100 nm) to move more slowly than a protein with radius < 5 nm unless "diffusive transport" in the model is encompassing an active or facilitated transport mode beyond just classic diffusion. She also noted that her laboratory has looked a lot at the renin-angiotensin-system systemically and in kidneys: the kinetics of AngII, ACE, and ACE2 in the lungs would be of interest for connecting the next iteration of the ACE2 receptor model to connect to ARDS. Pfizer may also have relevant related models.

A mathematical biologist with expertise in infectious diseases noted that the model could study immune responses and the impact of mucosal structure in future versions. She suggested quantifying damage or disease metrics. She also noted that ultimately it would be useful to note which parameter estimates might be species-specific and which are not, to be able to switch between experimental and clinical systems, e.g., it is worth recording if current estimates are from human, macaque, etc. She also noted that it may be important to determine if apoptotic cells are replaced or if there is permanent damage (in the model). If the model is run longer, it would be worthwhile to translate the visual sense of damage to a quantitative metric.

An independent team of clinically-focused modelers noted their work on modeling immune expansion in "off screen" lymph nodes and offered to link their model to our immune infiltration functions.

A mathematical biologist with a focus on model and data standards noted the need for clearly specifying each model's assumptions, inputs, and outputs to drive robust parallel development. He noted that it is critical to consider information flow between submodels and revise these data flows as the iterations proceed. He suggested that we state separate execution of submodels as a key design goal to support parallel development. Lastly, he noted that software should be released in conjunction with validation data and methodologies.

#### Core team discussion and priorities for v2

The core team met by virtual conference on April 1, 2020 to discuss the first preprint, model results, and feedback. The core team set as priorities (1) to formalize design specifications for each individual model component and interfaces between components, (2) form teams responsible for each component, (3) focus v2 development on refactoring into this modular format, (3) begin development of the submodels, and (4) begin refine parameter estimates. The clearer specification and organization of submodels was the top priority. As time permits, it was also viewed as important to begin a receptor trafficking model.

The core team agreed to keep working via the dedicated Slack workspace to rapidly coalesce on the submodel teams. Each subteam has a separate channel in the workspace.

### Version 2 (April 1-May 9, 2020)

Version 2 incorporated key v1 feedback, with a focus on introducing a more modular design, improving default model parameters, better initialization options, and a new ACE2 receptor trafficking submodel. This design cycle lasted longer due in part to work spent on subteam organization. As with Version 1, the Version 2 model was developed by the overall leads (Macklin, Heiland, Wang) to refine key model infrastructure for the forming subteams. (See main text section **Three main phases of community-driven development**.)

Version 2 also began work to test the design documents that were first discussed by the core team during the v1 model feedback. The interactive nanoHUB model included new usability refinements, notably an option to animate the model outputs.

#### Model changes

The v2 model was expanded to include the following submodel components (**Fig. 2.1**):

- **T:** tissue (which contains epithelial and other cells)
- **RT:** ACE2 receptor trafficking (including virus endocytosis)
- **V:** viral endocytosis, replication, and exocytosis responses
- **VR:** cell response to viral replication, including cell death and IFN synthesis
- **E:** epithelial cell (incorporates RT, V and VR).

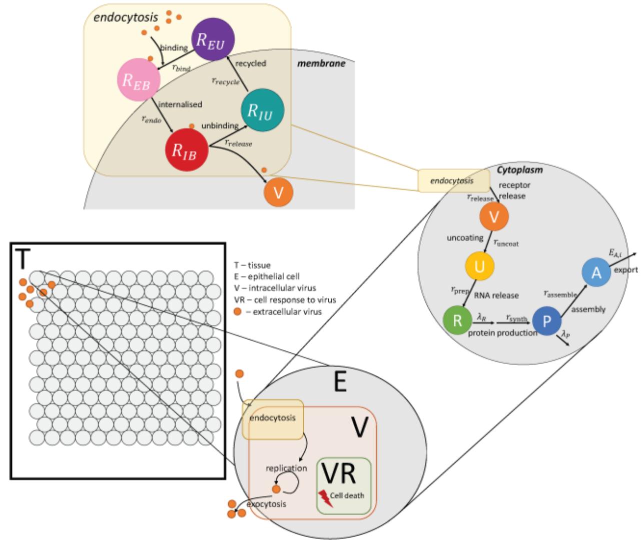


**Fig. 2.1: Version 2 model schematic:** In the overall model structure, a ***tissue component*** **(T)** contains multiple ***epithelial cells*** **(E)**. Viral particles diffuse through this domain and can enter the cells (by adhering as an uptake term) or be exported by cells. Each cell includes an ACE2 ***receptor trafficking*** model (**RT**) where virus can bind an unoccupied surface receptor, which is endocytosed to release its virion and eventually return to the surface. Each cell's ***virus lifecycle*** model **(V)** simulates viral replication kinetics, and its ***viral response*** model **(VR)** uses a dose-response model to control cell apoptotic response to viral load.

Based on community feedback, the default virion diffusion coefficient was reduced by a factor of 10 to 90 $\mu m^2$/min. We may reduce this parameter further based upon oncolytic virus therapy modeling experience by Morgan Craig and Adrianne Jenner.

#### Biological hypotheses

The v2 model was similar to v1 with a simplified set of biological hypotheses:

2.T.1 Virus diffuses in the microenvironment with low diffusion coefficient
2.T.2 Virus adhesion to a cell stops its diffusion (acts as an uptake term)

2.RT.1 Virus adheres to unbound external ACE2 receptor to become external (virus)-bound ACE2 receptor
2.RT.2 Bound external ACE2 receptor is internalized (endocytosed) to become internal bound ACE2 receptor
2.RT.3 Internalized bound ACE2 receptor releases its virion and becomes unbound internalized receptor; the released virus is available for use by the viral lifecycle model **V**

2.RT.4 Internalized unbound ACE2 receptor is returned to the cell surface to become external unbound receptor
2.RT.5 Each receptor can bind to at most one virus particle

2.V.1 Internalized virus (previously released in 2.RT.3) is uncoated
2.V.2 Uncoated virus (viral contents) lead to release of functioning RNA
2.V.3 RNA creates viral protein at a constant rate unless it degrades
2.V.4 Viral protein is transformed to an assembled virus state
2.V.5 Assembled virus is released by the cell (exocytosed)

2.VR.1 As a proxy for viral disruption of the cell, the probability of cell death increases with the total number of assembled virions
1.VR.2 Apoptosed cells lyse and release some or all of their contents

(In the above, X.C.Y denotes prototype X, model component C, biological hypothesis Y, allowing us to easily refer to any individual hypothesis or assumption in discussion and community feedback.) In the next version of this model, we will use the design document protocols for each of these components.

##### Unit tests

The v2 prototype had no changes in qualitative unit tests; once the ACE2 receptor trafficking model works correctly, the model will behave as in v1.

#### Translation to mathematics, rules and model components

##### Cell response (Viral response submodel ***VR***)

There were no changes from the v1 model.

#### Initialization

In v2, we added the option to specify the *multiplicity of infection (MOI):* the ratio of initial virions to number of epithelial cells. These virions are placed randomly in the extracellular space. We use a default MOI = 0.01 to model a fine mist of virions landing on the tissue. Users can also set an option to only infect the centermost cell, which sets $V = 1$ for that cell.

#### Refined parameter estimates

Detailed experimental characterization of ACE2 receptor trafficking in SARS-CoV[9] permits an initial estimation of key model parameters. This experimental work reported that endocytosed receptors were observed in 3 hours post infection, and that 10 hours later (13 hours elapsed time), receptors were observed in vesicles. This estimates the time scale of binding and endocytosis to be on the order of 3 hours, and that virion release occurs on the order of 10 hours. Thus:

$$\frac{1}{r_{\text{bind}} R_{EU}(0)} + \frac{1}{r_{\text{endo}}} \sim 3\text{ hours} \qquad (1)$$

and

$$\frac{1}{r_{\text{release}}} \sim 10\text{ hours}. \qquad (2)$$

Supposing that binding is relatively fast compared to endocytosis, we set $\frac{1}{r_{\text{bind}} R_{EU}(0)} \sim 1\text{ min}$, and so $\frac{1}{r_{\text{endo}}} \sim$

$3 \text{ hours}$. Recycled receptors were observedwithin 14 hours (1 hour after the appearance of endocytosed receptors), so we set $\frac{1}{r_{\text{recycle}}} \sim 1 \text{ hour}$. Assuming there are 1,000 to 10,000 ACE2 receptors per cell, we set the parameters (to order of magnitude) at

$$r_{\text{bind}} = 0.001 \text{ min}^{-1} \quad (3)$$

$$r_{\text{endo}} = 0.01 \text{ min}^{-1} \quad (4)$$

$$r_{\text{release}} = 0.001 \text{ min}^{-1} \quad (5)$$

$$r_{\text{recycle}} = 0.01 \text{ min}^{-1} \quad (6)$$

$$r_{\text{bind}} = 0.001 \text{ min}^{-1} \quad (7)$$

The report observed expression of viral proteins by 18 hours (5 hours after viral release from endocytosed ACE2 receptors). Assuming that $r_{\text{uncoat}} \sim r_{\text{prep}} \sim r_{\text{synth}}$, each parameter has magnitude $0.01 \text{ min}^{-1}$. We similarly set $r_{\text{assemble}} = r_{\text{exo}} = 0.01 \text{ min}^{-1}$ in the v2 model.

#### Other implementation notes

To differentiate between incoming imported and exported virions within the computational implementation, we modeled two diffusing fields (for extracellular concentrations of *V* and *A*). However, the models only require extracellular *V*. At the end of each computational step (advancing by one diffusional time step), we iterate through each voxel and transfer all of the extracellular diffusing *A* to *V*. We also created diffusing fields for uncoated virions, RNA, and viral proteins, although these were removed from later model versions.

By setting the virus uptake rate *U* as noted above, PhysiCell (via BioFVM) automatically removes the correct amount of virions from the extracellular diffusing field and places them in an internalized virus particle variable *n*. By PhysiCell's automated mass conservation:

$$\Delta n = \Delta t \, r_{\text{bind}} n_V R_{EU} = \Delta t \, U_i V_i \rho. \quad (8)$$

If *n* was previously set to zero, then the current value of *n* represents $\Delta n$. By assumption 2.RT.5, $\Delta n$ is equal to the change in the number of external virus-bound receptors (one virion per receptor). Thus, these receptors ($\Delta n$) represent the net increase in bound external receptors. So at each time step, we:

1) Increase $R_{EB}$ by *n*
2) Decrease $R_{EU}$ by *n*
3) Set *n* = 0 (because these virions have been "delivered" to the receptor trafficking model)

#### Software release

The core model associated with the v2 prototype is Version 0.2.1. The nanoHUB app associated with the v2 prototype is Version 2.1. GitHub releases and Zenodo snapshots are given in the Appendix.

The cloud-hosted interactive model can be run at https://nanohub.org/tools/pc4COVID-19.

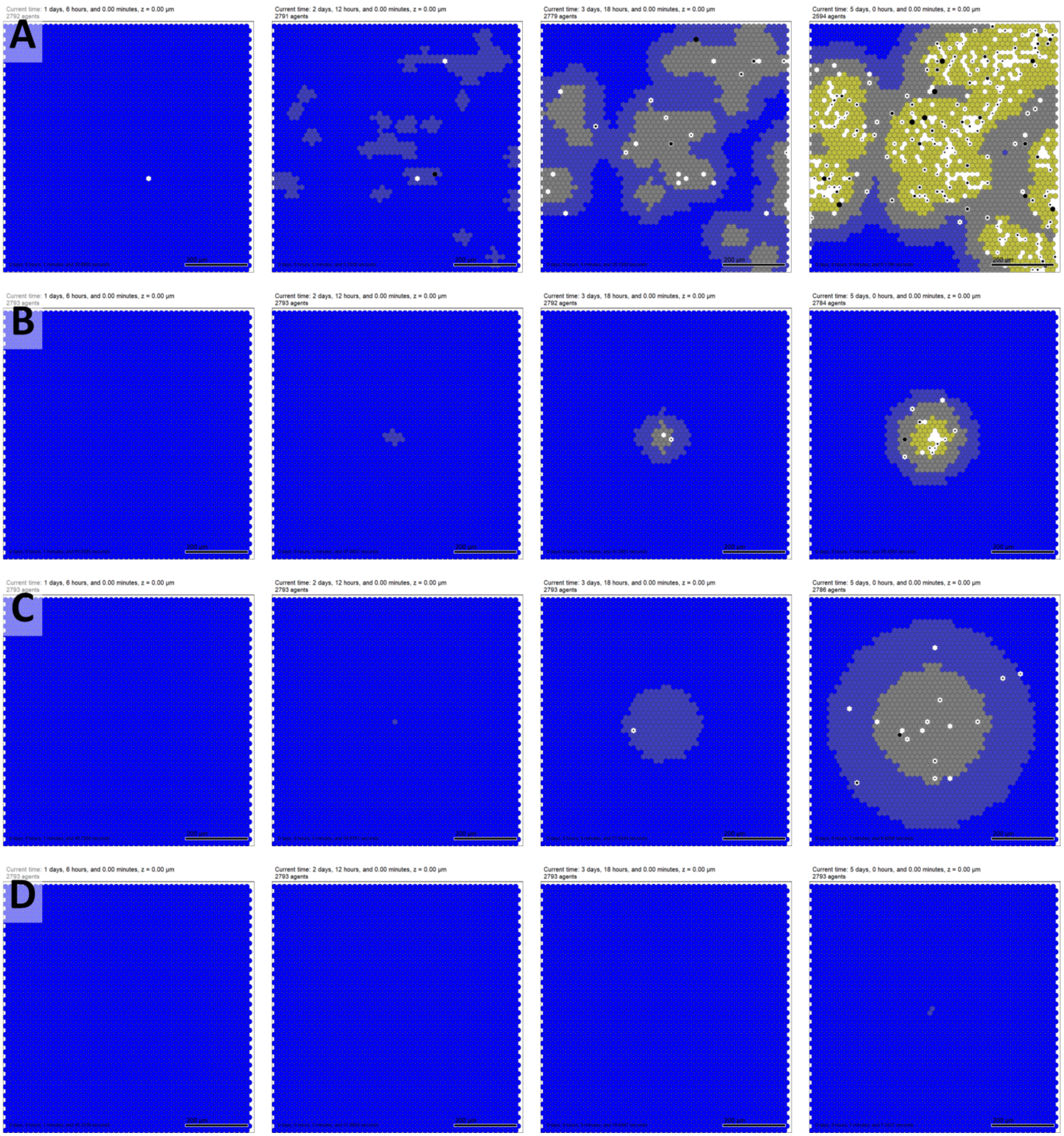


**Fig. 2.2: Version 2 sample model results at 30, 60, 120, and 180 hours.** In all plots, epithelial cells are colored from blue (no assembled virions) to bright yellow (1000 or more virions). Black cells are apoptotic, and white regions show damaged tissues where apoptotic cells have degraded to expose (unmodeled) basement membrane. Bar: 200 $\mu$m. **A.** Simulation time course using the new initialization with an MOI (multiplicity of infection) of 0.01. As virions land randomly on the tissue, they initiate multiple infections that spread and merge. **B.** After infecting a single cell (with the new default parameters), the infected region (plaque) spreads radially as in the v1 model, but at a slower rate. As before, tissue degradation (black apoptotic cells and white cleared tissue) has greatest frequency near the original site of infection. **C.** If the number of ACE2 receptors is cut by a factor of 10, fewer virions infect cells, leading to slower viral replication. However, the reduced rate of virus binding and endocytosis leaves more extracellular viral particles to disperse, leading to a larger spread of the region of infection. **D.** Decreasing instead the rate of virus release from internalized ACE2 receptor drastically slows the viral dynamics.

#### Model behavior: what does the current version teach us?

Except as noted below, all simulation results use the v2 model default parameters, which are supplied in the XML configuration parameter file of the version 0.2.1 core model repository.

In all plots, dark blue cells have 0 assembled virus, pale blue cells have 1-9 assembled virions, grey cells have 10-99 assembled virions, light yellow cells have 100-999 assembled virions, and bright yellow cells contain 1000 or more assembled virions. Black cells are apoptotic, and white spaces show regions devoid of cells (extensive tissue damage).

##### Infection by a single virus versus a dispersion of virions

Compared to the previous method of initially infecting a single cell with a single virion, the v2 model simulation using the new MOI initialization (MOI=0.01) showed viral particles nucleating multiple infections spread as independent plaques that later merge (**Fig. 2.2 A** and **Fig. 2.2 B**). For higher MOIs, some cells can be infected by more than one virion, leading to faster viral replication.

##### Targeting the endocytosis cascade versus targeting ACE2 receptor

As more subcellular mechanisms are added to the model, we can ask *what if* questions about potential pharmacologic interventions[10]. Using the v2 model, we first investigated the impact of reducing the number of ACE2 receptors on each cell by a factor of 10 (e.g., by an intervention that targets ACE2 receptor or reduces its expression). We found that while this reduced the number of viral particles infecting each cell (thus slowing replication in individual cells), it paradoxically *accelerated* the spread of the infected region through the tissue (**Fig. 2.2 C**). This phenomenon can be understood by dimensional analysis: the effective transport length scale $L$ of the virus particle is $L = \sqrt{\frac{D}{U}}$, where $U$ is the uptake rate of the viral particles. In the v2 model, $U$ is proportional to the number of unbound external ACE2 receptors. If this number is reduced, then the length scale increases, leading to a faster dissemination of virus particles, exposing more tissue to virus particles, and ultimately infecting more cells earlier in the disease time course. On the other hand, with slower viral replication in individual cells, tissue damage may be delayed. (**Fig. 2.2 D**).

We similarly investigated whether decreasing the rate of viral release from virus-bound endocytosed receptors by reducing $r_{release}$ by a factor of 10. This drastically impaired the spread of the infection: ACE2 receptors trapped and internalized more viral particles, which then replicated more slowly, thus reducing the severity of the infection.

#### Selected feedback from domain experts within the coalition and the community

The core team reviewed the v2 model and project progress on weekly between April 8, 2020 and May 4, 2020.

The team discussed the potential need for an improved viral replication model. In particular, for low virus counts early in cellular infection, the continuum hypothesis needed for ordinary differential equations may not hold, and non-physical behaviors (e.g., infection by less than a single virus) may prevent the eradication of infections in the model. A discrete modeling approach may be required, although limiting mass transfers (e.g., from $R_{EU}$ to $R_{EB}$) to integer amounts could also help address this issue. The core team also reaffirmed the need to create a simplified immune system model to continue progress.

The core team also identified needed refinements in xml2jupyter, particularly the ability to run additional analytics on simulation outputs and visualize the results in the Jupyter notebook interface.

The core team formed the subteams, identified chief scientists, and organized the first rounds of subteam meetings. The core team also discussed the need to include subteam updates in the weekly core meetings. This was first implemented in the May 4, 2020 call, and the development cycle discussed above reflects these community-driven changes to team management.

We received additional feedback from the community from a postdoctoral fellow at Barcelona Supercomputing

Center (BSC), who noted the number of virion particles should be constrained to integer values. He also suggested a branch of the sub-models may be reimplemented as stochastic differential equation. In addition, the fellow pointed out that BSC is developing COVID-19 molecular disease maps, mainly by curating interactions between viral and cellular proteins from several data sources and domain experts. It may be possible to "translate" these process descriptions to activity flow models for Boolean network simulations in PhysiBoSS[11]. Future collaborations could test the COIVD-19 tissue simulator developed by this coalition in PhyiBoSS.

##### Core team discussion and priorities for v3

The highest priority for v3 is to start transitioning the development of the submodels to the subteams, thus moving the project from Phase 1 to Phase 2. In particular, the team was keen to implement a basic immune model.

### Version 3 (May 10 - July 27, 2020)

Version 3 focused on implementing a realistic representation of the tissue-level immune response to SARS-CoV-2 and transitioning development of the submodels to the subteams. Due to the complexity of the immune system, a significant portion of time was spent developing a realistic minimal model of the immune response.

This also represents the first model release to begin the transition from Phase 1 to Phase 2: the immune team took on primary development of the C++ for their submodel. This development cycle also performed software hardening on the core PhysiCell toolkit to facilitate complex immune behaviors (particularly phagocytosis and CD8$^+$ T cell attacks on infected cells) while improving multithreading safety and cross-platform compatibility. Moreover, we performed a code refactoring to take advantage of new *cell definition* functionality in PhysiCell 1.7.1, which eased the development of the immune model with multiple cell types.

#### Key hypotheses

The overall aim of this submodel is to include features of the immune response to SARS-CoV-2 that are specific to the local tissue environment. The main immune cellular components included at this stage are tissue-resident macrophages, infiltrating neutrophils, and CD8$^+$ T cells, which are recruited as the infection progresses. The general pattern of events that this model encompasses are summarized here. When epithelial cells in the tissue become infected with SARS-CoV-2, they secrete chemokines that cause macrophages to migrate towards them following a chemokine gradient. In addition, the infected cells may die as a result of the infection (see the cell response model ***VR***), and dead cells will release factors that cause macrophages to migrate towards them. Macrophages phagocytose dead cells and remove them from the tissue. When macrophages encounter dead cells, they begin to secrete pro-inflammatory cytokines, and phagocytose any dead cell material they find. The result of pro-inflammatory cytokine secretion is the in-

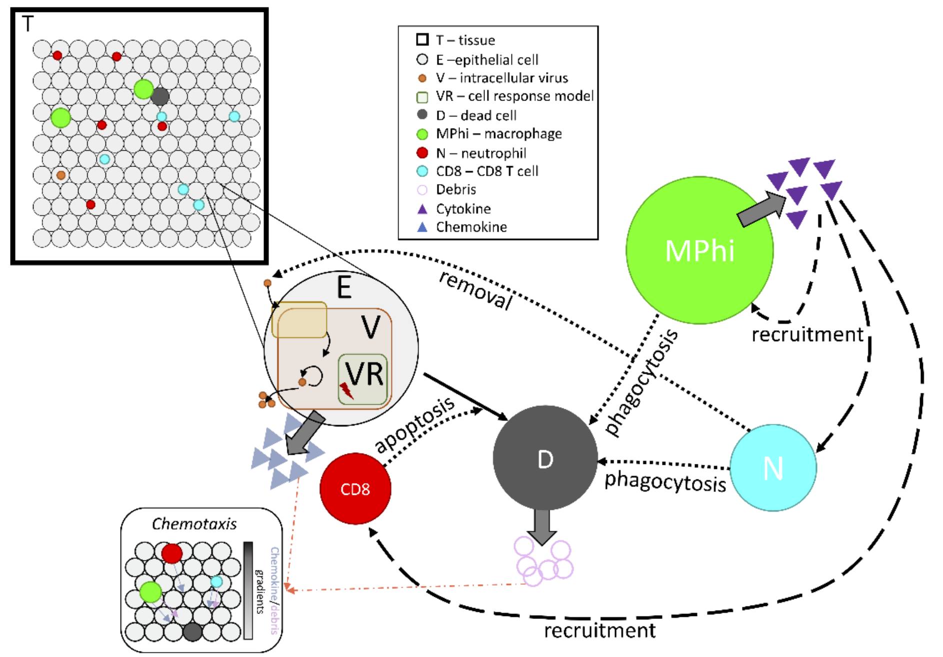


**Fig. 3.1: Immune submodel schematic.** Immune cells (macrophages, neutrophils, and CD8$^+$ T cells) patrol within the *tissue component* (T), containing multiple *epithelial cells* (E). Cells infected by virus secrete chemokine, which attracts immune cells along the chemokine gradient. CD8$^+$ T cells induce apoptosis in infected cells, creating dead cells that are phagocytosed by macrophages and neutrophils that are attracted along debris gradients. Upon activation, macrophages secrete cytokine that recruits other immune cell types.

filtration of neutrophils into the tissue. Neutrophils are short-lived cells and are replenished in the tissue as long as pro-inflammatory cytokines are still being produced. $CD8^+$ T cells, presumed to be specific for SARS-CoV-2, enter the tissue at a later time, and their role is to kill infected cells. $CD8^+$ T cell entry is dependent upon the presence of pro-inflammatory cytokines. Death of an infected cell is more likely after prolonged contact with one or more $CD8^+$ T cells. $CD8^+$ T cells may also interact with macrophages that have phagocytosed dead cells or virus as these macrophages will now be able to present viral antigens to both $CD4^+$ and $CD8^+$ T cells; however antigen presentation is not directly included in this initial version of the model. As described below, simulations were performed using a field of already infected epithelial cells, and the behavior of the immune cells in this tissue appears to follow the established rules.

Owing to the immune response cascade outlined above, we first integrated macrophages, neutrophils, and $CD8^+$ T cells into the SARS-CoV-2 tissue model. The adaptive immune response in a naïve host begins a few days after innate immune action. For this version, we simplify dynamics by modeling only $CD8^+$ T cells, and we do not yet model antigen presentation via dendritic cells or macrophages. Thus, we assume that $CD8^+$ T cells are recruited around day 4 of infection in response to infected cells and pro-inflammatory cytokine production, where infected cells are killed in response to sustained total contact with one or more $CD8^+$ T cells. It is assumed that immune actions only affect infected cells that are past the eclipse phase and are thus generating virus.

Immune cells travel in a biased correlated random walk along chemical gradients[2]. To control for spatial migration, the submodel contains three diffusing chemicals in addition to free virions: pro-inflammatory cytokines secreted by macrophages, $CD8^+$ T cells, and post-eclipse phase infected cells recruit immune cells into tissue from blood or lymph nodes, which could ultimately be modeled in a separate submodel. We assume all immune cells migrate in the tissue toward infected cells along a chemokine gradient, which is assumed to be secreted by infected cells for simplicity. Infected cells and macrophages also secrete IFN-I, which will reduce viral burst size from neighboring infected cells in future versions of the model.

#### Model changes

The v2 model was expanded to include the following submodel components (**Fig. 3.1**):

- **T:** tissue (which contains epithelial and other cells, and diffusible factors)
- **RT:** ACE2 receptor trafficking (including virus endocytosis)
- **V:** viral endocytosis, replication, and exocytosis responses
- **VR:** cell response to viral replication, including cell death and IFN synthesis
- **E:** epithelial cell (includes RT, V and VR).
- **D:** dead cell
- **MPhi:** macrophage
- **N:** neutrophil
- **CD8:** $CD8^+$ T cell

#### Biological hypotheses

The v3 model introduced new assumptions regarding how the infected and dead cells are cleared and how immune cells act in the model (indicated by X.C.Y, where X denotes prototype, C denoted modeling component, and Y denotes a biological hypothesis, for easy reference):

3.T.1 Virus diffuses in the microenvironment with low diffusion coefficient
3.T.2 Virus adhesion to a cell stops its diffusion (acts as an uptake term)
3.T.3 Pro-inflammatory cytokine diffuses in the microenvironment
3.T.4 Pro-inflammatory cytokine is taken up by recruited immune cells
3.T.5 Pro-inflammatory cytokine is eliminated or cleared
3.T.6 Chemokine diffuses in the microenvironment
3.T.7 Chemokine is taken up by immune cells during chemotaxis
3.T.8 Chemokine is eliminated or cleared
3.T.9 Debris diffuses in the microenvironment
3.T.10 Debris is taken up by macrophages and neutrophils during chemotaxis

3.T.11 Debris is eliminated or cleared

3.RT.1 Virus adheres to unbound external ACE2 receptor to become external (virus)-bound ACE2 receptor
3.RT.2 Bound external ACE2 receptor is internalized (endocytosed) to become internal bound ACE2 receptor
3.RT.3 Internalized bound ACE2 receptor releases its virion and becomes unbound internalized receptor. The released virus is available for use by the viral lifecycle model **V**
3.RT.4 Internalized unbound ACE2 receptor is returned to the cell surface to become external unbound receptor
3.RT.5 Each receptor can bind to at most one virus particle.

3.V.1 Internalized virus (previously released in 2.RT.3) is uncoated
3.V.2 Uncoated virus (viral contents) lead to release of functioning RNA
3.V.3 RNA creates viral protein at a constant rate unless it degrades
3.V.4 Viral protein is transformed to an assembled virus state
3.V.5 Assembled virus is released by the cell (exocytosis)

3.VR.1 After infection, cells secrete chemokine
3.VR.2 As a proxy for viral disruption of the cell, the probability of cell death increases with the total number of assembled virions
3.VR.3 Apoptosed cells lyse and release some or all of their contents

3.E.1 Live epithelial cells undergo apoptosis after sufficient cumulative contact time with adhered $CD8^+$ T cells.
3.E.2 Live epithelial cells follow all rules of RT
3.E.3 Live epithelial cells follow all rules of V
3.E.4 Live epithelial cells follow all rules of VR
3.E.5 Dead epithelial cells follow all rules of D.

3.D.1 Dead cells produce debris

3.Mphi.1 Resident (unactivated) and newly recruited macrophages move along debris gradients.
3.MPhi.2 Macrophages phagocytose dead cells
3.Mphi.3 Macrophages break down phagocytosed materials
3.Mphi.4 After phagocytosing dead cells, macrophages activate and secrete pro-inflammatory cytokines
3.Mphi.5 Activated macrophages can decrease migration speed
3.Mphi.6 Activated macrophages have a higher apoptosis rate
3.Mphi.7 Activated macrophages migrate along chemokine and debris gradients
3.Mphi.8 Macrophages are recruited into tissue by pro-inflammatory cytokines.
3.MPhi.9 Macrophages die naturally and become dead cells.

3.N.1 Neutrophils are recruited into the tissue by pro-inflammatory cytokines
3.N.2 Neutrophils die naturally and become dead cells
3.N.3 Neutrophils migrate locally in the tissue along chemokine and debris gradients
3.N.4 Neutrophils phagocytose dead cells and activate
3.N.5 Neutrophils break down phagocytosed materials
3.N.6 Activated neutrophils reduce migration speed
3.N.7 Neutrophils uptake virus

3.CD8.1 $CD8^+$ T cells are recruited into the tissue by pro-inflammatory cytokines
3.CD8.2 $CD8^+$ T cells apoptose naturally and become dead cells

3.CD8.3 CD8+ T cells move locally in the tissue along chemokine gradients
3.CD8.4 CD8+ T cells adhere to infected cells. Cumulated contact time with adhered CD8+ T cells can induce apoptosis (See 3.E.1)

Unit tests

To confirm the dynamics of the immune model qualitatively reproduce the *in-situ* dynamics, we monitored the population numbers of immune cells (macrophages, neutrophils, CD8+ T cells) over time and compared with our biological expectations.

#### Translation to mathematics, rules, and model components

There were no changes to the ACE2 receptor trafficking model ***RT*** or the intracellular viral replication dynamics model ***V.***

#### Initialization

An initial population of $MPhi_0$ macrophages is seeded randomly throughout the tissue.

#### Estimates for immune parameters

The diffusion coefficient for the chemokine, pro-inflammatory cytokine, and debris, $D_{\text{chemokine}}, D_{\text{cytokine}},$ and $D_{debris},$ were set at $555.56\ \mu m^2/min$ which was estimated by Matzavinos *et al* as the diffusion coefficient for monoclonal antibodies[12]. This is equivalent to $8 \times 10^{-3}\ cm^2/day$, which is similar to $1.25 \times 10^{-3}\ cm^2/day$ estimated by Liao *et al.*[13,14]. Decay and secretion rates of the pro-inflammatory cytokine, chemokine, and debris were assumed to be equivalent. Decay rates for the signaling substrates, $\lambda_{chemokine}, \lambda_{cytokine}$ and $\lambda_{debris},$ were all set to $1.02 \times 10^{-2}/min$, which was estimated as the decay rate of IL-6 by Buchwalder *et al.*[15].The secretion rate for each signaling substrate, $S_{chemokine}, S_{cytokine}$ and $S_{debris},$ was $0.8254/\rho^*\ 1/min$, obtained through fitting the secretion rate of infected cells to the production of IL-6 by infected basal epithelial cells measured by Ye *et al.*[16] over 25 hour[16]. The uptake rate of pro-inflammatory cytokine, $U_{cytokine},$ was estimated to be $0.0018\ (pg/ml)^{-1}day^{-1}$ from measurements of the binding rate of IL-6 [17]. The chemokine uptake rate, $U_{chemokine},$ was estimated to be $0.0510\ (pgml)^{-1}day^{-1}$ [18]. The uptake rate of debris, $U_{debris},$ was assumed to be equivalent to that of $U_{chemokine},$ because it acts as a chemoattractant.

Macrophages, neutrophils and CD8+ T cells all have different sizes. Macrophages have an average diameter of $21\ \mu m$ [19], giving a total volume of $4849 \mu m^3$. Neutrophils have an average diameter of $14\ \mu m$ [20], giving a total volume of $1437\ \mu m^3$. When activated, CD8+ T cells have a diameter of 0.7 $9.7\ \mu m$[21], giving a total volume of $478 \mu m^3$. For all immune cells, the volume of nucleus was assumed to be 10% of the cells total volume[22].

The active migration rate of macrophages and CD8+ T cells along the chemokine gradient was $s_{mot,a} = 4 \mu m/min$ based on *in vitro* and *in vivo* measurements of leukocyte chemotaxis rates[23]. We assume these cells have a migration bias of 0.5 (unitless). Neutrophils move faster with stronger bias along the chemokine gradient at $s_{mot} = 19 \mu m/min$, with a bias of 0.91[23]. Once macrophages and neutrophils encounter material to phagocytose, their motility reduces to $s_{mot,p} = 0.4 \mu m/min$ and if a CD8+ T cell connects to an infected cell they are no longer motile, i.e., $s_{mot,p} = 0 \mu m/min$. Cells persist on their given trajectory for $5$ minutes before a new trajectory is chosen. All immune cells undergo apoptosis at different rates, with neutrophils undergoing apoptosis on average after 18.72 hours[24] (i.e. $a_{I,N} = 8.87 \times 10^{-4}$ min$^{-1}$), macrophages on average every 3.3 days[25] (i.e. $a_{I,M\Phi} = 2.083 \times 10^{-4}$ min$^{-1}$) and CD8+ T on average every 2.5 days (i.e. $a_{I,T} = 2.778 \times 10^{-4}$ min$^{-1}$)[26].

Macrophages are well known for their capability in phagocytosing dead cell debris[27]. As such, we set the probability of a macrophage phagocytosing a dead cell in its neighborhood to $p_{phag,M\Phi} = 1$. To account for the fact that neutrophils split their time between phagocytosing dead cells and taking up virus[28], we set the probability that neutrophils phagocytose dead cells as $p_{phag,N} = 0.7$.

For the recruitment of immune cells, parameters were chosen to achieve immunologically reasonable arrival times for the immune cell subsets. Neutrophil and macrophage infiltration into tissue is faster than T cell infiltration, with neutrophils and macrophages arriving within 1 to 2 days after infection[29] and CD8⁺ T cells arriving closer to 4-5 days after infection[30]. The minimum and saturated signal concentrations, $\rho_{min}$ and $\rho_{sat}$, for macrophages and neutrophils were, therefore, assumed to be equivalent and fixed as $\rho_{min} = 0.1\ \mathrm{substrate}/\mu m^3$ and $\rho_{sat} = 0.3\ \mathrm{substrate}/\mu m^3$. Whereas, CD8⁺ T cells had higher signal concentrations for their minimum and saturated recruitment signals, i.e. $\rho_{min} = 0.4\ \mathrm{substrate}/\mu m^3$ and $\rho_{sat} = 0.7\ \mathrm{substrate}/\mu m^3$. The recruitment rate for the different immune types was assumed to be equivalent, i.e. $r_{\mathrm{recruit}} = 4 \times 10^{-9}\ \mathrm{cells/min}/\mu m^3$ and the immune recruitment rate time-step was $\Delta t_{immune} = 10\ \mathrm{min}$.

Direct observations of CD8⁺ T cell-infected cell interactions and quantification of infected cell fate revealed that death required a median of 3.5 distinct CD8⁺ T cell contacts. Killed infected cells have a cumulative median contact time of 50 min and individual contacts between CD8⁺ T cells and infected cells lasts on average 8.5 min[31]. We therefore set $T_{\mathrm{CD8\_contact\_death}} = 50\ \mathrm{min}$.

#### Other implementation notes

This simplified immune model does not yet include many key immune agents, including dendritic cells, natural killer (NK) cells, B cells, antibodies, the complement system, and most cytokines. No anti-inflammatory cytokines are modeled, nor can this model return to homeostasis following potential infection clearance. Dynamics of cytokine binding and unbinding to receptors are also omitted. The model does not yet incorporate known SARS-CoV-2 immune evasion techniques, such as a delayed IFN-I response and lymphopenia (decreased CD8⁺ T cells) from early in infection. In addition, the antigen-presentation from macrophages and subsequent activation process of CD4⁺ and CD8⁺ T cells has been omitted. Many of these important mechanisms are planned for inclusion in future versions. See further discussion in v3 modeling results below.

#### Software release

The core model associated with the v3 prototype is Version 0.2.1. The nanoHUB app associated with the v3 prototype is Version 3.2. GitHub releases and Zenodo snapshots are given in the Appendix.

The cloud-hosted interactive model can be run at https://nanohub.org/tools/pc4COVID-19.

#### Model behavior: what does the current version teach us?

Except as noted below, all simulation results use the v3 model default parameters, which are supplied in the XML configuration parameter file of the version 0.3.2 core model repository.

In all plots, dark blue cells have 0 assembled virus, pale blue cells have 1-9 assembled virions, grey cells have 10-99 assembled virions, light yellow cells have 100-999 assembled virions, and bright yellow cells contain 1000 or more assembled virions. Black cells are apoptotic, and white spaces show regions devoid of cells (extensive tissue damage). Unactivated macrophages are green, activated macrophages are magenta, CD8⁺ T cells are red, and neutrophils are cyan. Apoptotic immune cells are light orange.

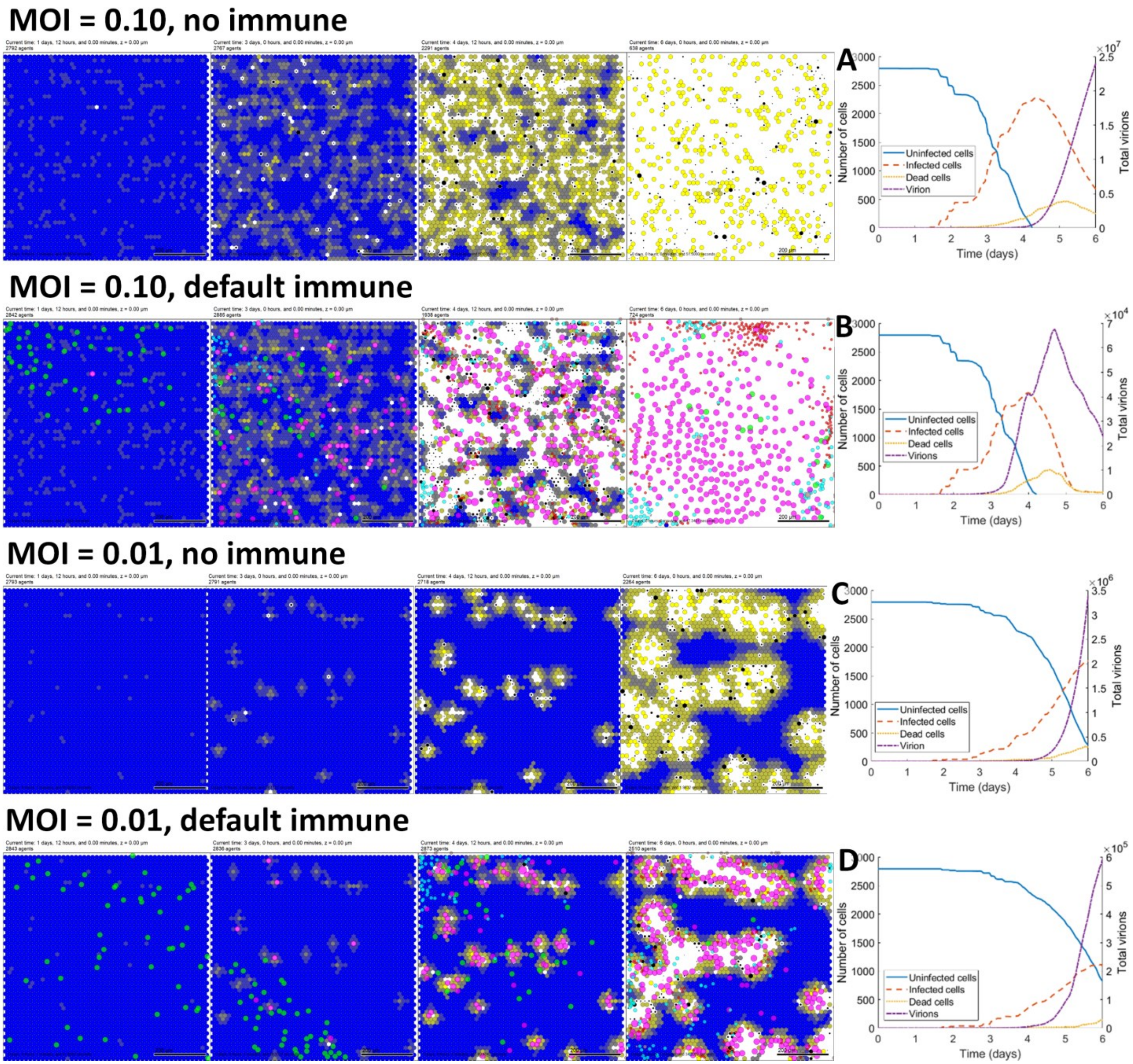


**Fig. 3.2: Version 3 model at 46, 72, 108, and 144 hours (default immune settings).** In all plots, epithelial cells are colored from blue (no assembled virions) to bright yellow (1000 or more virions). Black cells are apoptotic, and white regions show damaged tissues where apoptotic cells have degraded to expose (unmodeled) basement membrane. Green cells are macrophages, magenta cells are activated macrophages, cyan cells are neutrophils, and red cells are $CD8^+$ T cells. Bar: 200 $\mu$m.

**Rows 1-2:** Simulated dynamics starting with an MOI (multiplicity of infection) of 0.10 without an immune response (**Row 1**) and with an immune response (**Row 2**). **Plots A-B** show uninfected (blue), infected (orange), and dead (yellow) cell counts and total extracellular virion (purple) without an immune response (**A**) and with an immune response (**B**). The immune response clears infected and dead cells more quickly and limits the maximum extracellular viral load, but the underlying tissue is completely destroyed.

**Rows 3-4:** Simulated dynamics starting with an MOI (multiplicity of infection) of 0.01 without an immune response (**Row 3**) and with an immune response (**Row 4**). **Plots C-D** show uninfected, infected, and dead cell counts and total extracellular virion without an immune response (**C**) and with an immune response (**D**) (same coloring as A-B). The immune response slows the spread of the infection and increases uninfected cell survival.

#### Impact of adding the immune response (default parameters)

**Figs. 3.2-3.5** demonstrate the results of simulating SARS-Cov-2 infection under different parameter regimes. **Fig. 3.2** simulates the dynamics without and with an immune response for a MOI of 0.10 (top results) and 0.01 (bottom results), using the default immune parameters. When the MOI is 0.10, most of tissue is destroyed, either by the virus (**Fig. 3.2 top row**) or the immune system (**Fig. 3.2 second row**). Reducing the MOI allows some of the tissue to survive the infection in the absence (**Fig. 3.2, third row**) or presence (**Fig. 3.2, fourth row**) of the immune response. All subsequent model results will show a MOI of 0.01 to highlight differences in dynamics to changes in immune parameters. As discussed below, this may not be necessary once the interferon response in infected cells is added to the model.

At high MOI, macrophages are rapidly activated (**Fig. 3.2, first panel in row 2**), and the release of inflammatory cytokines results in the infiltration of CD8$^+$ T cells by day 3 (**Fig. 3.2, second panel in row 2**). The CD8$^+$ T cells kill all infected cells and the tissue is destroyed. At low MOI (0.01), the number of tissue cells surviving is greater (**Fig. 3.2, fourth row, panel 4**). However, macrophage activation is delayed, which further delays the infiltration of CD8$^+$ T cells. At the end of the simulation, on day 6, there is a large number of infected cells (**Fig. 3.2 D**), and the viral titers are still rising. This suggests that, under these default conditions, the lower MOI simply delays the

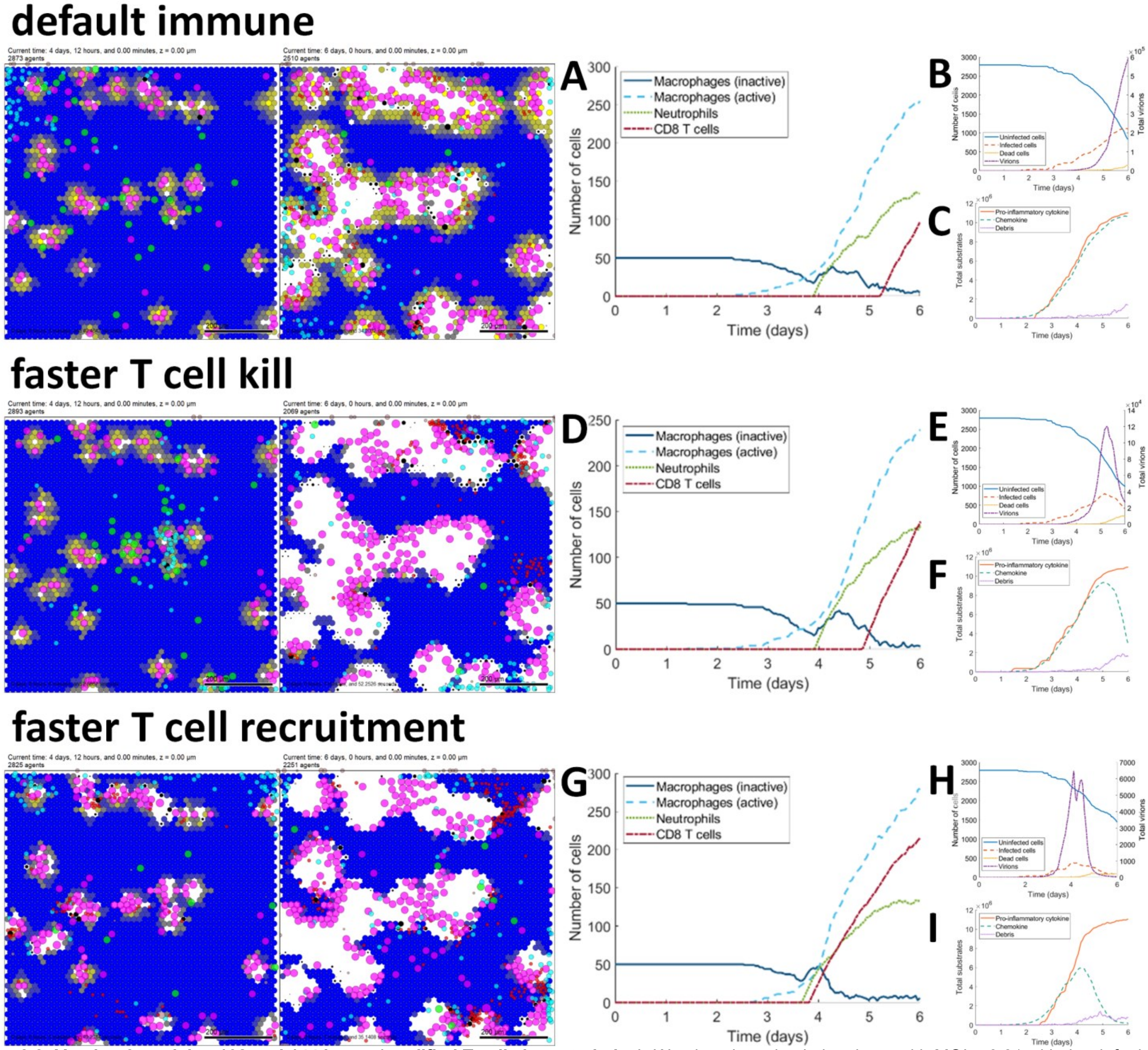


**Fig. 3.3: Version 3 model at 108 and 144 hours (modified T cell characteristics).** We show late simulation times with MOI = 0.01 with the default immune parameters (**top row**), faster T cell killing (**middle row**), and faster T cell recruitment (**bottom row**). **A, D, and G** plot immune cell dynamics, **B, E, and H** plot infected cell and virion dynamics, and **C, F, and I** plot changes in signaling factors for each simulated scenario. Faster T cell killing and faster T cell recruitment both slow infection dynamics and (nearly) control the infection by 6 days. Cells are colored as in Fig. 3.2.

infection; the tissue is completely destroyed if the simulation is continued beyond 6 days, as seen with the higher MOI of 0.10 (result not shown).

#### Changing T cell parameters

In the initial model v3 simulation results introduced in **Fig. 3.2**, it appeared that the dynamics of CD8$^+$ T cell recruitment and activation relative to viral replication might be important. Thus, we varied some of the immune cell parameters to determine whether the survival of the tissue could be improved. **Fig. 3.3 (second row)** shows the results when the rate of CD8$^+$ T cell killing was doubled by reducing the threshold contact time for cell death from 50 min to 25 min. Even macrophage, neutrophil, and CD8$^+$ T cell recruitment were slightly reduced compared to the default parameters (compare **Fig. 3.3 D** to **Fig. 3.3 A**), the increased ability of CD8$^+$ T cells to kill infected cells results in fewer infected cells and the viral titers are falling (**Fig. 3.3 B and E**).

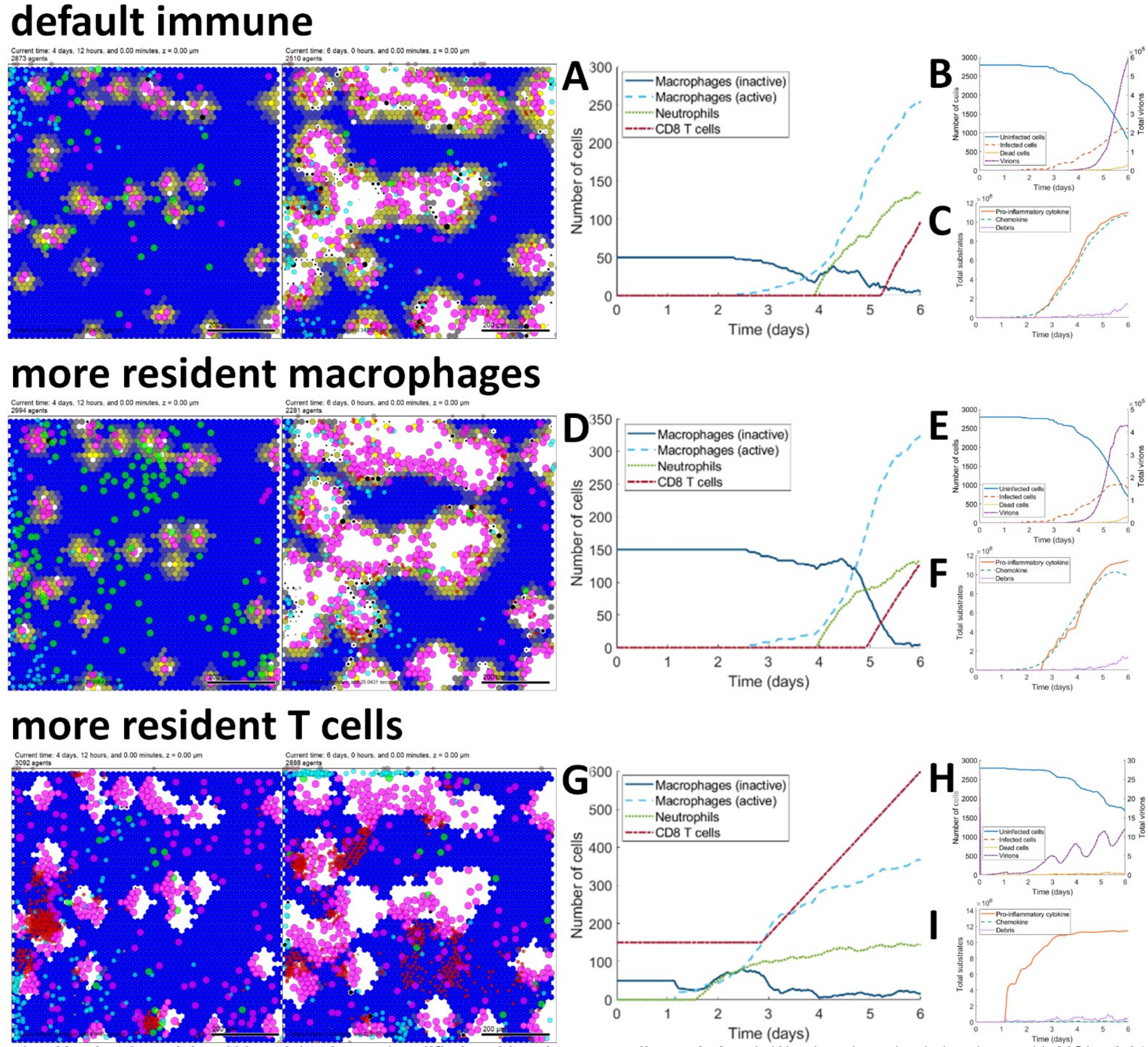


**Fig. 3.4: Version 3 model at 108 and 144 hours (modified resident immune cell populations).** We show late simulation times with MOI = 0.01 with the default immune parameters (**top row**), increased resident macrophages (**middle row**), and addition of resident T cells (**bottom row**). **A, D, and G** plot immune cell dynamics, **B, E, and H** plot infected cell and virion dynamics, and **C, F, and I** plot changes in signaling factors for each simulated scenario. Increasing the resident macrophage population did not appreciably inhibit the infection, while introducing resident T cells reduced viral titers and better controlled the infection. Cells are colored as in Fig. 3.2.

We also increased the recruitment rate of CD8⁺ T cells to the tissue in response to inflammatory cytokines by reducing $\rho_{\min}$ from 0.4 to 0.1, and by reducing $\rho_{\text{sat}}$ from 0.7 to 0.4; see **Fig. 3.3 (third row)**. This resulted in greater sparing of tissue, and the virus was completely cleared under these conditions (**Fig. 3.3 H**).

#### Changing resident immune cell populations

We also investigated whether the outcome would be influenced by the number of macrophages and T cells that might be resident within the tissue at the time of infection. We first increased the initial number of macrophages from 50 to 150 (**Fig. 3.4, second row**). Interestingly, this did not result in an improved outcome over that seen with the default parameters (compare **Fig. 3.4, top row**). CD8⁺ T cell recruitment was enhanced (**Fig. 3.4 D**) but the viral load was continuing to increase at day 6 (**Fig. 3.4 B**), suggesting that increased number of macrophages did not result in more virus control. This could be because in the present model macrophages essentially removed dead cells and do not have a role in killing infected cells. Moreover, because the model's macrophages cannot activate until dead infected cells are present, increasing the number of macrophages cannot trigger a faster immune response. This suggests that after reaching a minimal number of macrophages, adding more resident macrophages has a minimal impact on improving immune response.

In contrast, the presence of resident T cells (**Fig. 3.4, bottom row**) did result in an improved outcome, with more tissue spared and essentially no viral replication (**Fig. 3.4 H**). The final viral count on day 6 was under 15 as T cells were able to kill every infected cell before it could release a significant amount of assembled virions (**Fig. 3.4 H**). This drastically slowed the spread of the infection through the tissue. We also note that the faster T cell killing resulted in earlier accumulation of apoptotic cells, leading to faster activation of macrophages and hence accelerated immune cell recruitment (**Fig. 3.4 G**).

The addition of an interferon response (which could prevent nearby cells from endocytosing these few virions)

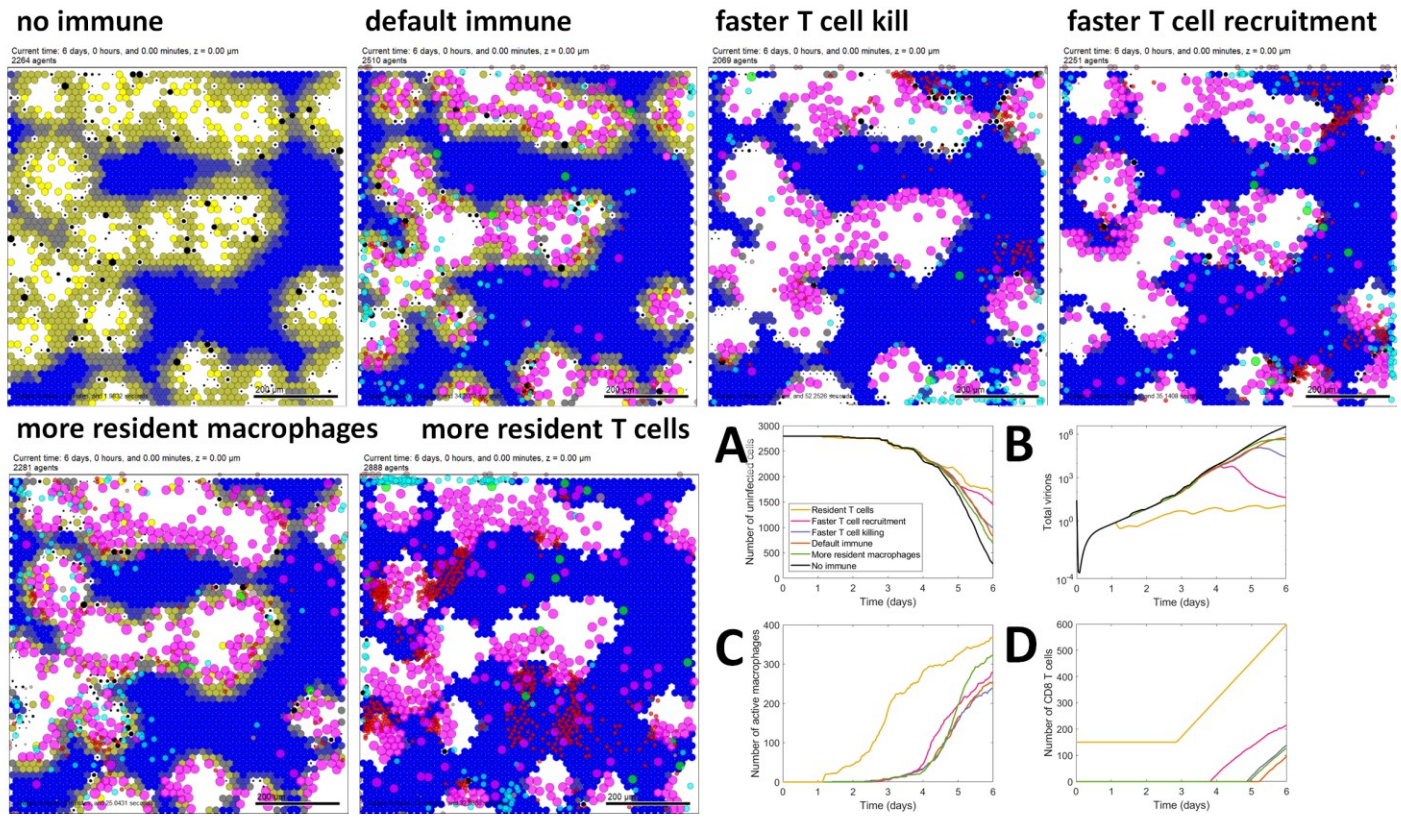


**Fig. 3.5: Comparison of version 3 model runs.** We show late simulation times with MOI = 0.01 with the default immune parameters (**top row**), increased resident macrophages (**middle row**), and addition of resident T cells (**bottom row**). **A, D, and G** plot immune cell dynamics, **B, E, and H** plot infected cell and virion dynamics, and **C, F, and I** plot changes in signaling factors for each simulated scenario. Increasing the resident macrophage population did not appreciably inhibit the infection, while introducing resident T cells reduced viral titers and better controlled the infection. Cells are colored as in Fig. 3.2.

or antibodies (which could directly neutralize the released virions) could potentially completely control the infection in future versions of the model.

##### Further comparison of v3 model results

When we compare these simulations side by side (**Fig. 3.5**) we can see that the two scenarios with the best outcomes are those in which the recruitment rate of T cells is increased and when resident T cells are included. These resulted in the largest amount of surviving uninfected tissue (**Fig. 3.5 A**) as well as viral clearance (**Fig. 3.5 B**). It is interesting to note that the increase in recruitment resulted in T cells arriving in the tissue only one day earlier than the default scenario (**Fig. 3.5 D**), but this had a dramatic effect. Under these conditions, the virus expanded but was cleared within 6 days of infection. This model provides a mechanism to explore how $CD8^+$ T cell recruitment could be enhanced in COVID-19 patients. While the presence of resident T cells prevented the viral infection from propagating there was still a significant amount of tissue loss in the current model.

##### Discussion of v3 model results

The inclusion of the basic tissue immune system to the model has provided some interesting new avenues for research. It is important to note some of the caveats with the existing model. In order to be able to reliably see the beneficial effect of the immune response we had to reduce the viral MOI tenfold. This was because at the higher MOI, either the virus killed the tissue in the absence of the immune response, whereas when the immune response was present it was the immune response that destroyed the tissue (although in this model, the immune cells only destroyed infected cells and did not damage uninfected cells). It is important to note that the present model does not include the cellular interferon response, which would have the effect of slowing viral replication and making more cells resistant to viral infection. When this aspect is included, it is possible that the MOI will not need to be reduced to the same degree.

The results of the simulations so far suggest that controlling the recruitment of $CD8^+$ T cells is a critical parameter leading to viral clearance and reduced tissue destruction. Recent single cell RNA sequencing studies of cells from bronchoalveolar lavage fluid from COVID-19 patients have demonstrated that individuals with moderate disease have increased numbers of $CD8^+$ T cells infiltrating the lungs compared to those with severe disease[32]. This suggests that aspects of the model that can influence this parameter should be explored further. This could include the addition of a more detailed lymph node model in which the kinetics of activation and proliferation of specific $CD8^+$ T cells can be explored.

The presence of resident $CD8^+$ T cells prior to infection effectively prevented viral replication, although at the cost of some tissue damage. It is known that resident memory $CD8^+$ T cells in the lung can provide protection from subsequent infection[33]. It is possible that certain individuals infected with SARS-CoV-2 have been exposed to related coronaviruses and recent studies have observed that up to 40% of unexposed individuals have detectable T cell responses to SARS-CoV-2 proteins[34,35]. Moreover, presence of $CD8^+$ T cells is plausible in uninfected tissues that are adjacent to infected tissues: an initial infection may resemble the default scenario of extensive damage, but the recruited $CD8^+$ T cells could provide protection to nearby tissues from such damage. In the present simulations, it was assumed that all of the resident $CD8^+$ T cells could recognize SARS-CoV-2, but, in reality, only a small fraction of these T cells would be specific for the virus. This can be explored in future simulations.

The fact that increasing the macrophage number did not affect the outcome could reflect the fact that in the present model, these cells have no role in removing infected cells and only remove cells once they have died. Thus, they do not influence the infection beyond secreting inflammatory cytokines that are necessary for the recruitment of $CD8^+$ T cells and neutrophils to the tissue. In subsequent model versions it may be possible to add an interaction between macrophages that have taken up virus and/or dead infected cells with $CD8^+$ T cells. This interaction could further activate macrophages and allow them to kill infected cells before they die.

#### Selected feedback from domain experts within the coalition and the community

During the extensive v3 model development cycle, the modeling coalition met weekly to discuss and refine the model assumptions and record feedback for future work. Several members presented results at virtual seminars and virtual conferences; feedback from audience questions and interactions are also reflected here.

Several members noted that because the ACE2 and viral replication dynamics models use ODEs, it is possible that a cell in a low extracellular viral concentration could endocytose less than one full virion. These cells (with < 1 endocytosed virion) can still synthesize viral proteins and assemble virions in the current ODE model, leading to artificially fast viral propagation in the tissue. Future model versions must compensate for this by only allowing integer numbers of virions to uncoat and replicate. Moreover, for a coated virus such as SARS-CoV-2, we must ensure that only live cells release assembled virions; this is the default model setting in v3 (lysed cells release 0% of their assembled virions by default settings).

It was also widely noted the antiviral effects of interferons should be incorporated to more accurately model the rate of spread of an infection. This is of critical importance in light of recent news reports that interferon beta treatment is emerging as a treatment option[36,37]. Within infected cells, further feedbacks (e.g., on endocytosis) may be needed to prevent further re-infection of those cells. We may also need to model heterogeneity in cells' susceptibility to infection and cytokine production. Cytokine production by infected cells may or may not vary with the amount of virus in those cells.

While the initial immune model in v3 was able to address open questions on the effect of T cells, further work is needed, including explicit cross-talk and feedback between pro- and anti-inflammatory cytokines. Future models should include more neutrophil behaviors, including their own secretion of pro-inflammatory cytokines.

Future immune models should also include activation of antigen-presenting cells (APCs) and T helper cells. Macrophages that have taken up dead infected cells should present antigens to $CD4^+$ and $CD8^+$ T cells. Interactions of macrophages with $CD4^+$ T cells should render them capable of killing infected cells, particularly those with antibodies bound to their surface.

While lymphopenia is a topic of significant clinical interest[38,39], there is currently no mechanism for it in the model. This could be addressed by linking T cell death to the level of inflammatory cytokines, since the degree of lymphopenia has been correlated with levels of IL-6, IL-10 and TNF-$\alpha$[38]. Also, the current model only captures a "cytokine storm" in the sense that as more macrophages are recruited, they also secrete pro-inflammatory cytokines, leading to an accelerated accumulation of cytokines as macrophages accumulate. Future models will need to address this more mechanistically.

In terms of the model development process, we found that there may need to be more flexibility in the length of each development cycle. The transition from Phase 1 to Phase 2 of the project requires substantial training of new developers and creation of software infrastructure. The two-to-three week development cycle noted above is more appropriate to late Phase 2 when all this infrastructure is in place and model changes are more minor from one version to the next.

#### Core team discussions and priorities for v4

In the next development cycle, we plan to introduce type I IFN secretion and its antiviral effects in nearby uninfected cells, particularly reduced receptor endocytosis and viral replication. This will allow us to investigate recent reports on interferon beta

We also plan to introduce a refined, expert-driven model of viral replication to avoid the model artifacts discussed above and to ensure that cells can only replicate virus if they are infected by at least one *whole* virion, that they can only replicate viral proteins if they have at least one full set of viral RNA coated, and they can only assemble virions if they have at least one set of replicated viral proteins. Adding discrete / integer checks on these behaviors may introduce delays analogous to delay differential equations that improve model realism.

We plan to continue refining the immune response submodel as addressed above, with a focus on improving pro- and anti-inflammatory responses and adding missing immune cell types. We plan to link this with a new lymph node model that will more mechanistically regulate T cell expansion, "education" and recruitment.

### Discussion

Within three weeks of the World Health Organization's declaration of a global pandemic of COVID-19[40], community-based prototyping built upon an existing PhysiCell 3D cell-modeling framework to rapidly develop Version 1 of an intracellular and tissue-level model of SARS-CoV-2[2]. A growing coalition of domain experts from across STEM fields are working together to ensure accuracy and utility of this agent-based model of intracellular, extracellular, and multicellular SARS-CoV-2 infection dynamics. Version 1 development underscored the necessity of clearly explaining model components, defining scope, and communicating progress as it occurs for invaluable real-time feedback from collaborators and the broader community. This rapid prototyping already helped in growing the coalition and recruiting complementary expertise; for instance, a team modeling lymph node dynamics and immune infiltration joined during the Version 1 cycle after seeing initial progress.

The version 1 prototype also showed the scientific benefit of rapid prototyping: even a basic coupling between extracellular virion transport, intracellular replication dynamics, and viral response (apoptosis) showed the direct relationship between the extracellular virion transport rate and the spread of infection in a tissue. More importantly, it showed that for viruses that rapidly create and exocytose new virions, release of additional assembled virions at the time of cell death does not significantly speed the spread of infection. Moreover, decreasing the cell tolerance to viral load does not drastically change the rate at which the infection spreads, but it does accelerate the rate of tissue damage and loss, which could potentially trigger edema and ARDS earlier. This suggests that working to slow apoptosis may help preserve tissue integrity and delay adverse severe respiratory responses. That such a simple model could already point to actionable hypotheses for experimental and clinical investigations points to the value of rapid model iteration and investigation, rather than waiting for a "perfect" model that incorporates all processes with mechanistic molecular-scale detail.

Version 2 showed promise of increasing mechanistic details to evaluate potential inhibitors. For example, it was found that that reducing the expression of ACE2 receptors could paradoxically lead to faster spread of the infection across the tissue, although the individual infected cells would replicate virus more slowly. On the other hand, taking advantage of high receptor expression but interfering with viral release from internalized receptors may help slow infectious dynamics. Generally, adding sufficient actionable cell mechanisms to the model framework allows us to ask pharmacologically-driven questions on potential pharmacologic interventions, and how these findings are affected by heterogeneity, stochasticity, and the multiscale interactions in the simulated tissue.

Version 3 allowed our first investigations of immune system responses. We found that T cell behaviors are critical to controlling the spread of an infection through the tissue. In particular, rapid recruitment as well as the presence of "educated" $CD8^+$ T cells prior to infection (e.g., after responding to infection in a nearby tissue) had a significant protective effect, even in the current model that does not explicitly model antibodies. This is consistent with emerging studies that link T cell responses to patients with the best recovery[32,34,35].

As work on future versions progresses, teams will work in parallel on submodels to add, parameterize, and test new model components. It will be important to balance the need for new functionality with the requirement for constrained scope, while also balancing the importance of model validation with timely dissemination of results. Thus, this preprint will be updated with every development cycle to invite feedback and community contributions. Between cycles, the most up-to-date information about this model can be found at http://COVID-19.physicell.org.

### Version 4 (developed July 16, 2020-November 20, 2020)

The most significant change in Version 4 was the integration of a lymphatic system model: a systems of ordinary differential equations now represent arriving dendritic cells that drive expansion of T cell populations. Immune cells traffic between the spatially-resolved tissue model (Versions 1-3) and this new lymphatic compartment.

Version 4 also refined and expanded the version 3 minimal model of the tissue-level immune response to SARS-CoV-2. Dendritic cells and $CD4^+$ T cells have been added to capture antigen presentation dynamics and the interplay between macrophages and T cell signaling. Macrophage activation and phagocytosis mechanisms have also been refined, and we have introduced a model of cell death via pyroptosis. Fibroblast-mediated collagen deposition has been included to account for the fibrosis at the damaged site in response to immune response-induced tissue injury. Epithelial cells' production of Type-I Interferons and Interferon Stimulated Genes' effects on viral replication have been included.

While the version 4 code was released in November 2020, simulation runs, analysis, and scientific documentation continued through April 2021 concurrently with Version 5 code development.

#### Biological hypotheses

The v4 immune submodel introduces dendritic cells and $CD4^+$ T cells as a result of (assumed) antigen presentation in the lymph node as well as the resulting recruitment of $CD8^+$ and $CD4^+$ T cells to tissue. Immune cell recruitment is expanded. Rules regarding macrophage function have been altered to allow for variable engulfment or activation states as well as T cell modulation of macrophage activity. This version also introduces a model of cell death via pyroptosis. Fibroblast recruitment and collagen secretion are also considered for fibrosis. Below, assumptions are indicated by X.C.Y, where X denotes prototype, C denoted modeling component, and Y denotes a biological hypothesis, for easy reference. Assumptions in **bold** have been added or modified from the previous model version.

4.T.1 Virus diffuses in the microenvironment with low diffusion coefficient
4.T.2 Virus adhesion to a cell stops its diffusion (acts as an uptake term)
4.T.3 Pro-inflammatory cytokine diffuses in the microenvironment
4.T.4 Pro-inflammatory cytokine is taken up by recruited immune cells
4.T.5 Pro-inflammatory cytokine is eliminated or cleared
4.T.6 Chemokine diffuses in the microenvironment
4.T.7 Chemokine is taken up by immune cells during chemotaxis
4.T.8 Chemokine is eliminated or cleared
4.T.9 Debris diffuses in the microenvironment
4.T.10 Debris is taken up by macrophages and neutrophils during chemotaxis
4.T.11 Debris is eliminated or cleared
**4.T.12** Anti-inflammatory cytokine is triggered for constant secretion at the site and time that a $CD8^+$ T cell kills an epithelial cell
**4.T.13** Anti-inflammatory cytokine diffuses in the microenvironment
**4.T.14** Anti-inflammatory cytokine is eliminated or cleared
**4.T.15** Collagen is non-diffusive

4.RT.1 Virus adheres to unbound external ACE2 receptor to become external (virus)-bound ACE2 receptor
4.RT.2 Bound external ACE2 receptor is internalized (endocytosed) to become internal bound ACE2 receptor
4.RT.3 Internalized bound ACE2 receptor releases its virion and becomes unbound internalized receptor. The released virus is available for use by the viral lifecycle model **V**
4.RT.4 Internalized unbound ACE2 receptor is returned to the cell surface to become external unbound receptor
4.RT.5 Each receptor can bind to at most one virus particle.

4.V.1 Internalized virus (previously released in 4.RT.3) is uncoated
4.V.2 Uncoated virus (viral contents) lead to release of functioning RNA
4.V.3 RNA creates viral protein at a constant rate unless it degrades
**4.V.4** Viral RNA is replicated at a rate that saturates with the amount of viral RNA
**4.V.5** Viral RNA undergoes constitutive (first order) degradation
4.V.6 Viral protein is transformed to an assembled virus state
4.V.7 Assembled virus is released by the cell (exocytosis)

4.VR.1 After infection, cells secrete chemokine
4.VR.2 As a proxy for viral disruption of the cell, the probability of cell death increases with the total number of assembled virions
4.VR.3 Apoptosed cells lyse and release some or all of their contents
**4.VR.4** Once viral RNA exceeds a particular threshold *(max_apoptosis_half_max),* the cell enters the pyroptosis cascade
**4.VR.5** Once pyropotosis begins, the intracellular cascade is modelled by a system of ODEs monitoring cytokine production and cell volume swelling
**4.VR.6** Cell secretion rate for pro-inflammatory increases to include secretion rate of IL-18
**4.VR.7** Cell secretes IL-1β which causes a bystander effect initiating pyroptosis in neighboring cells
**4.VR.8** Cell lyses (dying and releasing its contents) once its volume has exceed 1.5× the homeostatic volume

4.E.1 Live epithelial cells undergo apoptosis after sufficient cumulative contact time with adhered $CD8^+$ T cells.
4.E.2 Live epithelial cells follow all rules of RT
4.E.3 Live epithelial cells follow all rules of V
4.E.4 Live epithelial cells follow all rules of VR
4.E.5 Dead epithelial cells follow all rules of D.
**4.E.6** Infected epithelial cells secrete pro-inflammatory cytokine
**4.E.7** Antigen presentation in infected cells is a function of intracellular viral protein

4.D.1 Dead cells produce debris

4.MPhi.1 Resident (unactivated) and newly recruited macrophages move along debris gradients.
**4.MPhi.2** Macrophages phagocytose dead cells. Time taken for material phagocytosis is proportional to the size of the debris
4.MPhi.3 Macrophages break down phagocytosed materials
4.MPhi.4 After phagocytosing dead cells, macrophages activate and secrete pro-inflammatory cytokines
4.MPhi.5 Activated macrophages can decrease migration speed
4.MPhi.6 Activated macrophages have a higher apoptosis rate
4.MPhi.7 Activated macrophages migrate along chemokine and debris gradients
4.MPhi.8 Macrophages are recruited into tissue by pro-inflammatory cytokines
**4.MPhi.9** Macrophages can die and become dead cells only if they are in an exhausted state
**4.MPhi.10** Macrophages become exhausted (stop phagocytosing) if internalised debris is above a threshold
**4.MPhi.11** $CD8^+$ T cell contact stops activated macrophage secretion of pro-inflammatory cytokine
**4.MPhi.12** $CD4^+$ T cell contact induces activated macrophage phagocytosis of live infected cells

4.N.1 Neutrophils are recruited into the tissue by pro-inflammatory cytokines
4.N.2 Neutrophils die naturally and become dead cells
4.N.3 Neutrophils migrate locally in the tissue along chemokine and debris gradients
4.N.4 Neutrophils phagocytose dead cells and activate
4.N.5 Neutrophils break down phagocytosed materials
4.N.6 Activated neutrophils reduce migration speed
4.N.7 Neutrophils uptake virus

**4.DC.1** Resident DCs exist in the tissue
**4.DC.2** DCs are activated by infected cells and/or virus
**4.DC.3** Portion of activated DCs leave the tissue to travel to the lymph node
**4.DC.4** DCs chemotaxis up chemokine gradient

**4.DC.5** Activated DCs present antigen to CD8s increasing their proliferation rate and killing efficacy (doubled proliferation rate and attachment rate)

**4.DC.6** Activated DCs also regulate the CD8 level in within a threshold by enhancing CD8 clearance.

4.CD8.1 CD8⁺ T cells are recruited into the tissue by pro-inflammatory cytokines

4.CD8.2 CD8⁺ T cells apoptose naturally and become dead cells

4.CD8.3 CD8⁺ T cells move locally in the tissue along chemokine gradients

4.CD8.4 CD8⁺ T cells adhere to infected cells. Cumulated contact time with adhered CD8⁺ T cells can induce apoptosis (See 4.E.1)

**4.CD8.5** Activated DCs present antigen to CD8 T cells, which increases the CD8 T cell proliferation rate

**4.CD8.6** Activated DCs also regulate the CD8 level in within a threshold by enhancing CD8 clearance.

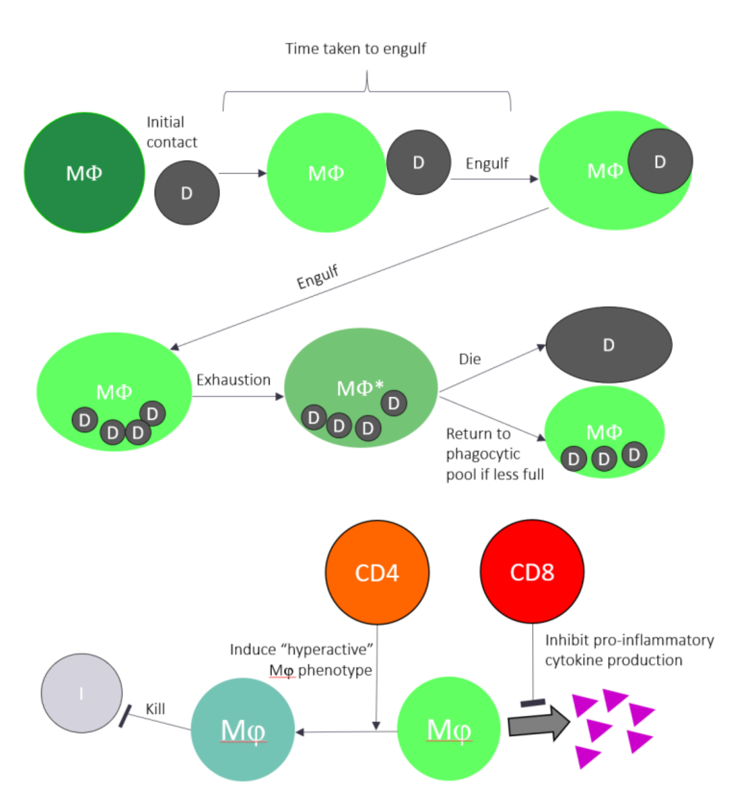


**Fig. 4.2: Macrophage Modulation.** Macrophages engulf dead cell material and grow in size. Too much internalized debris can induce macrophage exhaustion, which causes death or a pause in new phagocytosis. Macrophages that previously engulfed ma-

**4.CD4.1** CD4⁺ T cells are recruited into the tissue by pro-inflammatory cytokines

**4.CD4.2** CD4⁺ T cells apoptose naturally and become dead cells

**4.CD4.3** CD4⁺ T cells move locally in the tissue along chemokine gradients

**4.CD4.4** CD4⁺ T cells are activated in the lymph node by three signals: (1) antigenic presentation by the DCs, (2) direct activation by cytokines secreted by DCs, (3) direct activation by cytokines secreted by CD4⁺ T cells.

**4.CD4.5** CD4⁺ T cells are suppressed directly by cytokines secreted by CD4⁺ T cells.

**4.F.1** Fibroblast cells are recruited into the tissue by anti-inflammatory cytokine

**4.F.2** Fibroblast cells apoptose naturally and become dead cells

**4.F.3** Fibroblast cells move locally in the tissue along up gradients of anti-inflammatory cytokine

**4.F.4** Fibroblast cells deposit collagen continuously

#### Immune sub-model

##### Summary and key hypotheses

The overall aim of this submodel is to include features of the immune response to SARS-CoV-2 that are specific to the local tissue environment. The main immune cellular components included at this stage are tissue-resident macrophages, infiltrating neutrophils, dendritic cells, CD8⁺ and CD4⁺ T cells, all of which are recruited as the infection progresses. The general pattern of events that this model encompasses are summarized here. When epithelial cells in the tissue become infected with SARS-CoV-2, they secrete chemokines that cause macrophages to migrate towards them following a chemokine gradient; they also produce pro-inflammatory cytokines to aid in immune cell recruitment. It is assumed that immune actions only affect infected cells that are past the eclipse phase and are thus productively generating virus. The infected cells may die by apoptosis or pyroptosis following infection or by apoptosis as a result of immune killing (see the cell response model ***VR***), and dead cells will release factors that cause macrophages to migrate towards them.

Macrophages are innate immune cells that phagocytose dead cells to remove them from the tissue. During phagocytosis (which takes some time to complete), macrophages grow in size relative to the material they engulf. If the amount of internalized debris crosses a pre-determined threshold, macrophages become exhausted and stop phagocytosing new material until they either die or reduce in size so they return to the phagocytic pool after losing internalized material. When macrophages phagocytose dead cells, they also begin to secrete pro-inflammatory cytokines that recruit neutrophils and other immune cells into the tissue. Neutrophils are short-lived innate immune cells that kill infected and bystander epithelial cells and are replenished in the tissue as long as pro-inflammatory cytokines are still being produced.

The presence of virus and infected cells in the tissue induces dendritic cells to activate and egress out of tissue to lymph nodes, where they present antigen to induce activation and proliferation of virus-specific $CD4^+$ and $CD8^+$ T cells; a model of immune dynamics in the lymph node is being developed within the consortium (see lymph node model LN) but are not yet explicitly integrated. Some activated dendritic cells remain in tissue and can increase proliferation and cytotoxic function of recruited $CD8^+$ T cells. $CD8^+$ T cells and $CD4^+$ T cells, presumed to be specific for SARS-CoV-2, are activated by antigen presentation in the lymph node and enter the tissue in response to pro-inflammatory cytokines around 4 days after infection. The role of $CD8^+$ T cells is to kill infected cells. Death of an infected cell is more likely after prolonged contact with one or more $CD8^+$ T cells. Both types of T cells additionally interact with macrophages that have phagocytosed dead cells or virus and modulate macrophage activity in two ways: $CD8^+$ T cells inhibit pro-inflammatory cytokine production by macrophages, and contact with $CD4^+$ T cells induces a hyperactive state in macrophages that enables them to phagocytose live infected cells. Other ways in which $CD4^+$ T cells can coordinate the adaptive immune response include: different Th cell subsets, i.e. Th1 Th2, Th17 and Treg, which all may play a role in various stages of the disease by secreting distinct sets of cytokines. The ability of the $CD4^+$ T cells to induce a hyperactive state would most likely be specific to Th1, whereas Th2 and Treg cells may inhibit macrophage activity. The presence of Th17 cells would likely increase the number of neutrophils. There is evidence from recent profiling of clinical COVID-19 cases that cytokine dysregulation plays an important role in determining severity. Thus, it will be important in future model versions to consider varying the functions of $CD4^+$ and $CD8^+$ T cells to reflect these different scenarios.

Tissue damage activates the anti-inflammatory cytokines which then in turn recruit fibroblasts. Fibroblast chemotaxis towards damage site and deposit collagen to repair the tissue. Yet, excess deposition of collagen can lead to fibrosis. See below for further details.

Immune cells travel in a biased correlated random walk along chemical gradients[109]. To control for spatial migration, the immune submodel contains three diffusing chemicals in addition to free virions. Pro-inflammatory cytokines, which are secreted by macrophages and post-eclipse phase infected cells, recruit immune cells into the tissue from circulation or lymph nodes. We assume all immune cells migrate in the tissue toward infected cells along a chemokine gradient. For simplicity, chemokines are assumed to solely be secreted by infected cells. Macrophages also migrate up a debris gradient released by apoptotic cells. Infected cells and macrophages also secrete IFN-I, which will reduce viral burst size from neighboring infected cells in future versions of the model. As described below, simulations have been performed using a field of infected epithelial cells, and the behavior of the immune cells in this tissue appears to follow the established rules.

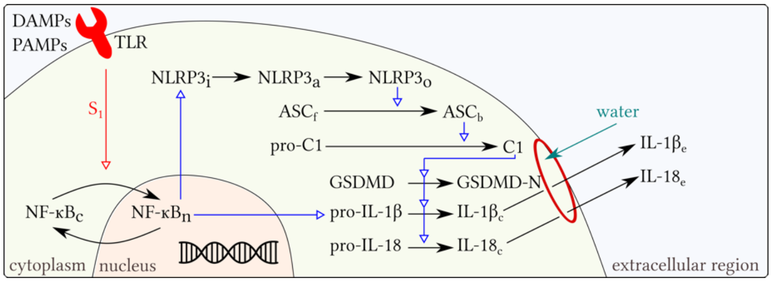


**Fig. 4.3: Pyroptosis pathway.** Viral RNA levels within the cell act as a DAMP/PAMP that initiates the pyroptosis cascade. The intracellular processes result in the secretion of cytokines IL-1$\beta$ and IL-18, as well as cell swelling and rupture.

#### Model changes

The model was expanded to now include the following submodel components overall (**Fig. 4.1 - 4.3**):

- **T:** tissue (which contains epithelial and other

cells, and diffusible factors)

- **RT:** ACE2 receptor trafficking (including virus endocytosis)
- **V:** viral endocytosis, replication, and exocytosis responses
- **VR:** cell response to viral replication, including cell death and IFN synthesis
- **E:** epithelial cell (includes RT, V and VR).
- **D:** dead cell
- **MPhi:** macrophage
- **N:** neutrophil
- **DC**: Dendritic cell
- **CD8:** CD8⁺ T cell
- **CD4**: CD4⁺ T cell
- **F**: Fibroblast

##### Chemotaxis (Chemotaxis model ***DC*** and ***CD4***)

DCs and CD4⁺ T cells in the tissue undergo chemotaxis up the chemokine gradient using the same chemotaxis model for CD8⁺ T cells. They follow the same cell velocity and migration bias defined in Eq. (26) and (27).

##### Phagocytosis dynamics (Phagocytosis of apoptotic cells ***MPhi*** and ***N***)

The time for macrophages to phagocytose material is proportional to the size of the apoptotic material[146,147] they are ingesting. Given an apoptotic cell volume $V_{apop}$ and an internalization rate $r_{internalise}$, the time for a macrophage to internalize an apoptotic cell is $t_{phag} = V_{apop}/r_{internalise}$. During this internalization period, a macrophage is unable to phagocytose anything else.

The phagocytic capacity of an individual macrophage is finite, and experiments have shown that macrophages reach a point of saturation or exhaustion, beyond which their phagocytic activity is impaired[148]. To model this, if the volume of a particular macrophage exceeds the threshold volume $K_{threshold}$, then it enters an exhausted state in which it can no longer phagocytose material. The standard PhysiCell volume model[109] will shrink the cell's volume back below this threshold (see Ghaffarizadeh *et al.*[149] for cell-volume change details) and once the macrophages volume is again below $K_{threshold}$, it will begin phagocytosis again. All other phagocytosis rules described previously still apply.

##### CD8⁺ and CD4⁺ T cell interaction with phagocytosing macrophages (***MPhi***)

Activated macrophages (i.e. macrophages that have phagocytosed at least one apoptotic body) interact with CD4⁺ or CD8⁺ cells. This interaction will only occur if the distance between the activated macrophage and the CD4⁺ or CD8⁺ T cell is less than the sum of their radii multiplied by the factor $\epsilon_{distance}$. A CD4⁺ T cell in the neighborhood of an activated macrophage (i.e. macrophages that have phagocytosed at least one apoptotic body) induces a hyperactive state in that macrophage. Hyperactive macrophages are able to phagocytose infected cells with at least one intracellular viral protein. Phagocytosis of infected cells follows the same phagocytosis model rules as apoptotic bodies. A CD8⁺ T cell in the neighborhood of an activated macrophage stops the macrophage's secretion of pro-inflammatory cytokine, i.e. $s_{chemokine} = 0$.

##### Activation and migration of dendritic cells (***DC***)

Naïve DCs in the tissue are activated if the number of virions in the voxel containing the DC exceeds $v_{DC}$, or if there is an infected cell in their neighborhood for the definition of neighborhood, see Ghaffarizadeh *et al.*[149] with at least 1 viral protein. Once activated, DCs have a probability of departure to the lymph node such that the average departure time is $t_{DC,exit}$ after activation. The amount of DCs entering the lymph node is then scaled by some scaling factor set to the number of epithelial cells present.

##### DC and CD8⁺ T cell interaction (***DC***)

An activated DC will present antigen which will induce CD8⁺ T cell proliferation at rate $\gamma_{CD8\,prolif}$. Additionally, through this interaction, a CD8+ T cell will also increase its infected cell attachment rate $r_{attach*}$. This interaction will only occur if the distance between the DC and CD8+ T cell is less than the sum of their radii multiplied by the factor $\epsilon_{distance}$.

##### Estimates for immune parameters

Initial number of DCs in lung tissue $DC_0$ is approximately in the ratio of one DC to every 100 epithelial tissue cells[150]. DCs have been reported to have a body diameter ranging from 10-15$\mu m$ [151] however inverted microscopy and Wright-Giemsa staining show the extent to which dendrites reach outside the DC body[152]. In the 3D agent-based cellular model of lymph node DCs and T cells, Gong *et al.*[153] modelled DCs as 4 times the size of T cells. A such, we assign DCs a diameter of 15$\mu m$ (3.7 fold larger volume than T cells). We assumed DC migration speed in the tissue was 2-3$\mu m/min$ [154] and persistence time equivalent to CD8s (details in V3). We assume that DCs become activated if there are at least $\nu_{DC} = 10$ virions in their voxel.

In Marino *et al.'s*[155] model for tuberculosis, they estimated that it takes an activated DC between 1hr and 24hrs to exit the lung through lymphatics. In line with this estimate, we model the exit time of activated DCs by assigning a random exit time $t_{DC,exit}$ hours drawn from a uniform distribution $U(1,24)$.

The macrophage internalization rate of dead cell debris ($r_{internalise}$) is set to be 1$\mu m^3/s$ [146,147]. The macrophage exhaustion volume is arbitrarily chosen to be $K_{threshold} = 6500$, which is a 1.34 fold increase on homeostatic volume. The average diameter of a CD4+ T cell from Cyto-Trol is about 4.8$\mu m$ [156], significantly smaller than the CD8+ T cell diameter of 9.7$\mu m$. Similar to DCs, CD4+ T cell migration speed in the tissue was equivalent to that of CD8+ cells. CD8+ T cell interaction with an activated (antigen-presenting) DC induces CD8+ T cell proliferation at rate $\gamma_{CD8\,prolif} = 0.00208\,min^{-1}$ .This value was estimated from a mathematical model of CD8+ T cell response during acute lymphocytic choriomeningitis virus[157]. We assume that DC interaction would increase CD8+ T cell to infected cell attachment rate $r_{attach}$ by 50%.

A designated interaction between two immune cells will occur if the Euclidean distance between the cell centres is less than $\epsilon_{distance}(r_1 + r_2)$, where $r_1$ and $r_2$ are the radii of the two immune cells potentially interaction. We chose $\epsilon_{distance} = 1.75$, to allow cells to interact outside a small distance from their boundary.

##### Unit tests

To confirm the dynamics of the immune model qualitatively reproduce the *in-situ* dynamics, we monitored the population numbers of immune cells (macrophages, neutrophils, dendritic cells, CD8+ T cells, and CD4+ T cells) over time and compared with our biological expectations.

#### Viral binding, endocytosis, replication, and exocytosis

##### Translation to mathematics, rules, and model components

There were no changes to the ACE2 receptor trafficking model ***RT***, extracellular transport tissue model ***T,*** dead cell dynamics ***D***, chemotaxis model ***MPhi, N*** and ***CD8,*** neutrophil viral clearance ***N***, and CD8 T cell induction of apoptosis model ***CD8.*** Additional rules or changes to previously described rules are detailed below.

##### Minor changes to recruitment and viral uptake

Previously recruitment of immune cells used to round to the nearest int (see Eq. 29), this has been changed to always round down then do a probability draw to see if an extra cell is recruited. This leads to a smoother transition in the earlier states of recruitment.

$$\alpha = rate \times \Delta time - floor(rate \times \Delta time) \tag{9}$$

if $U(0,1) < \alpha$,

$$number\ of\ new\ cells = floor(rate \times \Delta time) + 1$$

In addition, to attempt to deal with a discrete to continuum transition the rule for low virion uptake by cells was changed to a probability draw when flux values are under one (i.e less than one virion will be recruited in a time-step)

$$dv_t = r_{bind} * V_{cell} * Virion_{external} * Receptor_{unbound} \tag{10}$$

If $0 < dv_t < 1$

$$virion_{nearest\ voxel} = virion_{nearest\ voxel} - V_{voxel}^{-1} \tag{11}$$

$$Receptor_{bound} = Receptor_{bound} + 1 \tag{12}$$

If $dv_t > 1$

$$\alpha = dv_t - floor(dv_t) \tag{13}$$

$$\Delta v_{external} = -floor(dv_t) * V_{voxel}^{-1} \tag{14}$$

$$\Delta Receptor_{bound} = dv_t \tag{15}$$

if $U(0,1) < \alpha$,

$$virion_{nearest\ voxel} = virion_{nearest\ voxel} - V_{voxel}^{-1} \tag{16}$$

$$Receptor_{bound} = Receptor_{bound} + 1 \tag{17}$$

##### Minor changes to viral replication

The model now incorporates viral RNA replication within the host cell. Only the viral RNA ordinary differential equation is directly modified as follows:

$$\frac{dR}{dt} = r_p U + \frac{r_{rep\ max} R}{R + r_{rep\ half}} - \lambda_R R \tag{18}$$

Here, $r_{rep\ max}$ is the maximum replication rate of viral RNA and $r_{rep\ half}$ represents the viral RNA concentration where the viral replication rate is half of $r_{rep\ max}$.

#### Cell response (Viral response submodel ***VR***)

##### IFN response

An early version of the IFN model was added in v4. In this model IFN interferes with protein synthesis reducing the rate of viral protein synthesis.

$$r_{synthesis} = r_{synthesis}(1 - IFN_{voxel} * IFN_{maximum}^{-1} * r_{IFN\ activation}) \tag{19}$$

Where $IFN_{voxel} * IFN_{maximum}^{-1}$ is bounded to 1.

IFN secretion is controlled through RNA detection and paracrine signals.

$$\Delta IFN_{voxel} = \frac{RNA - RNA_{detect}}{RNA_{maximum} - RNA_{detect}} * r_{IFN\ infection\ secretion} + \frac{IFN_{voxel}}{IFN_{maximum}} * r_{maxparacrine} \tag{20}$$

Where the fractions are bounded to 1.

#### Pyroptosis

Once the viral RNA levels within a cell exceed the threshold (R$\geq 200$), or IL-1β levels in the microenvironment reach the threshold (Cytokine$\geq 100$), the cell can undergo pyroptosis, a form of inflammatory cell death[158]. The pyroptosis cascade within each cell is modelled via a system of ODEs capturing the key components of the pathway.

Many aspects of the pathway are dependent on whether the inflammasome base is still forming ($F_{ib} = 1$) or whether it has formed ($F_{ib} = 0$).

This then initiates the translocation of NF-κB into the nucleus at the rate $k_{nfkb_{ctn}}$. The NF-κB can translocate back to the cytoplasm at the rate $k_{nfkb_{ntc}}$. Therefore, we describe the evolution of nuclear NF-κB, $NFkB_n$, through the equation:

$$\frac{dNFkB_n}{dt} = F_{ib} k_{nfkb_{ctn}} (1 - NFkB_n) - k_{nfkb_{ntc}} NFkB_n \tag{21}$$

Nuclear NF-κB then regulates the transcription of inactive NLRP3 protein, we assume this transcription follows a standard hill function form with a transcription coefficient of $a_{nlrp3}$. This inactive NLRP3 can then become activated at the rate $k_{nlrp3_{ita}}$. We additionally assume some natural decay of the NLRP3 at the rate $d_{nlrp3}$. We therefore describe the evolution of inactive NLRP3, $NLRP3_i$, through the equation:

$$\frac{dNLRP3_i}{dt} = a_{nlrp3} \frac{NFkB_n^{\gamma_{nfkb}}}{H_{nfkb}^{\gamma_{nfkb}} + NFkB_n^{\gamma_{nfkb}}} - k_{nlrp3_{ita}} NLRP3_i - d_{nlrp3} NLRP3_i. \tag{22}$$

Once activated NLRP3 can oligomerize to form the inflammasome base, at the rate $k_{nlrp3_{atb}}$. This process continues until enough NLRP3 has oligomerised/bound together to form the inflammasome base when $F_{ib}$ switches to zero. We describe the evolution of active NLRP3, $NLRP3_a$, through:

$$\frac{dNLRP3_a}{dt} = k_{nlrp3_{ita}} NLRP3_i - k_{nlrp3_{atb}} F_{ib}\, NLRP3_a - d_{nlrp3} NLRP3_a. \tag{23}$$

The evolution of bound NLRP3, $NLRP3_b$, is described through:

$$\frac{dNLRP3_b}{dt} = k_{nlrp3_{atb}} F_{ib}\, NLRP3_a. \tag{24}$$

Once $NLRP3_b \geq 1$, then the inflammasome base has formed and $F_{ib}$ switches to zero.

Once the inflammasome base has formed, ASC protein is recruited and binds to the inflammasome at the rate $k_{asc_{ftb}}$. The change of bound ASC, $ASC_b$, is described through:

$$\frac{dASC_b}{dt} = k_{asc_{ftb}} (1 - F_{ib}) NLRP3_b (1 - ASC_b). \tag{25}$$

Pro-caspase 1 is then recruited to the inflammasome site, and is cleaved by bound ASC to become caspase 1, at the rate $k_{c1_{ftb}}$. Therefore, caspase 1, $C_1$, evolves through:

$$\frac{dC_1}{dt} = k_{c1_{ftb}} ASC_b (1 - C_1). \tag{26}$$

Caspase 1 has the capacity to cleave gasdermin and pro-interleukins within the cell. We assume that caspase cleavage of these molecules follows a hill function with coefficients $a_{...}$. Therefore, the cleaved N terminal of gasdermin, $GSDMD_n$, evolves through:

$$\frac{dGSDMD_n}{dt} = a_{gsdmd} \frac{C_1^{\gamma_{c1}}}{\left(H_{c1}^{\gamma_{c1}} + C_1^{\gamma_{c1}}\right)} (1 - GSDMD_n). \tag{27}$$

Similarly to NLRP3, we assume that pro-interleukin 1β, $IL1b_p$, is transcribed by NF-κB and decays at the rate $d_{il}$. The pro-form is then cleaved by caspase 1 to form the cytoplasmic form, that is:

$$\frac{dIL1b_p}{dt} = a_{il1b_p} \frac{{NFkB_n}^{\gamma_{nfkb}}}{H_{nfkb}^{\gamma_{nfkb}} + {NFkB_n}^{\gamma_{nfkb}}} - a_{il1b_c} \frac{C_1^{\gamma_{c1}}}{H_{c1}^{\gamma_{c1}} + C_1^{\gamma_{c1}}} IL1b_p - d_{il} IL1b_p. \tag{28}$$

Cytoplasmic interleukin 1β, $IL1b_c$, can also decay in the same manner as the pro-form. Additionally, $IL1b_c$, can transport out of the cell via pores formed by gasdermin on the cell surface, at the rate $k_{il1b_{cte}}$. That is, $IL1b_c$, evolves via:

$$\frac{dIL1b_c}{dt} = a_{il1b_c} \frac{C_1^{\gamma_{c1}}}{H_{c1}^{\gamma_{c1}} + C_1^{\gamma_{c1}}} IL1b_p - k_{il1b_{cte}} GSDMD_n \, IL1b_c - d_{il} IL1b_c. \tag{29}$$

External interleukin 1β levels only depend on transport out of the cell in the ODE model,

$$\frac{dIL1b_e}{dt} = k_{il1b_{cte}} GSDMD_n \, IL1b_c. \tag{30}$$

We only consider the level of cytoplasmic interleukin 18, $IL18_c$, which is cleaved from its pro-form by caspase 1 and transports out of the cell via gasdermin pores, in the same way as interleukin 1β. That is, the evolution can be described by:

$$\frac{dIL18_c}{dt} = a_{il18} \frac{C_1^{\gamma_{c1}}}{\left(H_{c1}^{\gamma_{c1}} + C_1^{\gamma_{c1}}\right)} (1 - IL18_c - IL18_e) - k_{il18_{cte}} GSDMD_n \, IL18_c. \tag{31}$$

External interleukin 18 levels only depend upon transport out of the cell in the ODE model,

$$\frac{dIL18_e}{dt} = k_{il18_{cte}} GSDMD_n \, IL18_c. \tag{32}$$

Finally, once gasdermin pores form on the cell surface external material can enter the cell causing the cell to swell. Therefore, we allow the cell volume, $V$, to increase through the equation:

$$\frac{dV}{dt} = k_{vol_c} GSDMD_n\, V. \qquad (33)$$

Once the cell volume reaches 1.5 times the original value, then the cell bursts and all cellular processes cease.

The cytokines released by the cell, IL-1β and IL-18 are then modelled in the epithelial environment. We allow IL-1β to potentially initiate pyroptosis in nearby epithelial cells, through a bystander effect[158-160]. Whereas, IL-18 acts as a chemoattractant for immune cells, directing them to the local environment[158,159,161-163]. Both cytokines are modelled as a diffusible field.

##### Estimation for pyroptosis parameters

NLRP3 has a half-life of approximately 6hrs, therefore, we choose a decay rate $d_{nlrp3}$=0.002 $\min^{-1}$[164]. Pro-IL-1β has a half-life of approximately 2.5 hrs, therefore we choose a decay rate $d_{il}$=0.004 $\min^{-1}$ [165].

For the remaining parameters, we use data from three experimental studies Bagaev et. al.[166], de Vasconcelos et. al.[167], Martín-Sánchez et. al.[168] to estimate a timeline for the events regarded in our mathematical model.

Bagaev et. al. reported that the nuclear NF-κB concentration peaked at 10 minutes post-activation, after which it decreased to a half-maximal level in a gradual manner over 100 minutes. de Vasconcelos et. al[167] indicate that following pyroptosis, cell volume increased to around 1.5 times the original cell volume, gradually for approximately 13 minutes prior to membrane rupture. Furthermore, their results suggest that the time between pyroptosis beginning and complete membrane rupture is approximately 2 hours. Additionally, we can estimate from this data that the NLRP3 inflammasome base is formed somewhere between 56 to 30 minutes prior to cell rupture. In the model, we use the average value (43 minutes) as a time-point for when the inflammasome starts forming (i.e.when the concentration of $NLRP3_b$ reaches the threshold value).

In Martín-Sánchez et. al.[168] their results highlighted that the release of IL-1β coincided with membrane permeability (pores forming), and eventually all of the pro-IL-1β present at the start of the experiments was cleaved and released from the cell, with approximately 90% released within 120 minutes. Therefore, we fit our parameters to result in a large proportion of cytokines to be released from the cell following pyroptosis.

#### Lymph node model

The DCs, migrated to the lymph node, are responsible for the development of adaptive immune response in a broad spectrum, where T lymphocytes contribute a major area. In parallel with the CD8⁺ cytotoxic T-cells mediated infected cell-killing, the CD4⁺ helper T-cells functions in two different domains. The type I helper T-cells (Th1) deal with the inflammation and cellular immune response, where type II helper T-cells (Th2) provide help to B-cells in antibody production. In the current version of this study, we do not include the B-cell activation and antibody response, but mostly the activation and proliferation of CD8⁺ and CD4⁺ T-cells are focused. The migrated DCs present the antigen to the precursor/naive T-cells while producing the primary inflammatory cytokines[132]. Stimulated by the antigen presenting cells (DCs) and corresponding to the ambient cytokines environment, the naïve T-cells start differentiate into two major Th1 and Th2 effector cells. During this time period, previously activated memory helper T-cells also recirculate and start to proliferate.

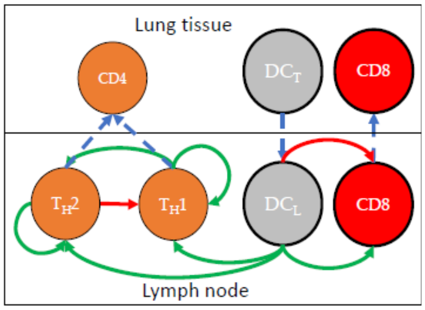


**Fig. 4.1 - supplemental 1: Lymph node model.** Once in the lymph node DCs work to recruit CD4 and CD8 cells to the tissue. Green and red arrows in the figure represent activation and repression. The dotted blue arrows are migration across the tissue barrier.

The mutual regulations of Th1 and Th2 are mediated by the secreted pro- and anti-inflammatory cytokines, although the effects of the cytokines in the current lymph node submodel have been substituted in terms of effective Th1 and Th2 cells. Th1 and Th2 cells have autoregulatory proliferations, where Th2 represses the Th1 cells proliferation. Th1 activation and clearance are regulated indirectly by DCs and directly by the cytokines from both Th1 and Th2, where Th2 clears naturally.

The time course of arrival of the dendritic cells to the LN can be presented as

$$\frac{dD_M}{dt} = k_D D(t - \tau_D) - \delta_{D_M} D_M \tag{34}$$

where, $D$ presents the number of dendritic cells in the tissue and $D_M$ is that migrated into the LN. $k_D$ and $\tau_D$ are the antigen presentation rate by dendritic cells and time taken by the DCs to migrate into the LN. The natural death rate of the $D_M$ is denoted as $\delta_{D_M}$.

In the LN, the proliferation, activation and clearance of the two types of helper T-cells ($T_{H1}$ and $T_{H2}$) and cytotoxic T-cells ($T_C$) are demonstrated in the following ODEs

$$\frac{dT_{H1}}{dt} = \sigma_{T_{H1}} \frac{T_{H1}}{(1 + T_{H2})^2} + \pi_{T_{H1}} \frac{D_M {T_{H1}}^2}{(1 + T_{H2})^2} - \delta_{T_{H1}} \frac{D_M {T_{H1}}^3}{(1 + T_{H2})} - \mu_{T_H} T_{H1} \tag{35}$$

$$\frac{dT_{H2}}{dt} = \sigma_{T_{H2}} \frac{T_{H2}}{(1 + T_{H2})} + \pi_{T_{H2}} \frac{(\rho + T_{H1})}{(1 + T_{H2})} \frac{D_M {T_{H2}}^2}{(1 + T_{H1} + T_{H2})} - \mu_{T_H} T_{H2} \tag{36}$$

$$\frac{dT_C}{dt} = \rho_{T_1} \frac{D_M T_C}{(\rho_{T_1} + D_M)} - \delta_{T_1} \frac{D_M T_C}{\left(\delta_{T_1} + D_M\right)} - \delta_C T_C \tag{37}$$

where, the $\sigma_{T_H}$'s and $\pi_{T_H}$'s are the proliferation and activation rate constants for type 1 and type 2 helper T-cells, respectively. $\delta_{T_{H1}}$represents the DC-mediated deactivation of type 1 helper T-cells, while both have the same natural death rate of $\mu_{T_H}$. The TCs/CD8⁺ T-cells are simultaneously become activated as well as cleared by the DCs by two TC population-dependent rates with rate constants $\rho_{T_1}$, $\rho_{T_2}$and $\delta_{T_1}$, $\delta_{T_2}$. As it currently stands all time delays are set to zero until functionality is built in to the solver.

#### Tissue fibrosis model

The accumulation of fibroblasts in the damaged site and excess deposition of fibrillar collagen leads to fibrosis during tissue regeneration. Sites of alveolar epithelial cell death activate latent anti-inflammatory cytokine (TGF-β), which recruites the fibroblasts[169,170]. We assume that the anti-inflammatory cytokine is activated and secreted continuously at the location of dead epithelial cells killed by CD8⁺ T cells. Fibroblasts chemotax towards the gradient of anti-inflammatory cytokines and deposit collagen. The pathological deposition of collagen can lead to acute or chronic fibrosis. The fibroblast recruitment is mediated by anti-inflammatory cytokine and a correlation for the dependence is adopted from experimental observation[171,172] described by

$$F_g\left(T_\beta\right) = 0.0492T_\beta^3 - 0.9868T_\beta^2 + 6.5408T_\beta + 7.1092 \tag{38}$$

Where, $T_\beta$ is the concentration of the anti-inflammatory cytokines and $F_g\left(T_\beta\right)$ is the recruitment signal for fibroblast depending on the concentration of the anti-inflammatory cytokines. We replace $\rho_{\mathrm{cytokine}}$ in Eq. (29) with $F_g\left(T_\beta\right)$ and set minimum recruitment signal $\rho_{\mathrm{min}}$ to 7.1092 (baseline value) and $\rho_{\mathrm{max}}$ to 12. The value of $\rho_{max}$ was selected based on the experimental observation of fibroblast density during myocardial infarction[169].

##### Estimates for fibrosis parameters

Fibroblast recruitment occurs during the tissue regeneration phase. So, we select a simulation condition where tissue is not completely destroyed after initial infection. We set the condition at faster T cell recruitment, faster T cell kill rate, and MOI = 0.01 (same as v3 Fig. 3.3).

The diffusion coefficient for anti-inflammatory cytokine is set at 555.56 $\mu m^2$/min (same as pro-inflammatory cytokine), and we assume that collagen is non-diffusive. Decay rate for anti-inflammatory cytokine is set to $1.04 \times 10^{-2}$ $min^{-1}$ [173]. We vary the secretion rate of anti-inflammatory cytokine (1 $min^{-1}$ and 15 $min^{-1}$) to observe the change in the concentration of collagen deposition (Fig. 4.4 – supplemental 1).

Fibroblasts have an average diameter of 13 μm [174], and the volume of the fibroblast nucleus is assumed to be 10% of total cell volume. The migration rate of fibroblast along anti-inflammatory cytokine gradient is 1 μm/min [174,175]. Fibroblasts undergo apoptosis at a rate of $8.3 \times 10^{-5}$ min$^{-1}$ [176]. The collagen deposition rate of fibroblast is $0.014\ \mathrm{\mu g/cell/min}$ [172,177].

##### Initialization

An initial population of $MPhi_0$ naïve macrophages and $DC_0$ naïve dendritic cells are seeded randomly throughout the tissue.

#### Other implementation notes

This simplified immune model does not yet include many key immune agents, including natural killer (NK) cells, B cells, antibodies, the complement system, and most cytokines. No anti-inflammatory cytokines are modeled, nor can this model return to homeostasis following potential infection clearance. Dynamics of cytokine binding and unbinding to receptors are also omitted. The model does not yet incorporate known SARS-CoV-2 immune evasion techniques, such as a delayed IFN-I response and lymphopenia (decreased CD8$^+$ T cells) from early in infection. In addition, the antigen presentation and subsequent T cell activation in the lymph node is not yet explicitly modeled. Many of these important mechanisms are planned for inclusion in future versions. See further discussion in the modeling results below.

#### Software release

The core model associated with the v4 prototype is Version 0.4.0. The nanoHUB app associated with the v4 prototype is Version 4.3. GitHub releases and Zenodo snapshots are given in the Appendix.

The cloud-hosted interactive model can be run at https://nanohub.org/tools/pc4COVID-19.

#### Model behavior: what does the current version teach us?

Except as noted below, all simulation results use the v4 model default parameters, which are supplied in the XML configuratioz`n parameter file of the version 0.4.0 core model repository.

##### Overview of v4 model results with base parameters

Initial simulations with all the added model features used the base parameters described above, and the results are shown in Figure 4.4. At time point 0 (Fig 4.4A) resident macrophages and DC can be seen. Following the introduction of virus, the evolution of the infection over time is shown in Figure 4.4B-D. The simulations demonstrate good survival of the tissue in the early time points (2.5 days Fig 4.4B), but survival is not maintained at later time points due to CD8 cells arriving *en masse* to kill the remaining infected cells (10 days, Figure 4.4D). The kinetics of collagen deposition (Fig 4.4E) follows the time course of tissue damage and rises rapidly after 8 days. Interferon (IFN) production (Fig 4.4F) occurs very rapidly and is maintained for 8 days, until cell death increases. Fig 4.4G and H show dynamics of the DC and T cell populations in the LN. It can be seen that there is a constant arrival of DC into the LN starting around day 1 and this is maintained throughout the course of the infection (Fig 4.4G). CD8 cells appear to rise exponentially after day 5 (Fig 4.4G) as do Th2 cells (Fig 4.4H), whereas there appears to be very little Th1 response (Fig 4.4H).

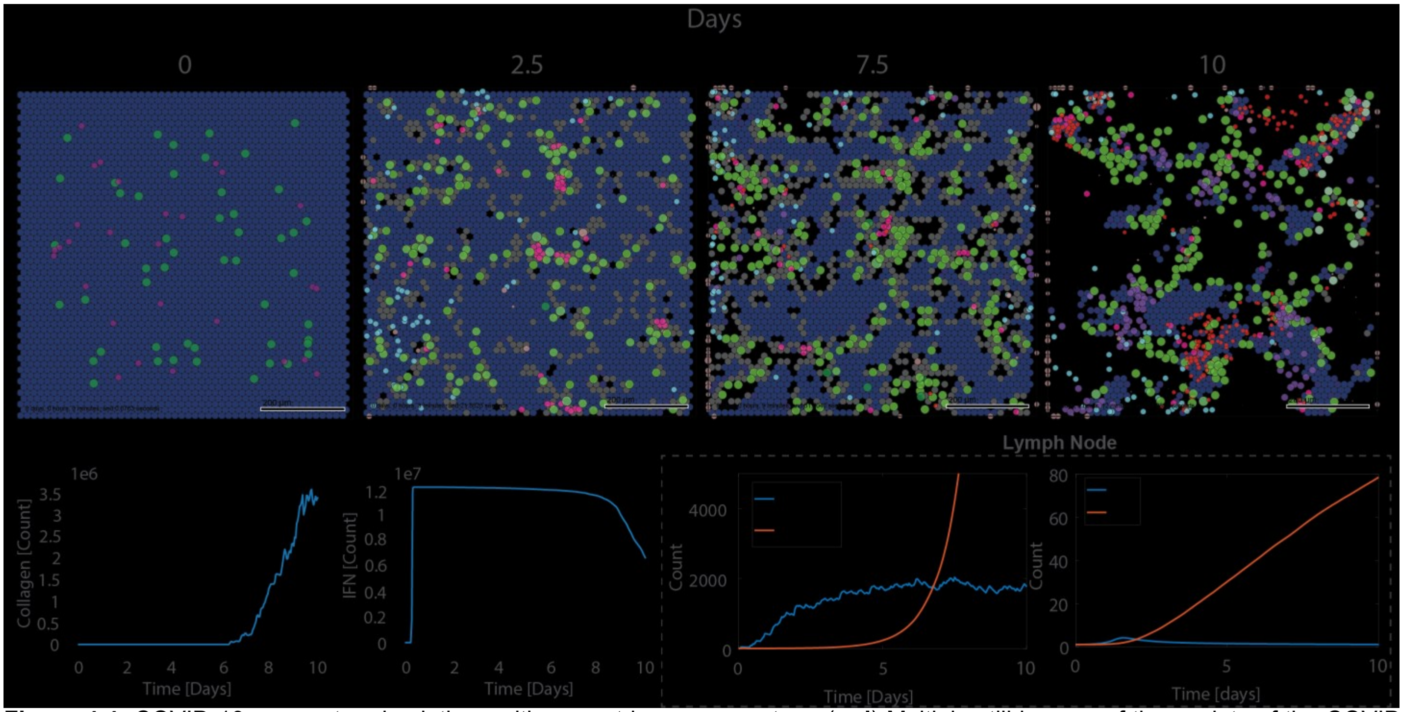


**Figure 4.4:** COVID-19 parameter simulations with current base parameters. (**a-d**) Multiple still images of time points of the COVID simulation at 0(a), 2.5(b), 7.5(c), and 10(d) days. (**e**) Corresponding time plots of Collagen across the simulated 10 days. Low Collagen is present in the system until post 6 days when mass cell death starts to initiate leading to a buildup of collagen. (**f**) Corresponding time plots of interferon (IFN) across 10 days. IFN count rapidly increases in the first day after infection was detected and maintains at a high level through the simulated 10 days. (**g-h**) Lymph node plots showing DC, Tc, Th1, and Th2 counts within the lymph node.

#### Impact of interferon signaling

It was clear from these simulations that the base set of parameters was not sufficient for longer-term tissue survival and recovery, and that there was an over exuberant CD8 T cells response and that the IFN response was not optimal. As a first step the parameters controlling the IFN response were varied. Three parameters were examined: 1) the maximum IFN secretion level ($r_{IFN\ infection\ secretion}$); 2) the relative paracrine secretion, which reflects the ability of uninfected cells to secrete IFN ($r_{maxparacrine}$); and 3) the maximum inhibition of viral protein synthesis by IFN ($r_{IFN\ activation}$).

The maximum IFN secretion rate was examined in the scenario where the paracrine secretion was set to zero (Fig 4.5A-C). In the absence of the paracrine effect and with the default IFN secretion parameter all cells are dead by Day 6 (Fig 4.5A). Increasing the secretion rate 5-fold has little effect (Fig 4.5B) but a 10-fold increase results in better survival (Fig 4.5C). Addition of the paracrine effect to the base IFN secretion rate (Fig 4.5E) significantly improves survival. This is marginally improved if the paracrine effect is increased 10-fold (Fig 4.5F), whereas a 10-fold reduction of the base paracrine effect results in a loss of all tissue (Fig 4.5D). Varying the IFN-induced inhibition of viral protein synthesis also has a dramatic effect on tissue survival (Fig 4.5G-I). The best survival is seen when IFN can inhibit 100% of protein synthesis (Fig 4.5I). Reduction to 80% reduces survival somewhat (Fig 4.5 H), and no tissue survives if this value is reduced to 60% (Fig 4.5G).

The impact of varying the protein synthesis by IFN on cell survival and viral dynamics was also examined (Fig 4.6A-F). From these plots the inhibition of protein synthesis by IFN has to be at least 80% in order to have any significant impact on viral production or cell survival (Fig 4.6A-F). High IFN action sees near 0, or even negative concentrations of extracellular virion (a result of our uptake assumption of only full virions). Rolling average of cellular infectivity shows the cells in the current model do not recover (Fig 4.6C-F), the system seems to only recover after mass cell death.

#### Further exploration dendritic cell and CD8 T cell dynamics

The parameters controlling the DC and CD8 T cell dynamics were also examined with the aim of identifying factors that contribute to the massive influx of CD8 T cells late in the infection. These included: 1) the initial number of DCs in the tissue; 2) the rate of recruitment of DCs to the tissue; 3) the presence of tissue resident CD8 T cells prior to infection. Changing the number of DCs present in the tissue from 3 – 50 (Fig 4.7A-C) had little impact on the infection. As seen in Fig 4.7D, the number of DCs in the tissue rapidly converged to a constant level, and this did not impact the number of CD8 T cells recruited to the tissue (Fig 4.7E). In contrast, varying the DC recruitment rate did not appear to affect tissue damage (Fig 4.7F-H) but had a major impact on DC numbers (Fig 4.7I) and CD8 T cell expansion (Fig 4.7J).

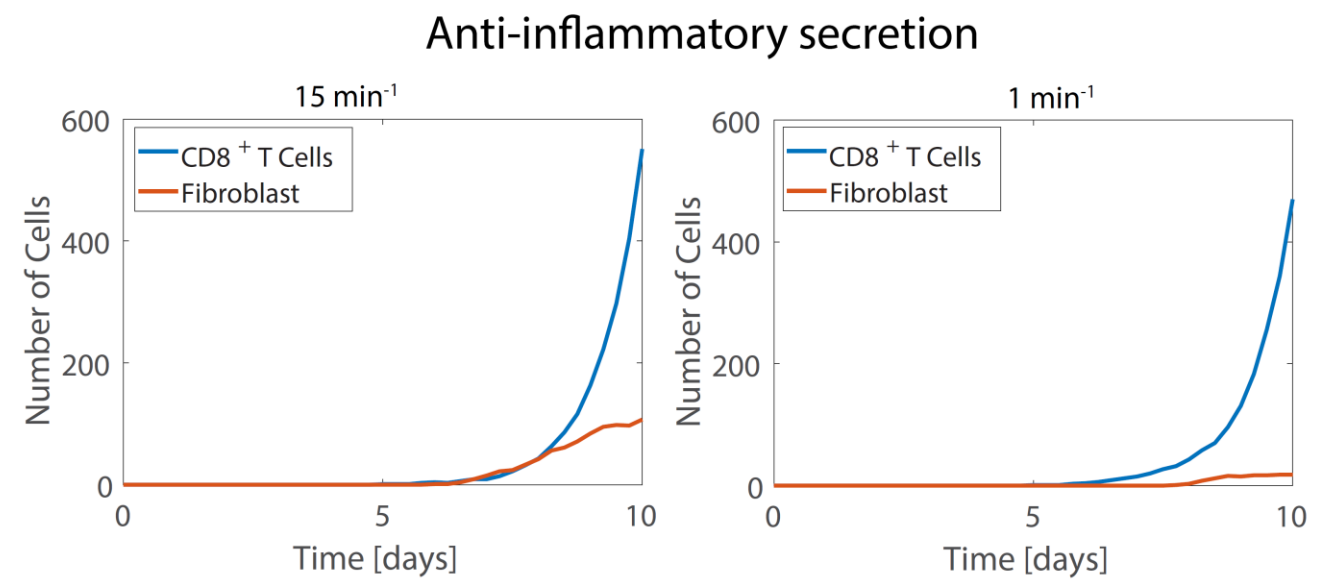


**Fig.4.4 - supplemental 1**: Dynamics of collagen deposition, CD8+ T cells, and fibroblast at anti-inflammatory cytokine secretion rate 15 $min^{-1}$ (Left), and 1 $min^{-1}$ (Right).

The introduction of tissue resident CD8 T cells at the start of the infection (Fig. 4.7K-M) resulted in increased tissue damage, presumably due to the presence of CD8 cells killing infected cells. The number of DCs in the

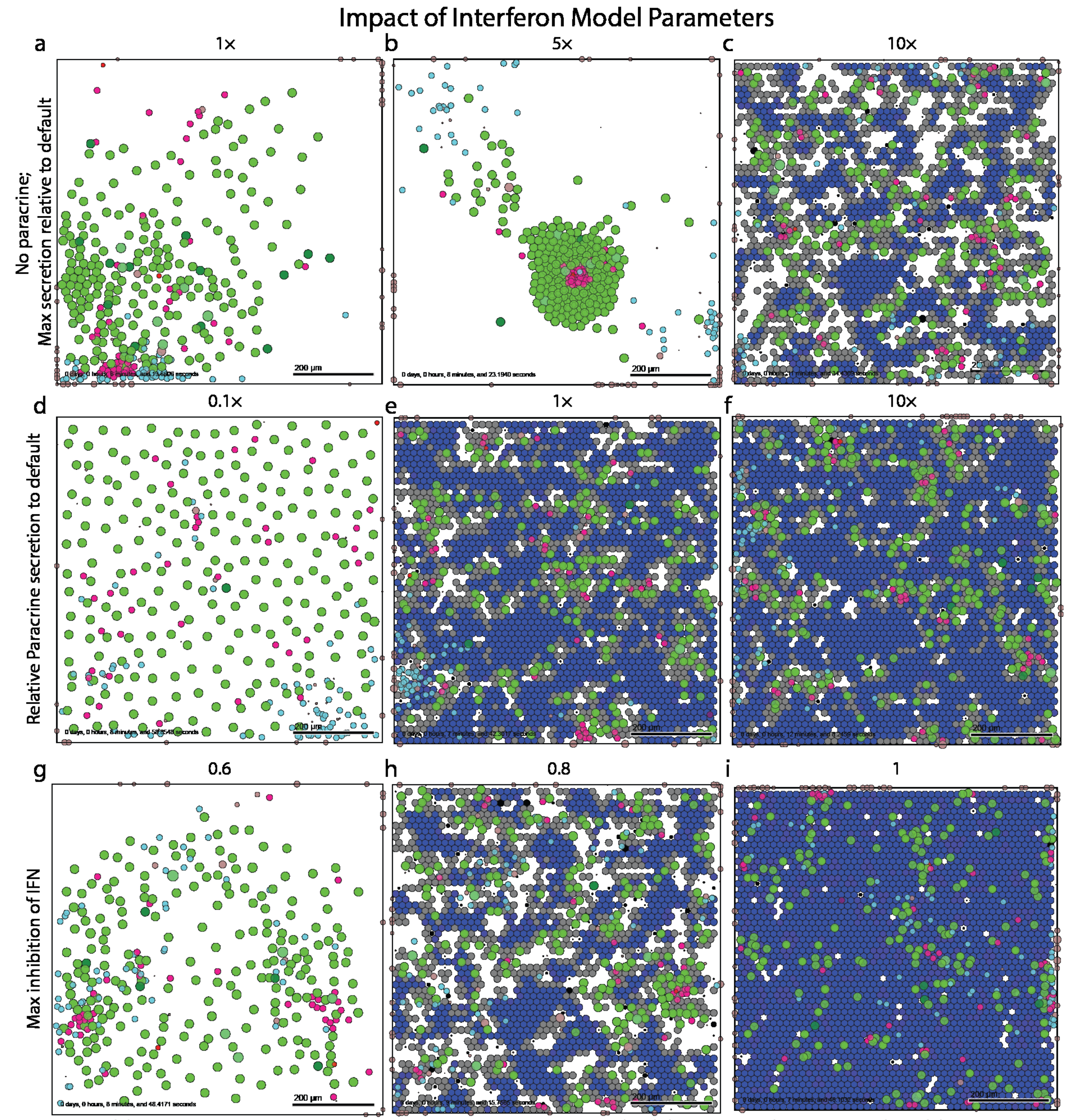


**Figure 4.5:** COVID-19 parameter simulations with current base parameters. (**a-d**) Still images of time points of the COVID simulation at 0(a), 2.5(b), 7.5(c), and 10(d) days. (**e**) Corresponding time plots of Collagen across the simulated 10 days. Low Collagen is present in the system until post 6 days when mass cell death starts to initiate leading to a buildup of collagen. (**f**) Corresponding time plots of interferon (IFN) across 10 days. IFN count rapidly increases in the first day after infection was detected and maintains at a high level through the simulated 10 days. (**g-h**) Lymph node plots showing DC, Tc, Th1, and Th2 counts contained within the lymph node.

tissue was not really affected by preexisting CD8 T cells (Fig 4.7N) except at the later time points, and a similar trend was seen with the CD8 T cell count (Fig 4.7O). It appears that in these scenarios, the infection spreads faster than the CD8 cell killing, which leads to large areas of destroyed tissue. Thus, it appears from these simulations that varying the DC recruitment rate has the most impact on controlling CD8 T cell expansion.

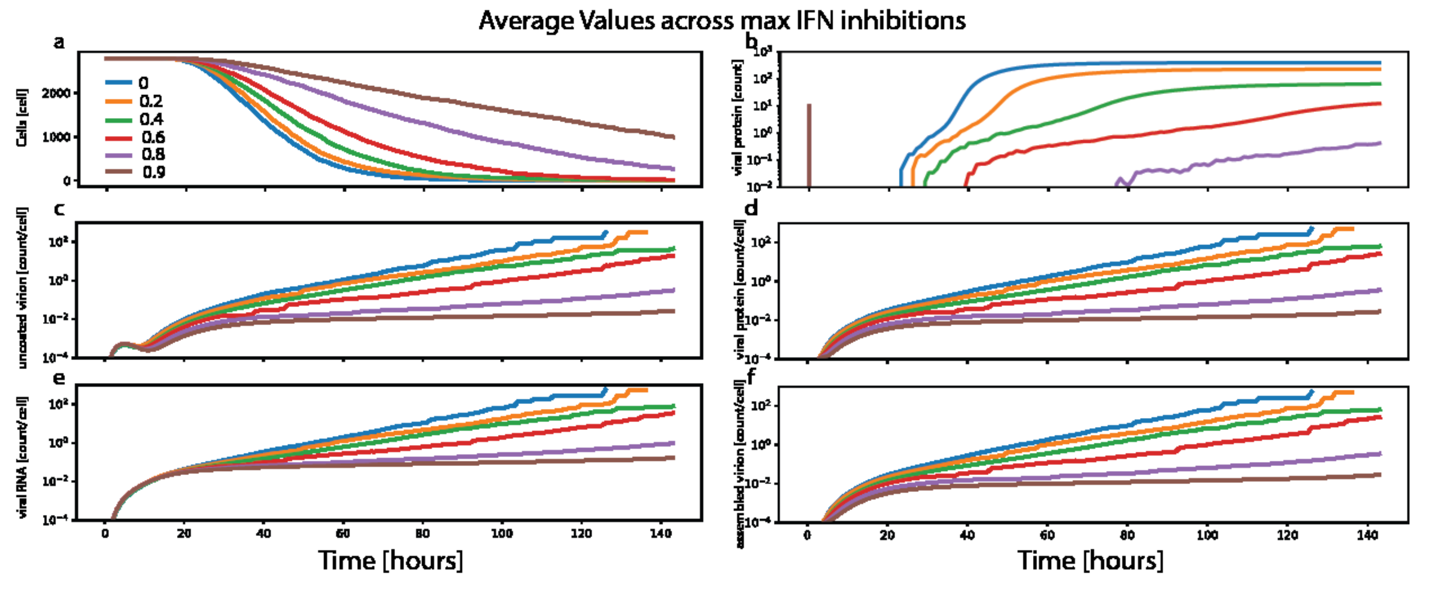


**Figure 4.6:** Time plots of various IFN max inhibition values across 6 days. (a) amount of living epithelial cells (b)external viral protein. (c-f) cellular load plots normalized against the living number of cells to generate a running average value. (c) uncoated virion (d)viral protein (e) viral RNA (f) assembled virion.

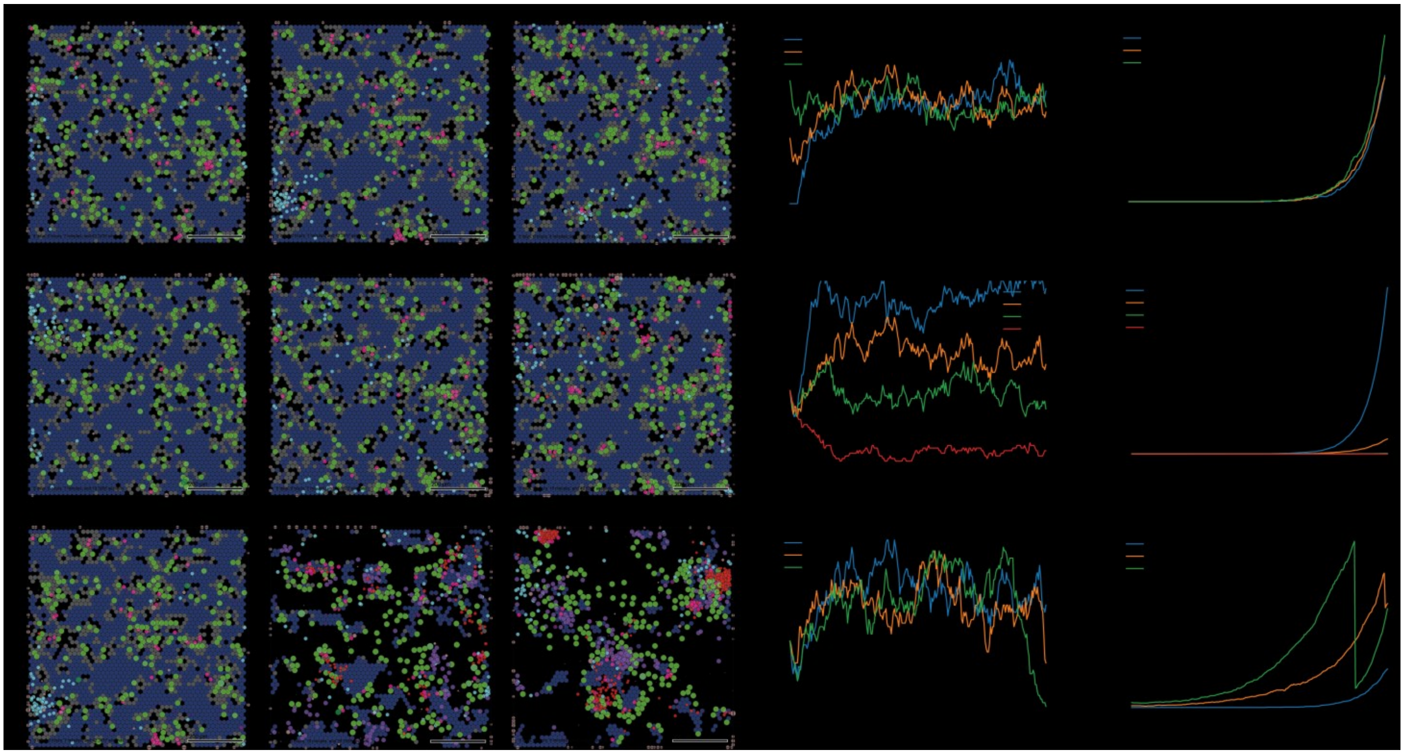

**Figure 4.7:** Multiple variations of immune cell recruitment rates. (a-c) DC initial condition still images at 6 days are shown for 3 (a), 28 (b; base parameter value), and 50 (c) DCs. Plots of DC counts (d) and CD8$^+$ T Cells (e). (f-h) DC recruitment rate still images at 6 days are shown for 0.1x (f), 0.66x (g), and 1.5x (h) the default recruitment rate. Plots of DC counts (i) and CD8$^+$ T Cells (j) at the different recruitment rates. (k-m) CD8$^+$ T cell initial condition still images at 6 days are shown for 0 (k; base parameter value), 10 (l), and 28 (m) CD8$^+$ T cells. Plots of DC counts (n) and CD8$^+$ T cells (o) for the different CD8 T cell initial conditions.

#### Discussion of current model version

This version of the model has included many new components, including more detailed models of viral replication, the introduction of the type 1 IFN response and a LN model that includes DC migration and activation of

CD4 and CD8 T cells. In addition, different macrophage activation states, a pyroptosis death model and the generation of fibrosis in response to tissue damage. The addition of these components has introduced many parameters that creates challenges in calibrating the model.

The inclusion of a more realistic viral replication model has created an infection that spreads very rapidly when compared to previous versions. This has resulted in an inability to clear virus under even the most favorable immune response conditions. Under the present conditions, new infected cells always appear and these cells are killed due to infection or by CD8 T cells and macrophages that are recruited to the tissue *en masse*. There are multiple approaches to address this issue. For one, better slowing viral spread through free parameters (i.e., viral uptake at the current state is underdeveloped which may be causing rapid viral uptake). Even initial viral seeding could be a solution: at present virus is introduced at the start of the simulation randomly across the whole tissue at an MOI of 0.1. This results in widespread virus seeding, which then spreads rapidly before the immune response begins. A more realistic approach may be to seed the virus in a discrete local area, and this will be explored in the next version of this model.

Another concern has been the rapid and exponential expansion of CD8 T cells, which then leads to excessive killing of infected cells. This is driven, in part, by the fact that under the present conditions, DCs continuously migrate to the LN to stimulate CD8 T cells. This could be remedied by tempering the infection as described above. Once the infection is cleared, DCs no longer migrate to the LN and the response is controlled. Data from influenza infections may be used to calibrate this aspect of the model[178].

In addition, some changes to the LN model might be considered. It is known that Th1 cells are necessary for optimal CD8 responses and the present model does not include this requirement. The model could be refined by adding the requirement that Th1 cells are necessary for the proliferation and expansion of CD8 T cells. In this scenario, activated DCs carrying viral antigen migrate to the LN where they activate CD8 T cells and induce the expansion of Th1 and Th2 cells. Th1 cells interact with CD8 T cells to induce their proliferation and expansion. Th1 and Th2 cells are mutually inhibitory[179], but in the present model the inhibition of Th2 cells by Th1 cells is not included, which results in the domination of the Th2 response. The Th2 response is necessary for optimal antibody production and thus it will be important to model this accurately. Recent data suggest that patients with severe COVID-19 disease had robust but delayed IgG anti-spike antibody responses [180]. Thus, the balance and timing between the Th1 and Th2 responses may be critical in determining whether the disease is mild or severe.

#### Priorities for Version 5

After extensive discussion of the Version 4 model results, the coalition set priorities for Version 5 development. First, it was recognized the cell killing by neutrophils can have a “bystander” killing effect: reactive oxygen species (ROSs) created and released by neutrophils and macrophages may kill nearby cells[181-183]. This could have multiple effects on the viral plaques: increased killing could accelerate tissue damage, but killing nearby infected cells could prevent them from successfully replicating and releasing virus. Thus, localized bystander killing could potentially have a protective role as well. Given the importance of antibodies to COVID-19 clinical care (via monoclonal antibody treatments[181], convalescent plasma[184], or the vaccine-driven immunity [185]), the next release will include a model for production, transport, and function of antibodies. The coalition also recognized that as a viral infection is brought under control, negative regulators must initiate an anti-inflammatory response to prevent sustained tissue damage; thus, Version 5 is expected to include anti-inflammatory responses.
